# Identifying the causal allele in the CD40 autoimmune locus enables discovery of context-specific trans-effects in B cells

**DOI:** 10.64898/2026.07.29.741564

**Authors:** Yoshihiko Tomofuji, Zepeng Mu, Hafsa Mire, Yu Zhao, Cassidy Liu, Vidya Jayanthi, Nicholas Sugiarto, Accelerating Medicines Partnership®: RA/SLE Network, Jeffrey A. Sparks, Soumya Raychaudhuri

## Abstract

Thousands of genetic variants are associated with autoimmune diseases, but causal variants, their mechanisms, and the pathogenic context in which they act are elusive. Knowledge of pathogenic contexts may enable effective targeted therapies, instead of broad immunosuppressive approaches. First, to focus on the genetics of immune response, we used surface marker CITE-seq data from 1,055,857 peripheral blood mononuclear cells from 356 individuals. We defined genetic associations to 148 surface proteins across eight cell types. We observed a signal in the CD40 locus, implicated in rheumatoid arthritis (RA) and other autoimmune conditions. RA risk variants increased CD40 protein expression by ∼20% on B cells, but with minimal mRNA effects. Second, we deployed base-resolution genome editing, with CRAFT-seq, capturing genomic DNA sequence at the edited site and multimodal phenotypes at single-cell resolution. We defined a single causal allele, rs1883832, within the Kozak motif. Third, we edited this allele, in primary B cells and conducted CRAFTseq to demonstrate *trans*-effects in >200 genes. These effects were only in the light zone germinal center-like state. Importantly, these *trans*-effects were not seen in population-scale cohorts of unstimulated B cells. This represents a framework to define disease causal alleles, their *cis*– and *trans*-effects. It demonstrates the power of defining causal genetic variation to find *trans*-effects through editing, which cannot easily be found in population studies.

## Introduction

Although hundreds of autoimmune disease loci are known^1^, their causal alleles and molecular effects remain unknown. For example, the *CD40* locus is implicated in multiple autoimmune diseases, including RA^2,3^, multiple sclerosis (MS)^4^, inflammatory bowel disease (IBD)^5^, and Graves’ disease (GD)^6^. However, the *CD40* causal allele, and its *cis-* and *trans*-effects are still unknown. CD40 is expressed on the surface of myeloid and B cells and drives important B-cell functions^7^. Non-selective B-cell depletion using CAR-T cells may have the potential for immune reset to autoimmune disease cure^8–10^. If the precise context that the CD40 allele acts in is defined, a more specific pathogenic B-cell state might be appreciated and targeted.

Immune cell states are often defined by surface proteins that initiate immune responses and state transitions. Surface protein quantitative trait loci (spQTL) may be an important complement to expression QTL (eQTL) to define disease-relevant cell states. Protein QTL studies thus far have relied on serum^11–13^ or bulk tissue^14^ assays, lacking cell-type information. With advances in single-cell profiling, multiplexing >100 surface proteins is now possible^15,16^, enabling mapping comprehensive cell-type-specific spQTLs. However, just like human disease genetic studies, spQTL studies are plagued by linkage disequilibrium (LD), preventing causal variant definition.

Experimental validation is necessary to define causal genetic variants. We and others have used CRISPR genome editing^17–19^ to introduce candidate causal variants into cells. To successfully define the effect of the edited variant, assays must account for bystander editing, low editing efficiency, and non-specific pool effects^20–23^. To address these issues, we developed CRAFTseq^23^, a single-cell method that sequences the genomic DNA of the edited region, while jointly quantifying the transcriptome and surface proteins. This enables comparison of cells with different editing outcomes derived from the same genetic background and culture environment; it facilitates accurate definition of causal SNP *cis-* effects with a modest number of cells^23^.

To understand the disease variant effect, it is as important to understand *trans*-effects as *cis-*effects. *Trans*-effects may induce changes in cellular function that trigger disease susceptibility^24^. However, *trans*-effect mapping efforts have been largely underpowered and not reproducible^25–29^. Since disease alleles may exert context-specific *trans*-effects^30–32^, these effects may be particularly challenging to find in population studies. Experimental editing strategies introducing the causal variant on an isogenic background within a specific context may accurately reveal context-specific *trans*-effects.

We performed cell-type-resolved spQTL mapping (**Figure 1A**). We observed that the CD40 spQTL in B cells overlaps with multiple autoimmune disease GWAS loci and performed CRAFTseq. We identified rs1883832 as a causal spQTL variant. With knowledge of the causal variant, we evaluated its effect in cultured primary B cells and identified *trans*-effects on >200 genes that were context-specific. This reflects a roadmap to integrate genome editing and single-cell QTL mapping to definitively map *cis-* and *trans-* causal mechanism of a disease risk locus.

**Fig. 1.**
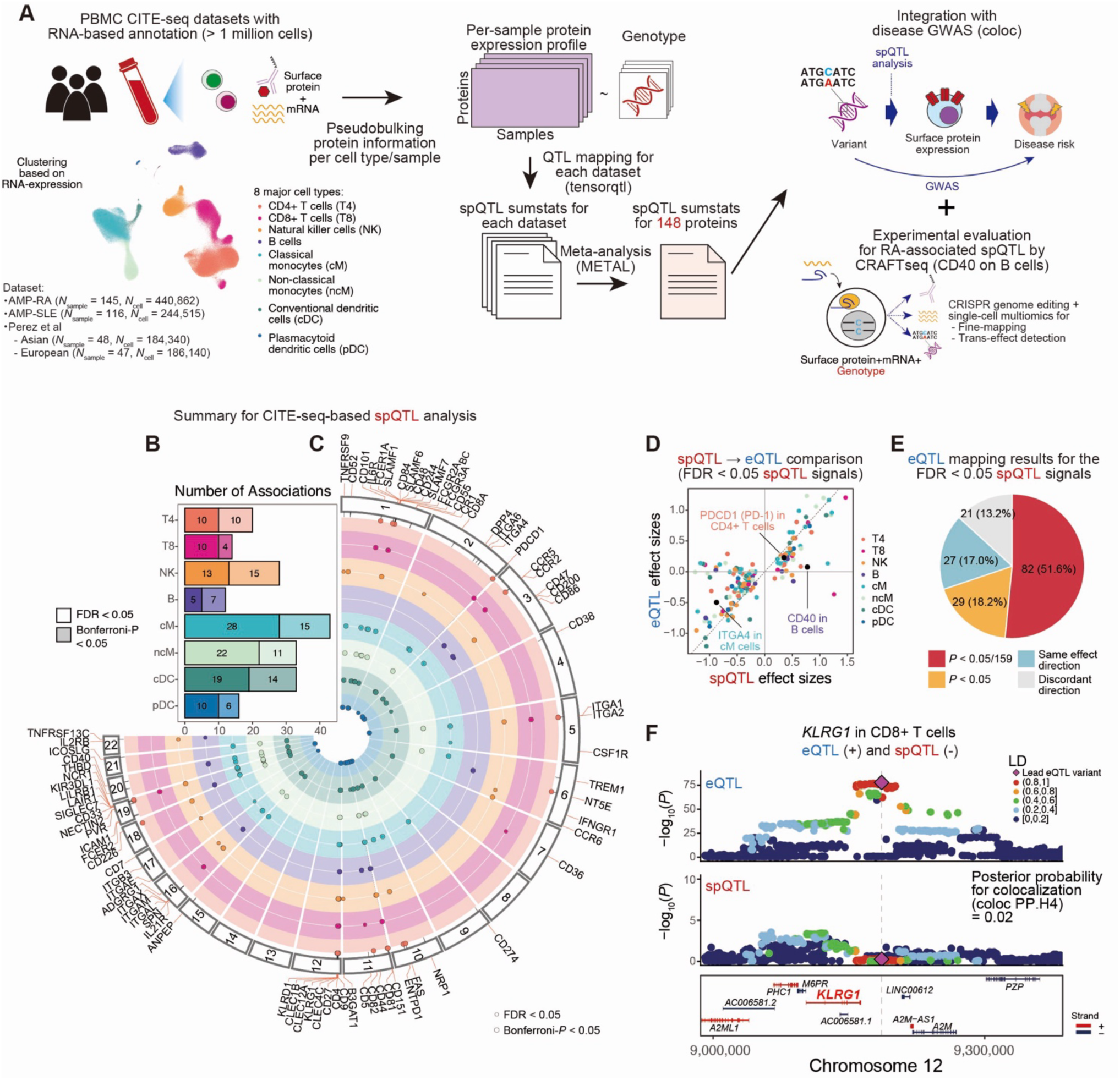
Surface protein quantitative trait loci mapping from CITE-seq data. (**A**) Schematic overview of the study. We utilized four CITE-seq datasets from three studies (AMP-RA, AMP-SLE, Perez et al., Asian/European). We first applied the CLR at the single-cell level and then calculated the mean across cells for each individual and cell type to obtain the pseudobulk expression profile (fig. S2A-E). Then, we performed spQTL mapping for each dataset and meta-analyzed across datasets. Using this spQTL summary statistics, we performed colocalization analysis with disease GWAS to nominate spQTL signals that can mediate variant-disease associations (fig. 2). Focusing on spQTL signals to CD40 on B cells, we performed CRAFTseq for fine-mapping (fig. 3) and *trans*-effect detection (fig. 4 and 5). (**B**) Number of the significant spQTL signals in meta-analysis for each cell type (FDR < 0.05). (**C**) Circos plot showing genome-wide spQTL signals. (**D**) Comparison of the spQTL (x-axis) and eQTL (y-axis) effect size for the lead variant of significant spQTLs. Dots are colored by cell type. Genes highlighted in the fig. 2. are colored with black and labeled. The diagonal dashed line indicates *y* = *x*. (**E**) Pie chart describing the P-values in the eQTL mapping for the lead variant of the significant spQTLs. (**F**) Locus plot showing eQTL (top) and spQTL (middle) signals at the *KLRG1* locus in CD8⁺ T cells, with gene models shown in bottom. Dots are colored according to the LD (*R*^2^) from the lead variant (rs34722215) calculated with 1000 Genome project reference panel (*N* = 2,504). CITE-seq: Cellular Indexing of Transcriptomes and Epitopes by sequencing; CLR: centered log-ratio transformation; eQTL: expression quantitative trait loci; FDR: false discovery rate; GWAS: genome-wide association study; LD: linkage disequilibrium; RA: rheumatoid arthritis; SLE: systemic lupus erythematosus; spQTL: surface protein quantitative trait loci.

## Results

### Identification of cell-type-resolved surface protein QTL from CITE-seq datasets

We mapped spQTLs across eight major cell types from peripheral blood mononuclear cells (PBMC; **Figure 1A, see Methods**). We used four CITE-seq (Cellular Indexing of Transcriptomes and Epitopes by Sequencing) datasets from three independent studies^33–35^. After stringent quality control, we examined 148 unique surface proteins (**Figures S1A to F, and Table S1;** *N*_sample_ = 356, *N*_cell_ = 1,055,857). We mapped spQTLs from pseudobulk surface protein expression profiles of each cell type. We confirmed robustness of spQTL with different pseudobulking methods, and observed that using surface proteins principal components as covariates enhances spQTL detection power (**Figures S2A to G**). Following a meta-analysis across datasets, revealed 199 significant spQTLs spanning 78 distinct proteins (FDR < 0.05; **Figures 1B, C and Table S2**), with consistent effects across datasets (**Figure S3A**). We identified 29 proteins with significant spQTL effects only in a single cell type, and the remaining 49 proteins had significant spQTL effects in more than one cell type (**Figure S4A**). For these 49 proteins with spQTL effects in multiple cell types, colocalization analysis^36^ demonstrated that the genetic effects were independent for 27 of 49 proteins (coloc PP.H3 > 0.5, indicating independent signals; **Figures S4A to D,** see **Methods**). These results suggest that spQTL signals can be cell-type-specific.

To assess whether sqQTLs were mediated through mRNA regulation, we mapped cell-type-resolved eQTL. We augmented a meta-analysis of the four CITE-seq datasets with the OneK1K scRNA-seq dataset (*N*_sample_ = 1,468, *N*_cell_ = 2,404,701)^37^ to increase statistical power (see **Methods**). We evaluated 159 of 199 spQTL lead variant-protein pairs for their eQTLs on the corresponding genes, and 111 (69.8%) spQTL variants had at least nominal eQTL effects (**Figures 1D and E**; *P* < 0.05). However, 48 (30.2%) spQTLs showed no evidence of an eQTL (*P* ≥ 0.05). We note that 22 of 48 (45.8%) spQTL variants tag non-synonymous variants (*R*^2^ > 0.8), which may affect antibody-protein binding (**Figures S4E to G**). Reciprocally, we examined spQTL effects at 207 significant eQTLs for 85 genes (FDR < 0.05; **Table S3**), which involved testing 199 lead variant-gene pairs for association with their corresponding surface proteins. A total of 84 (42.2%) eQTL variants showed no corresponding changes at the surface protein levels (**Figures S4H to J**). For example, *KLRG1* in CD8+ T cells had an eQTL effect with no spQTL effect (**Figures 1F and S4K to M**). This discordance may suggest post-transcriptional buffering.

We compared our spQTLs with independent plasma pQTLs mapped in UKB-PPP^13^. Although plasma protein expression cannot account for cell-type origin, 86 of 136 (64.7%) of spQTL signals reproduced in the UKB-PPP dataset (**Figures S4N and O**; *P* < 3.7 × 10^−4^), supporting the validity of our spQTLs.

### Integrating GWAS loci identifies disease-associated spQTL signals

To identify surface proteins that might mediate complex trait loci, we evaluated associations between spQTL variants and GWAS summary statistics. We prioritized ten autoimmune diseases and two PBMC-related blood cell traits^2,4–6,38–44^ (**Methods**). We focused on spQTLs that were highly significant (*P* < 2.1 × 10^−5^; correction for 199 spQTLs × 12 traits). Colocalization analysis^36^ identified 99 spQTL-GWAS pairs across 19 proteins that shared a common causal signals (coloc PP.H4 > 0.5 indicating a shared causal signal; **Figure 2A and Table S4**). Even after conservatively removing spQTLs tagging non-synonymous variants, 64 colocalizing signals remained.

**Fig. 2.**
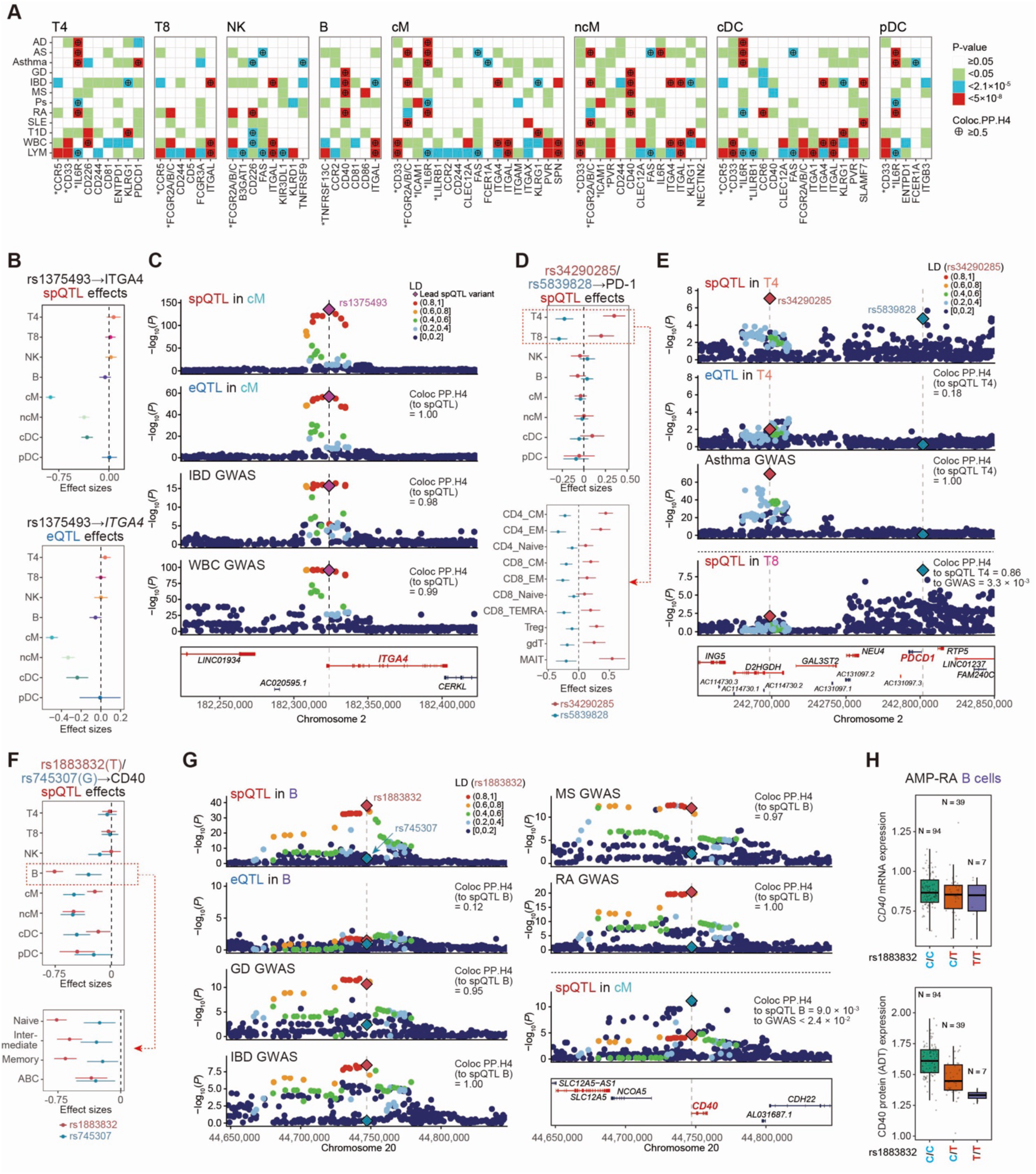
Colocalization between spQTL and GWAS signals. (**A**) Heatmap showing the PheWAS and colocalization analysis between spQTL (x-axis) and GWAS (y-axis). The color of the tiles indicates P-values from PheWAS. For PheWAS associations satisfying the significance threshold (0.05 / 2388 tests = 2.1 × 10^−5^), we performed colocalization analysis with coloc software, and PP.H4 > 0.5 is considered as a colocalization signal. Protein names are prefixed with an asterisk if the lead spQTL variant is in high LD (*R*^2^ > 0.8 in any major 1000 Genomes Project population) with a non-synonymous variant. (**B**) Forest plots showing the spQTL (top) and eQTL (bottom) signals of rs1375493 to ITGA4 for each cell type. Error bars indicate 95% CI and the dashed line indicates *x* = 0. (**C**) Locus plot showing eQTL, spQTL, IBD GWAS, and WBC GWAS signals at the *ITGA4* locus, with gene models shown at the bottom. eQTL and spQTL signals in classical monocytes are indicated. Dots are colored according to the LD (*R*^2^) from the lead spQTL variant (rs1375493) calculated with 1000 Genome project reference panel (*N* = 2,504). (**D**) Forest plots showing the spQTL signals of rs34290285 (red) and rs5839828 (blue) to PDCD1 for each major cell type (top) and T cell annotation (bottom). Error bars indicate 95% CI, and the dashed line indicates *x* = 0. (**E**) Locus plot showing CD4+ T cell spQTL, CD4+ T cell eQTL, Asthma GWAS, and CD8+ T cell spQTL signals at the *PDCD1* locus, with gene models shown in the bottom. Lead spQTL variants in CD4+ T cells (rs34290285; red), and CD8+ T cells (rs5839828; blue) are highlighted. Other dots are colored according to the LD (*R*^2^) from rs34290285 calculated with 1000 Genome project reference panel (*N* = 2,504). (**F**) Forest plots showing the spQTL signals of rs1883832 (red) and rs745307 (blue) to CD40 for each major cell type (top) and B cell annotation (bottom). Error bars indicate 95% CI, and the dashed line indicates *x* = 0. Although the T allele is the reference allele in the hg19 human reference genome, we show the effect size of the T allele because T is a minor allele in the population and the edited allele in most of the following editing experiments. (**G**) Locus plot showing B cell spQTL, B cell eQTL, GD GWAS, IBD GWAS, MS GWAS, RA GWAS, and classical monocyte spQTL signals at the *CD40* locus, with gene models shown in the bottom right. Lead spQTL variants in B cells (rs1883832; red) and classical monocytes (rs745307; blue) are highlighted. Other dots are colored according to the LD (*R*^2^) from rs1883832 calculated with 1000 Genome project reference panel (*N* = 2,504). (**H**) Boxplot showing the expression of CD40 mRNA (top) and surface protein (bottom) across rs1883832 genotypes in the AMP-RA dataset. AD: atopic dermatitis; AS: ankylosing spondylitis; CI: confidence interval; eQTL: expression quantitative trait loci; GD: Graves’ disease; GWAS: genome-wide association study; IBD: inflammatory bowel disease; LD: linkage disequilibrium; LYM: lymphocyte count; MS: multiple sclerosis; PheWAS: phenome-wide association study; Ps: psoriasis; RA: rheumatoid arthritis; SLE: systemic lupus erythematosus; spQTL: surface protein quantitative trait loci; T1D: type 1 diabetes; WBC: white blood cell count.

We identified several signals with clinical implications among these 64 spQTL-GWAS colocalizations. Integrin Subunit Alpha 4 (ITGA4 or CD49d), a critical component of α4β1 or α4β7 complexes, had both eQTL and spQTL effects in classical monocytes (rs1375493; β = –0.49 and –0.85, *P* = 1.3 × 10^−57^ and *P* = 1.7 × 10^−132^, respectively for eQTL and spQTL; **Figures 2B and S5A**). This spQTL signal colocalized with IBD and white blood cell (WBC) counts (**Figure 2C**). Decreased ITGA4 expression reduces risk of IBD and increases WBC. The α4β7 integrin is the target of vedolizumab^45,46^, an established biologic treatment for refractory IBD. Vedolizumab may act on T cells or myeloid cells^47^; our colocalization results suggest that myeloid cells may be the relevant cell type. This highlights how spQTLs may help identify therapeutic targets for drug development.

The PD1 (*PDCD1*) spQTL had independent lead variants in CD4+ (rs34290285; β = 0.35 and *P* = 8.3 × 10^−8^) and CD8+ T cells (rs5839828; β = –0.29 and *P* = 4.4 × 10^−9^; **Figures 2D and S5B,C**; *R*^2^ = 0.0042). Notably, the PD1-increasing allele of rs34290285 in CD4+ T cells colocalized with a reduced asthma risk (**Figure 2E**), while the rs5839828 allele in CD8+ T cells did not. To understand the cell-state specificity of these signals, we mapped the PD-1 spQTL in 10 fine-grained T cell states defined using T-CellAnnoTator^48^ (**Figures 2D and S5D to F**). We found rs5839828 exhibited stronger effects in CD4/8 effector memory and CD8+ TEMRA cells. In contrast, the asthma-associated rs34290285 showed larger effects in CD4+ central memory and MAIT cells, both of which have been implicated in asthma^49,50^. Human observational studies and murine models previously suggested a link between PD-1 and asthma^51,52^. These results suggest that cell-state-specific spQTL effects may drive asthma risk. While rs34290285 exerted a robust spQTL effect, its corresponding eQTL signal was only nominally significance (β = 0.23 and *P* = 0.0099). This discordance likely derived from the limited sensitivity of PDCD1 mRNA detection in CITE-seq (**Figure S5B**), suggesting that spQTL analysis can, in some cases, capture genetic effects that are difficult to find at the mRNA level due to low expression.

### CD40 autoimmune disease locus has a B cell-specific spQTL effect

CD40 is expressed in B and myeloid cells (**Figure S6A**). A CD40 spQTL in B cells colocalized with GWAS for multiple autoimmune diseases (rs1883832-T allele, β = –0.77 and *P* = 7.0 × 10^−39^; PP.H4 > 0.94; **Figures 2F, G**). This allele reduces CD40 protein expression by ∼20% per each T allele (54.7, 45.5, and 36.0 counts per 10K total UMIs for the CC, CT, and TT genotypes, respectively, in the AMP-RA dataset). The rs1883832-T allele decreases RA and GD risk but increases MS and IBD risk. There was an independent CD40 spQTL signal in classical monocytes (lead variant = rs745307; β = –0.51 and *P* = 7.7 × 10^−12^). The autoimmune disease GWAS lead variants were independent of the monocyte signal lead variant (*R*^2^ = 0.038) and did not colocalize with it (coloc PP.H4 < 0.0024, **Figures 2F, G and S6B, C**). This suggested that the disease signal is driven by B cells and not monocytes.

We clustered B cells into 4 states and observed that they all exhibited spQTL effects, with naive B cells showing the strongest effect (**Figure 2F and S6D to F**).

The rs1883832 spQTL has minimal evidence of a B-cell eQTL effect, which was only nominally significant (T allele β = –0.076 and *P* = 0.032; **Figures 2G, H, and S6G, H**). Even if this effect is an eQTL, the variance of mRNA explained was ∼10 times smaller than the surface protein variance explained (*R*^2^ = 0.0010-0.033 and 0.053-0.28 across datasets, respectively for mRNA and surface protein; **Table S5**).

CD40 is a key molecule mediating B–T cell interactions^7^. Within B cells, CD40 signaling drives memory formation, proliferation, and antibody class switching. Notably, CD40 targeting itself is already being investigated in clinical trials^53,54^. Given its high clinical potential, we focused on deep molecular characterization of the CD40 locus.

### Experimental fine-mapping of CD40 loci with single base editing and CRAFTseq

Statistical analysis alone could not resolve the causal variant within the CD40 locus. Within the locus, there are 10 candidate causal variants that are highly statistically significant in RA GWAS (**Figure 3A**). Of those, we selected four candidates overlapping open chromatin regions in B cells^55^ or the gene transcript (**Figure 3A**; rs6074022 [E1], rs4810485 [E2], rs4239702 [E3], and rs1883832 [5’ UTR]). To evaluate the impact of these variants on CD40, we edited Daudi B-cell lines and utilized CRAFTseq to assay mRNA and surface protein (**Figures 3B, S7A, B, and S8A**). With prime editors (PE) or adenine base editors (ABE), we introduced these target variants into Daudi cells in separate experiments. We also edited a splicing donor as a knockout (KO) control (chr20:44,747,035 T>C). To increase editing efficiency, we performed two rounds of editing for PE, though we used a single round of editing for the subsequent experiments (see **Methods**). We confirmed successful editing with Sanger sequencing (**Figure S7A**).

**Fig. 3.**
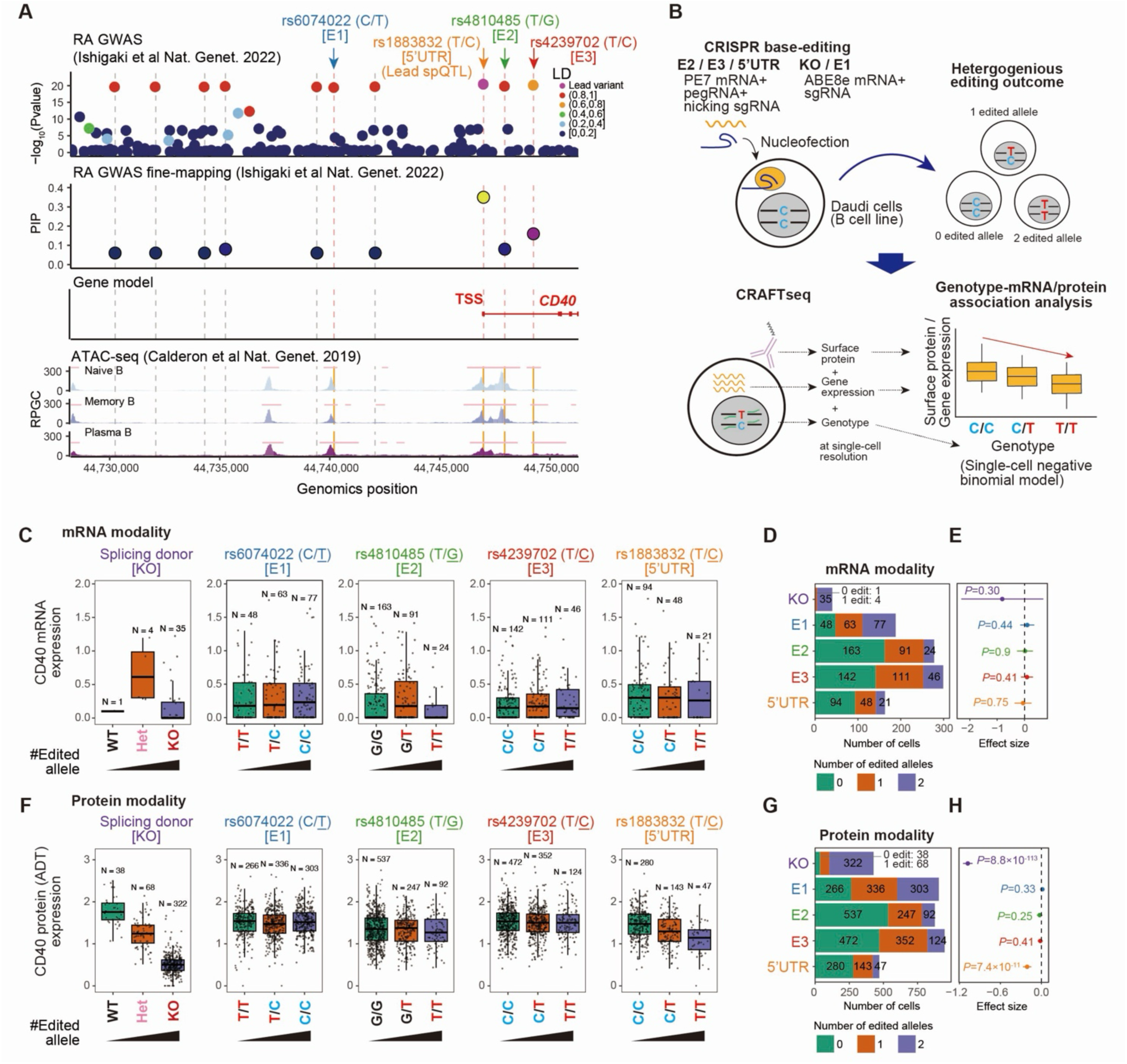
CRAFTseq identifies the causal variant at the CD40 spQTL. (**A**) Locus plot showing the –log_10_(P-values) (first row) and PIP (second row) from RA GWAS with gene models (third row). Dots are colored for LD (*R*^2^) from rs1883832 calculated with 1000 Genome project reference panel (N = 2,504) and PIP, respectively, for the first and second rows. B cell subsets ATAC-seq read coverages are shown in the fourth row, and peaks are indicated with pink horizontal lines. Positions of the variants with P-values < 1.0 × 10^−15^ are indicated with dashed lines, and those overlapping with B cell ATAC-seq peaks are colored pink. (**B**) Schematic overview for the CRAFTseq. We introduced our target variants into Daudi cells using PE or ABE systems (**Materials and Methods**). Then, we performed CRAFTseq, which is a multi-omics technology to obtain mRNA, surface protein, and genotype information at single-cell resolution. We compare the expression of mRNA or surface protein across different genotypes (editing outcomes). (**C**) Boxplot showing the expression of CD40 mRNA across different genotypes of each candidate variant. (**D**) Bar plots showing the cell counts for each edited genomic position used in mRNA-genotype association analysis. Bars are colored according to the number of edited alleles. (**E**) Forest plots showing the association between the expression of *CD40* mRNA and genotypes. Error bars indicate 95% CI, and the dashed line indicates x = 0. (**F**) Boxplot showing the expression of CD40 surface protein across different genotypes in different genotypes of each candidate variant. (**G**) Bar plots showing the cell counts for each edited genomic position used in surface protein-genotype association analysis. Bars are colored according to the number of edited alleles. (**H**) Forest plots showing the association between the expression of CD40 surface protein and genotypes. Error bars indicate 95% CI, and the dashed line indicates x = 0. ABE: adenine-base-editor; ATAC-seq: assay for transposase-accessible chromatin with sequencing; CI: confidence interval; GWAS: genome-wide association study; LD: linkage disequilibrium; PE: prime-editor; PIP: posterior inclusion probability; RA: rheumatoid arthritis; spQTL: surface protein quantitative trait loci.

With CRAFTseq we obtained multimodal single-cell data capturing DNA sequence of the targeted region, the genome-wide transcriptome, surface marker profiles, and CD45 index sorting to hash each editing experiment (see **Methods**). CRAFTseq typically enables us to perform a direct comparison of edited genotypes and expression of 3,000-6,000 genes and 50-150 proteins (**Table S6**). We applied stringent QC requiring (i) gene counts > 500 and mitochondrial gene < 20% for RNA, (ii) surface protein counts > 50 for protein, and (iii) successful genotyping for both RNA and protein (see **Methods**). We obtained 968 and 3,627 high-quality cells for RNA and surface protein analysis, respectively, for all five variants; (**Figures S8B to J and Table S6**).

We tested the effect of editing at single-cell resolution with a negative binomial model (see **Methods**). We detected no significant associations at *CD40* mRNA (*P* > 0.4, **Figures 3C to E and Table S7**). But, only the 5’ UTR variant rs1883832 had a significant effect on CD40 surface protein (T allele β = –0.21, *P* = 7.4×10^−11^; **Figures 3F to H and Table S7**); the effect was consistent with the spQTL. The rs1883832 protein effect remained significant in a conservative analysis restricted to cells passing both RNA and protein QCs (162 cells; β = – 0.14, *P* = 0.0058). This identified rs1883832 as the causal spQTL variant. We replicated the rs1883832 T allele in a larger independent CRAFTseq editing experiment (**Figures S7B and S9A to I)** and again observed a surface protein-specific effect (*N*_cells_ = 2,487, β = –0.14 and *P* = 2.1×10^-13^) with no evidence of an mRNA effect (*N*_cells_ = 957 cells, β = 0.10 and *P* = 0.34; **Figures S9J, K, Table S6 and S7).**

The rs1883832-T allele is located one base upstream of the start codon and disrupts the Kozak motif^56^, essential for efficient translation. This likely explains the protein-specific effect of rs1883832 in B cells.

### The rs1883832 *cis-*effect in primary B cells

Since cell lines may behave differently than primary cells, we wanted to investigate rs1883832 in primary B cells. Primary B cell studies enabled testing of the allelic effect across different genetic backgrounds and gene-editing strategies. Hence, we obtained primary B cells from rs1883832 CC (*N* = 2) and TT (*N* = 1) genotype individuals. We applied both C→T editing with PE and T→C editing with ABE to bidirectionally validate the allelic effects and to control for technical biases due to editing strategies. We cultured cells with CD40L + IL-4 + IL-21 to moderate proliferation and limit terminal differentiation^57^ (**Figure 4A**). CRAFTseq obtained data for 9,870 cells with ≥580 cells for each of the three genotypes per individual (**Figures 4B, S10 A to I, and Table S6**).

**Fig. 4.**
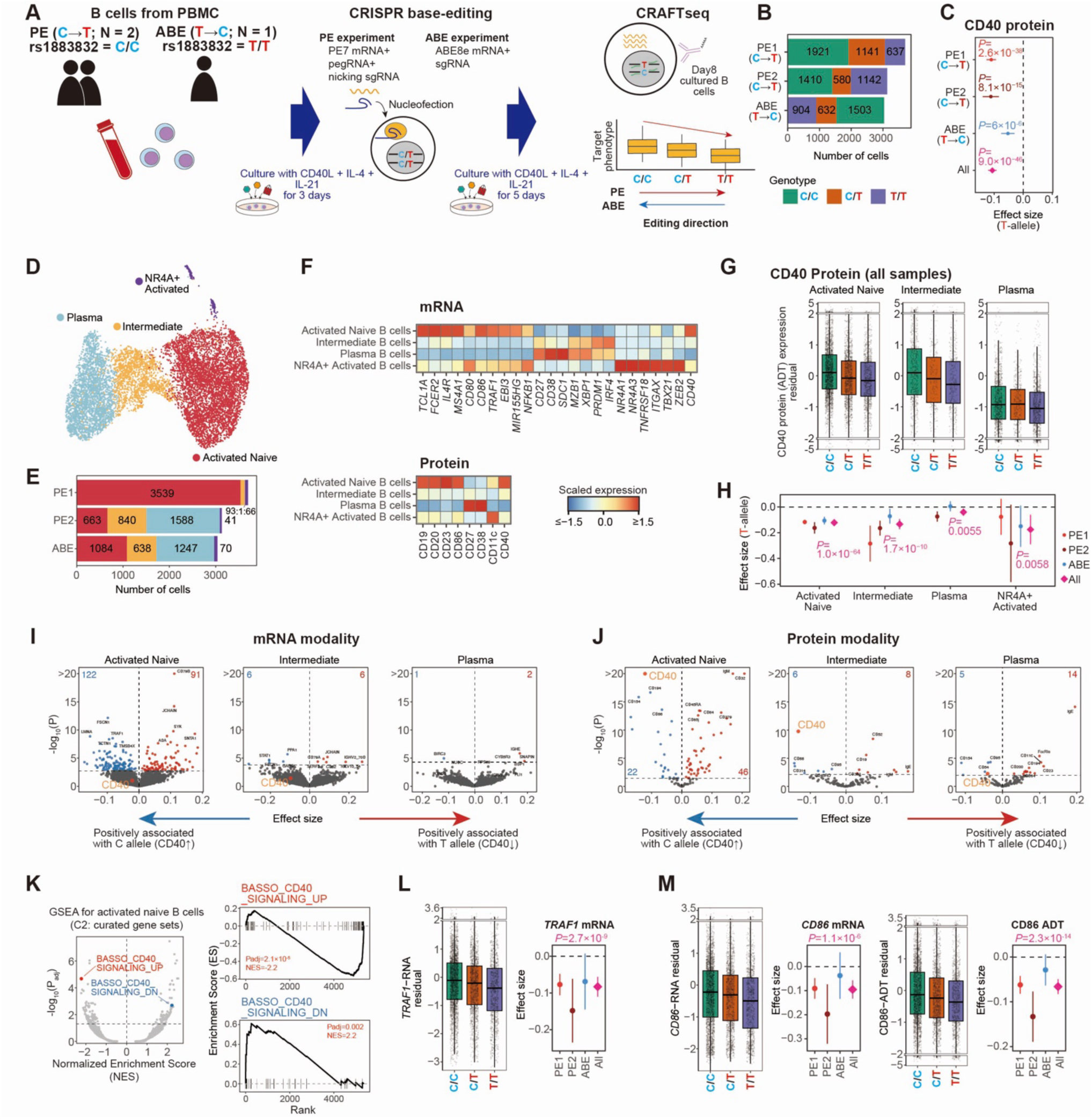
CRAFTseq defined cell-state-specific *cis-* and *trans*-effects of rs1883832 in B cells. (**A**) Schematic overview for the CRAFTseq in primary B cells. Primary human B cells are cultured with CD40L, IL-4, and IL-21 for 3 days and then edited at rs1883832 using prime editing (PE; C→T) or adenine base editing (ABE; T→C). After 5 days of culture with CD40L, IL-4, and IL-21, genotype, transcriptome, and surface protein abundance are jointly profiled at single-cell resolution using CRAFTseq. We compare the expression of mRNA or surface protein across different genotypes (editing outcomes). (**B**) Bar plots showing the number of cells edited for rs1883832 for each experiment. Bars are colored according to the genotype of rs1883832. (**C**) Forest plots showing the association between the expression of CD40 surface protein and rs1883832 genotypes in all the cultured B cells, for each individual experiment and the combined analysis. Error bars indicate 95% CI, and the dashed line indicates *x* = 0. (**D**) UMAP of mRNA profiles for B cells profiled with CRAFTseq. Dots are colored by cell annotation. (**E**) Bar plots showing the number of cells for each experiment. Bars are colored according to the cell annotation. (**F**) Heatmaps of marker gene expression across cell states in the mRNA (top) and surface protein (bottom) modalities. Color scales represent the Z-score calculated from the mean of normalized single-cell-level expression within each cell annotation. (**G**) Boxplots showing CD40 surface protein expression residualized for covariates across rs1883832 genotypes in different cell states. The y-axis includes scale breaks. (**H**) Forest plots showing the association between CD40 surface protein expression and rs1883832 genotypes across cultured B-cell subsets, for each individual experiment and the combined analysis. Error bars indicate 95% CI, and the dashed line indicates *x* = 0. (**I, J**) Volcano plots showing the effect sizes of the rs1883832 genotype (x-axis) and –log_10_(P-values) (y-axis) for all tested traits in the (I) mRNA and (J) surface protein modalities in different cell states. The horizontal dashed line indicates the significance threshold (FDR < 0.05), and the vertical dashed line indicates *x* = 0. Significant positive and negative associations are highlighted in red and blue, respectively, with the total number of significant *trans-*associations indicated at the top of each plot. The top 10 associations with the lowest P-values are labeled. (**K**) Volcano plots showing the normalized enrichment scores (x-axis) and –log_10_(P-values) (y-axis) from the GSEA of the rs1883832 genotype effects on mRNA expression in activated naive B cells (left). GSEA enrichment plots are also shown for pathways related to CD40 signaling pathways (right). (**L**) Boxplots showing *TRAF1* mRNA expression residualized for covariates across rs1883832 genotypes in activated naive B cells. The y-axis includes scale breaks. (**M**) Boxplots showing *CD86* mRNA (left) and surface protein (right) expression residualized for covariates across rs1883832 genotypes in activated naive B cells. The y-axis includes scale breaks. ABE: adenine-base-editor; CI: confidence interval; FDR: false discovery rate; GSEA: gene set enrichment analysis; PE: prime-editor; UMAP: uniform manifold approximation and projection.

First, we checked the *cis-*effect at the mRNA and surface protein modalities. Consistent with Daudi editing, we observed strong protein effects (T allele; β = –0.02 and *P* = 0.10 for mRNA; β = –0.11 and *P* = 9.0×10^−46^ for surface protein) that was consistent across individuals and in both editing methods (**Figures 4C, and S11A to C**).

Next, we wanted to assess if the rs1883832 *cis-*effect was state-specific. We integrate primary B cells across plates with Harmony^58^ into a single embedding and clustered cells into four distinct populations (**Figures 4D to F**). We annotated a cluster of activated naive B cells (expressing *FCER2* and *IL4R* naive marker genes and CD40 downstream genes *TRAF1* and *EBI3*), plasma B-cell cluster (expressing *CD38* and *MZB1*), an intermediate B-cell cluster positioned between the activated naive and plasma B-cell states, and a small cluster of NR4A+ activated B cells. Activated naive B cells showed the highest CD40 mRNA and protein expression (**Figures 4F and S11D**). We evaluated rs1883832 *cis-* effects of the edited allele within each cluster, and observed that it was detectable in all states. It was most pronounced in activated naive B cells (T allele; β = –0.12, *P* = 1.0×10^−64^), and relatively attenuated in the plasma B-cell cluster (β = –0.04, *P* = 5.5×10^−3^; **Figures 4G, H, and S11E, F**). These analyses demonstrated that rs1883832-T allele affects protein levels in cultured primary B cells, especially in activated naive B cells.

### The rs1883832 *trans-*effects in primary B cells are specific to activated naive B cells

Next, we wanted to assess if rs1883832 induces a *trans*-effect. Using the previously described edited primary B cell dataset, we assessed the effect of the rs1883832 edited allele on genome-wide gene and protein expression (**Figure 4A**, 9,870 cells, 3 individuals). We modeled single-cell effects with a negative binomial model (**see Methods**).

We observed that activated naive B-cells exhibited significant *trans-*associations for 213 genes and 68 proteins with consistent effects across all individuals and editing strategies (FDR < 0.05, **Figures 4I, J, S11G, H, and Table S8**). In other B-cell states, we detected little evidence of *trans*-effects. Using gene set enrichment analysis (GSEA), we observed that activated naïve *trans-*genes represented genes positively and negatively regulated by CD40 signaling^59^ (**Figure 4K**).

We identified that some *trans*-genes are implicated in RA pathogenesis. First, we found a negative association between the rs1883832-T protective allele and expression levels of known RA GWAS genes, including those involved in B-cell activation, survival, and proliferation (*TRAF1*, *BATF*, *IRF5*, and *BLK*) and other important genes (*TNFAIP3, LBH, ARHGAP26*; **Figures 4L and S12A to F**). This suggests a common pathway is implicated by RA GWAS loci. We also detected negative association between the rs1883832-T allele and expression of genes involved in B cell-T cell interaction such as *CCR7*, *CD58* (LFA3), and *CD86* (**Figures 4M, S12G and H**). Changes in mRNA and surface protein levels were highly correlated (e.g., CD58, CD86 and FCER2), suggesting that many protein-level effects may be mediated by transcriptional changes (**Figures 4M and S12H to J**). We also validated the reported CD27 protein *trans*-signal^60^; this effect occurred without corresponding mRNA changes suggesting a role for post-transcriptional regulation (**Figure S12K**).

### rs1883832 *trans*-effects are dependent on CD40L stimulation

We hypothesized that rs1883832 *trans*-effects are mediated through CD40 signaling and depend on CD40L engagement. To demonstrate, we used Daudi cells since primary B cells require CD40L for survival in culture.

First, to increase statistical power, we defined CD40L stimulation genes with bulk RNA-seq on CD40L-stimulated and –unstimulated Daudi cells (**Figure 5A and Table S9**). We defined 124 induced and 98 repressed genes; the genes were enriched for CD40 pathway genes (**Figure S13A**).

**Fig. 5.**
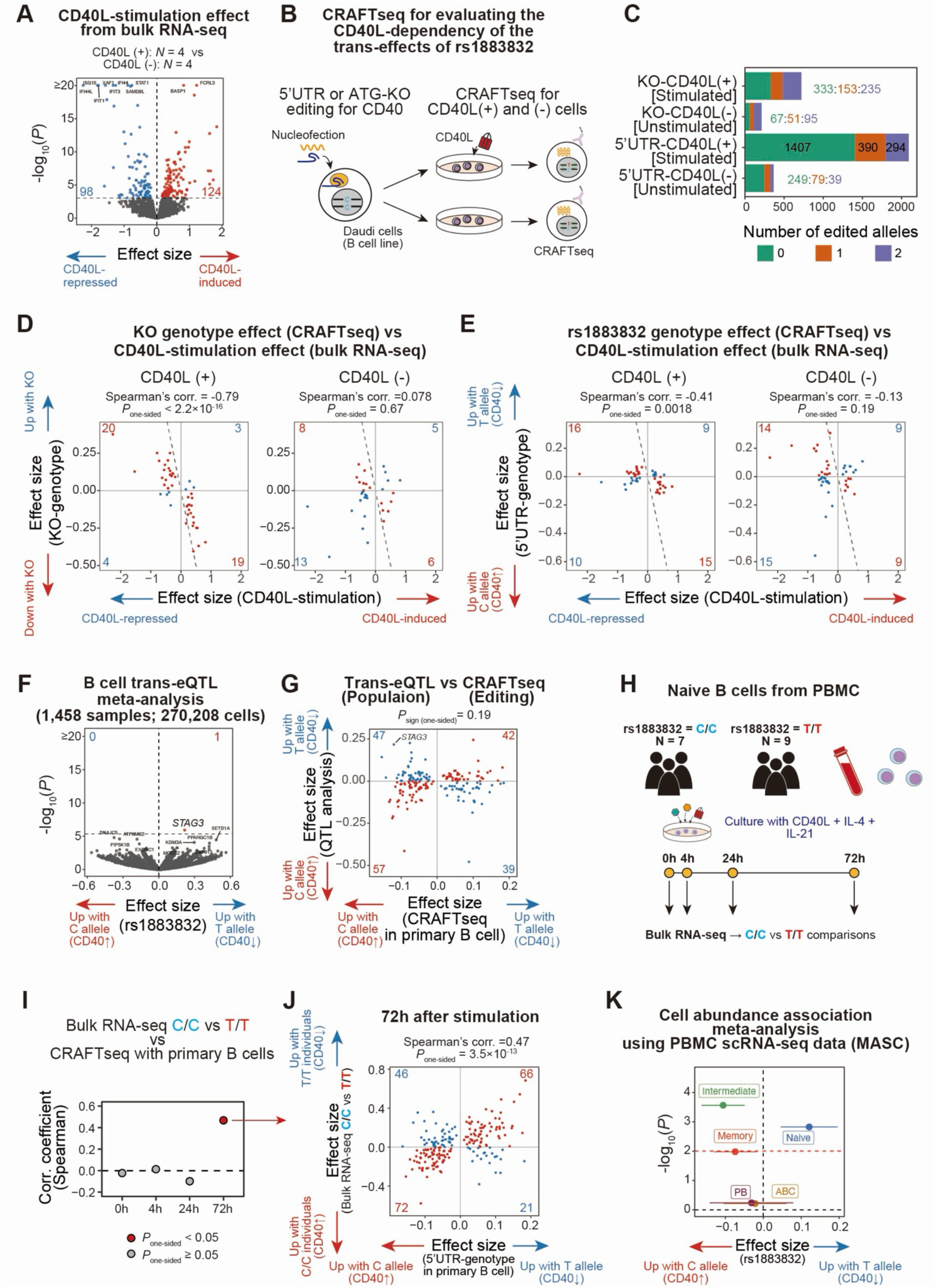
rs1883832 had CD40L-dependent trans-effects affecting B cell-states. (**A**) A Volcano plot showing the effect sizes of the CD40L-stimulation (x-axis) and –log_10_(P-values) (y-axis) for all tested traits in the bulk RNA-seq experiment. The horizontal dashed line indicates the significance threshold (FDR < 0.05), and the vertical dashed line indicates *x* = 0. Significant positive and negative associations are highlighted in red and blue, respectively, with the total number of significant associations indicated at the top of each plot. The top 10 associations with the lowest P-values are labeled. (**B**) Schematic overview for the CRAFTseq in Daudi cells with CD40L-stimulation. We introduced our target variants into Daudi cells using either the PE or ABE system (**Materials and Methods**). Edited Daudi cells were separated into two groups: those stimulated with CD40L and those not. After 48h, cells were profiled for genotype, transcriptome, and surface protein abundance at single-cell resolution using CRAFTseq. We compare the expression of mRNA or surface protein across different genotypes (editing outcomes) and stimulation conditions. (**C**) Bar plots showing the number of edited cells for each variant and CD40L-stimulation condition. Bars are colored according to the genotypes. (**D,E**) Comparison of the CD40L-stimulation effect in bulk RNA-seq (x-axis) and CRAFTseq effect sizes (y-axis) for ATG-KO (D) and 5’ UTR (E). CRAFTseq with Daudi cells stimulated (left) or not stimulated (right) with CD40L is shown. Dots are colored based on the consistency of the effect directions (blue, same direction; red, opposite direction). Consistency of the effect direction is tested with one-sided sign tests. Dashed lines indicate *x* = *y*. Features satisfying FDR < 0.05 in Daudi cell bulk RNA-seq experiment (CD40L stimulated vs unstimulated) are included. (**F**) A volcano plot showing the effect size of rs1883832 (x-axis) and –log_10_(P-values) (y-axis) for all tested genes in the trans-eQTL meta-analysis using B cells from single-cell RNA-seq data. The horizontal dashed line indicates the significance threshold (FDR < 0.05), and the vertical dashed line indicates *x* = 0. Significant positive and negative associations are highlighted in red and blue, respectively, with the total number of significant associations indicated at the top of each plot. The top 10 associations with the lowest P-values are labeled. (**G**) Comparison of effect sizes of rs1883832 from primary B cell CRAFTseq (x-axis; activated naive B cells) and trans-QTL meta-analysis (y-axis) for mRNA. Dots are colored based on the consistency of the effect directions (red, same direction; blue, opposite direction). Consistency of the effect direction is tested with a one-sided sign test. Features satisfying FDR < 0.05 in CRAFTseq are included. (**H**) Schematic overview for the QTL mapping from cultured naive B cells. We recruited individuals with C/C or T/T genotype at rs1883832 and magnetically isolated naive B cells from PBMC. We stimulated naive B cells with CD40L, IL-4, and IL-21, and performed bulk RNA-seq at 0h (before stimulation), 4 h, 24 h, and 72 h. For each time point, we compared mRNA expression profiles between the C/C and T/T genotypes. (**I**) Spearman correlation of effect sizes between bulk RNA-seq QTL mapping and primary B cell CRAFTseq (activated naive B cells) across time points. Genes which satisfied FDR < 0.05 in primary B cell CRAFTseq are included in this analysis. (**J**) Comparison of effect sizes of rs1883832 from primary B cell CRAFTseq (activated naive B cells; x-axis) and bulk RNA-seq (y-axis) at 72 h post-stimulation. Dots are colored based on the consistency of the effect directions (red, same direction; blue, opposite direction). Consistency of the effect direction is tested with a one-sided sign test. (**K**) A volcano plot showing the MASC results for association between rs1883832 and B cell subsets. Red horizontal line indicates the significance threshold after multiple test correction (*P* = 0.05 / 5). FDR: false discovery rate; KO: knockout; MASC: mixed-effects modeling of associations of single cells; PBMC: peripheral blood mononuclear cell; QTL: quantitative trait loci; UTR: untranslated region.

Second, we confirmed CD40L Daudi response genes are rs1883832 *trans-*regulated in primary B cells. We observed that CD40L stimulation differential effect size negatively correlated with the rs1883832-T allele effect, which reduces CD40 in primary B cells (Spearman’s ρ = –0.28, *P* = 0.0056; **Figure S13B**). This supported our approach of using Daudi cells to study CD40L dependency of *trans*-effects.

Third, to assess CD40L-dependence of *trans*-effects, we applied CRAFTseq to edit rs1883832 and KO variants into Daudi cells cultured with or without CD40L (3,392 cells; **Figures 5B,C, S14 A to J, Table S6 and S10**). We observed a more modest rs1883832 *cis*-effect on CD40 protein than that of the KO-allele regardless of CD40L (**Figure S15A**). We observed a strong inverse correlation between CD40L stimulation differential expression effect in bulk RNA-seq data and CD40-KO *trans*-effect of allele in the presence of CD40L (Spearman’s ρ = –0.79, *P* < 2.2 × 10^−16^); the rs1883832 variant exhibited moderate correlation (ρ = –0.41, *P* = 0.0018; **Figures 5D and E**). Notably, without CD40L these correlations were obviated (*P* > 0.19). For instance, *IL2RG* and *BIRC3* exhibited CD40L-dependent *trans*-effects (**Figures S15B and C**). The correlation was robust to down-sampling accounting for differences in numbers of cells cultured with and without CD40L (ρ = –0.57 and –0.25, *P* = 6.0×10^−5^ and 0.048, respectively for KO and rs1883832 variant; **Figure S15D**). Replacing CD40L-dependent genes from the bulk stimulation experiment with CD40L-dependent genes from CD40-KO CRAFTseq experiment obtained similar results (**Figures S16A to E**). These results demonstrate that many rs1883832 *trans*-effects are CD40L-dependent.

### Activated naive B cells are similar to tissue-specific light zone B cells

To understand which *in vivo* B-cell state that rs1883832 *trans*-effects act in, we examined a Tonsil atlas^61^, including diverse B-cell states (**Figure S17A**). This data set contained 155,527 different B and plasma cell states, including Light Zone (LZ) B cells, actively interact with T cells expressing CD40L. We used two strategies: examination of activated naive B-cell marker genes and single-cell reference mapping.

Marker genes from activated naive B-cells, such as *EBI3, TRAF1*, and *MIR155HG,* were expressed in LZ B cells from the Tonsil atlas (**Figures S17B and C**).

To reference map primary B cells from our CRAFT-seq experiment into the Tonsil atlas, we then applied Symphony^62^ to find similar cell states. We found that most CRAFTseq activated naive B cells mapped to LZ B-cell states (85.2%). In contrast, almost no B cells from PBMCs mapped to LZ B cell states (0∼0.4%), but mapped to a separate set of B cells representing memory or non-activated naive B-cell states (**Figures S18A to C**). This is unsurprising since circulating B cells receive minimal CD40L stimulation compared to B cells in lymphoid organs, and lack the activated cell-state.

To understand whether these cells were also present in autoimmune inflammatory tissues, we mapped B cells from inflamed synovial tissue from RA patients ^63^ to the Tonsil atlas. A subset of RA tissue cells annotated as GC-like B cells also mapped to LZ B-cell states (15.1%; **Figures S18A to C**). These data suggested *trans*-eQTL signals may act specifically in a subset of B cells in lymphoid and inflamed tissues that are absent in circulating B cells.

### *Trans*-eQTL effects for rs1883832 is absent in PBMC population data

Given the context-specificity of rs1883832 *trans*-effects, we expected them to be absent in PBMC population data. Conventional *trans*-QTL mapping for rs1883832 using B cells from single-cell PBMC datasets showed only a solitary *trans*-effect at *STAG3*. We observed no correlation with effects identified in primary B-cell CRAFTseq (**Figures 5F,G, and S19A to C**). Since CD40L was absent in these data sets, these context-specific *trans*-effects could not be found.

### *Trans*-eQTL effects for rs1883832 can be reproduced in B cells cultured with CD40L

To validate the condition-specific rs1883832 *trans*-eQTL effects in unedited primary B cells, we stimulated B cells with CD40L, IL-4, and IL-21 from 16 different individuals with TT (N = 9) or CC (N = 7) genotypes. We applied bulk RNA-seq at 0h, 4h, 24h, and 72h after stimulation (**Figure 5H**). Principal component analysis revealed that samples grouped together by time point. At 72h, samples clustered by genotype (**Figure S20A**). We examined *trans*-genes from our primary B-cell CRAFTseq analysis; differential expression between TT and CC genotypes correlated with rs1883832 *trans*-effect sizes only at 72h (Spearman’s ρ = 0.46, *P* = 3.5 × 10^−13^; **Figures 5I, J, S20B,C, and Table S11**). This is consistent with the idea that *trans*-effects are only apparent after substantial CD40L exposure. Since this small study was relatively underpowered, we aggregated differential expression of *trans*-genes positively associated with the T and C alleles in CRAFTseq. On average, at 72h, the expression of T-associated genes exhibited 1.09-fold changes (95%CI = 1.07-1.11; inverse-variance weighted), C-associated genes exhibited 0.98-fold change (95%CI = 0.97-1.00). Thus, we validated rs1883832 *trans*-effects observed in CRAFTseq in unedited primary B-cells in a stimulation-dependent manner.

### rs1883832 modulates B cell subset composition in PBMC

Given the critical role of CD40 signaling in B-cell differentiation^7^, we hypothesized that rs1883832 might influence B-cell differentiation in secondary organs, eventually changing B-cell subset composition in peripheral blood. To investigate this in PBMCs, we applied Mixed Effects Association of Single Cells (MASC) analysis^64^ and found that the rs1883832-T allele was associated with a reduction of intermediate (β = –0.11 and *P* = 2.7 × 10^−4^) and memory B cells (β = –0.074 and *P* = 0.010), alongside an increase in the naive B-cell population (β = 0.12 and *P* = 0.0015; **Figure 5K and Table S12**). These findings are consistent with previous flow cytometry-based GWAS^60^, showing association between rs1883832 and IgD^+^ B-cell populations (β = 0.083 and *P* = 0.0017; **Table S13**). Overall, our data associates rs1883832 with the naive-to-memory B-cell ratio, consistent with the known effect of CD40 signaling on B-cell differentiation ^7^.

## Discussion

A central goal of human genetics is identification of causal variants underlying complex disease loci. We defined the causal allele at the *CD40* autoimmune locus as the rs1883832 5’ UTR variant conferring a *cis*-spQTL effect, and ruled our other variants in LD. Our single-base-resolution editing approach showed that rs1883832 affects CD40 surface expression, with little or no transcriptional effect. Many strategies map cis-regulatory regions, for example, CRISPRi/a tiling screens^22^, and prioritize overlapping variants as causal. Since this variant is close to the CD40 promoter, such approaches would lead to inaccurate inferences about the variant’s mechanism of effect. This highlights the importance of accurately defining the molecular effect of the causal variant.

Trans-effects are notoriously hard to detect^25–29^, but are essential to understand the broader function of disease variants. Current population studies have limited statistical power, miss key cellular contexts, and find indirect non-cellular effects such as cell composition changes. After defining the rs1883832 causal allele, we were able to identify *trans*-effects that were specific to activated naive B cells in the presence of CD40L. These effects were robust across three individuals and two editing strategies. They reproduced in bulk B cells stimulated with CD40L. These results suggest the promise of experimental detection of *trans*-effects via genome editing and single-cell analysis. This is feasible only with knowledge of the causal variant. *In vitro* genome editing enables modeling of the biological context for *trans*-effects while eliminating confounding from cell-composition effects. Single-cell joint profiling of genotype and phenotype, offers high statistical power, and is even more critical for detecting subtle *trans*-effects than *cis*-effects.

Strategies to map the context-specificity of *trans*-effects may have the potential to reveal precise disease-relevant cell-states. The utility of context-specific *cis*-effects in narrowing disease-relevant cell-states is already appreciated^65,66^. The rs1883832 *cis*-effect is restricted to B cells, while its *trans*-effects are specific in activated naive B cells under CD40L stimulation. This *in vitro* state is similar to LZ B cells in tonsils and inflammatory tissues. The rapid advancement of genome-editing tools and immune organoid models^67^ may enable SNP investigations under more physiological conditions in the future.

Non-specific B-cell depletion therapies, such as rituximab^68^, are effective against autoimmunity, a concept further highlighted by the recent success of anti-CD19 CAR-T^8–10^. However, pan-B-cell depletion carries life-threatening risks, such as infection and cytokine release syndrome. It remains unclear whether a cure requires broad B-cell depletion or if targeting pathogenic B cells is sufficient. The states where GWAS variants act may be promising targets for curative therapies. Interestingly, the rs1883832 *trans*-targets include RA GWAS risk genes, suggesting a shared context-specific molecular process. The next essential step is defining causal variants across multiple GWAS loci to identify the shared pathways or cell states where they converge.

The rs1883832-T allele is protective for RA and GD, but confers risk for MS and IBD^60,69^. Although B-cell activation drives immune responses, it may also induce suppressive regulatory B cells or T cells^70^. Furthermore, effective microbial control is critical to maintain immune homeostasis as exemplified by the causal relationships between Epstein-Barr virus and MS^71^, and gut bacteria and IBD^72^. This underscores the complexity of the interplay between the immune system and autoimmune diseases; hence, rs1883832 may have disease-specific effects.

We demonstrate the utility of a cell-type-resolved spQTL analysis. Researchers often focus on mRNA for its ease of assay, yet it is a proxy for surface proteins^73^. In addition, some variants can specifically impact protein regulation rather than gene expression, as illustrated by the rs1883832 Kozak motif variant. For this variant, eQTL data alone can be misleading; it showed an eQTL effect in PBMC in previous studies^74^ and in monocytes in our study (**Figure S6H**). Our spQTL analysis is limited by the antibody-based assays, which vary in sensitivity across proteins and are potentially affected by non-synonymous variants (**Figures S4E to G**). Complementary technologies such as mass spectrometry offer potential for the future^75^.

In summary, the combination of spQTL mapping and single-cell resolution editing enabled functional definition of the *cis* and *trans* molecular effects of the rs1883832 disease allele. Our approach addresses a critical challenge in population genetics, and offers a powerful framework to accelerate the discovery of disease etiology from genetic association signals.

## Methods

### Preprocessing of the published genotype data

For the Perez et al dataset^33^, genotype data was processed as described by Rumker et al^76^. Briefly, genotype data downloaded from dbGAP (phs002812.v1.p1) was processed separately for each genotyping array (LAT and Omni). Following the exclusion of non-Asian/European ancestries and related individuals (PI_HAT > 0.125), standard quality control was performed using PLINK (v1.90)^77^. Variants were filtered based on missingness (< 1%), minor allele frequency (MAF; > 1%), and concordance with 1000 Genomes Project allele frequencies. Samples with outlier heterozygosity (> 3 SD) were also removed. Finally, the post-QC genotypes were phased with SHAPEIT2^78^ and imputed using Minimac3^79^ with the 1000 Genomes Project reference panel^80^. Following imputation, variants with high imputation quality (*R*^2^) > 0.7 were used for downstream analyses. Genotype principal components (PCs) were calculated with PLINK.

For AMP datasets^34,35^, genotyping was performed using the Illumina Multi-Ethnic Genotyping Array, as previously described by Kang et al^81^ and Sakaue et al^82^. Genotype data were processed with PLINK for quality control, where variants with high missingness (> 1%), low minor allele frequency (MAF < 0.01), and deviation from Hardy-Weinberg equilibrium (*P* < 10^−6^) were removed. Genotypes were phased with SHAPEIT2 and subsequently imputed across the whole genome using minimac3 in conjunction with the 1000 Genomes Project Phase 3 reference panel. Following imputation, variants with high imputation quality (*R*^2^) > 0.7 were used for downstream analyses. Genotype PCs were calculated with PLINK.

For the OneK1K dataset^37^, SNP array genotypes were processed as described in Rumker et al^76^. Briefly, variants were aligned to the hg19 reference, excluding duplicated and palindromic SNPs. Quality control was performed using PLINK, retaining samples with a call rate > 99% and variants with a call rate > 98%. Ancestry was confirmed as European by projecting samples onto the 1000 Genomes Phase 3 reference. Post-QC genotypes were phased with SHAPEIT2 and imputed via Minimac3 using the 1000 Genomes reference panel. Only variants with an imputation *R*^2^ > 0.7 were retained for downstream analysis. Genotype PCs were calculated with PLINK.

### Preprocessing of the published CITE-seq and scRNA-seq datasets

For the Perez et al dataset, we used already quality-controlled scRNA-seq data, which were publicly available (GSE174188). Briefly, Single-cell RNA-seq (scRNA-seq) data from the Perez et al. dataset were generated using the 10X Chromium Single Cell 3’ v2 chemistry. Peripheral Blood Mononuclear Cells (PBMCs) were isolated, pooled, and loaded onto the 10X Chromium Controller, with libraries subsequently sequenced on HiSeq 4000 or NovaSeq 6000 platforms. For surface protein detection, cells from processing batches 3 and 4 were stained with AbSeq antibodies (Becton Dickinson) after Fc blocking before loading on to the 10X Chromium Controller. Data preprocessing followed a multi-step pipeline: donor demultiplexing was achieved using freemuxlet^83^, while doublets were further excluded using Scrublet^84^. To ensure high data quality, contaminating populations, including platelets and red blood cells, were removed based on the expression of specific markers (e.g., PF4 and hemoglobin genes). The remaining singlets underwent normalization and technical batch correction using Scanpy^85^. Cell type identities were assigned through Louvain clustering and validated using CITE-seq surface protein expression. For the visualization of the cell annotation, we used the Uniform Manifold Approximation and Projection (UMAP) from the original study. To obtain the surface protein modality data (AbSeq) which was not included in the publicly available scanpy object, we processed the fastq files downloaded from the GEO database (GSE174188) using CITE-seq-Count software (https://github.com/hoohm/cite-seq-count) with the following options: –cbf 1, –cbl 16, –umif 17, –umil 26, –cell 30000, and –T 16. For surface protein analysis, we used only cells from processing batch 4, which is profiled with a larger antibody panel (99 target proteins) than processing batch 3 (16 target proteins). Then, surface protein ADT counts were retained for cells that were profiled in the above scRNA-seq data.

For the AMP-RA and AMP-SLE datasets, we used a dataset which was already quality-controlled by our previous studies^34,35^. Briefly, PBMCs were collected, cryopreserved in CryoStor CS10, and processed using a standardized CITE-seq protocol. Then, samples were thawed at 37°C and batches of up to 16 samples were stained with fluorescent antibodies (CD15 and CD45) and the TotalSeq-A Human Universal Cocktail V1.0 for surface protein detection. To ensure high-quality profiling, live CD45+ CD15-cells were isolated via FACS, with individual donors in each sub-pool (4 donors per sub-pool) distinguished by unique CD45 fluorochromes. All the cells from sub-pools were pooled and loaded onto the 10x Chromium platform for library preparation. Sequencing was performed on the Illumina NovaSeq platform. Data preprocessing was conducted using 10x Genomics Cell Ranger (v3.1.0 for AMP-RA; v6.1.1 for AMP-SLE), followed by donor demultiplexing using genotype-based matching with demuxlet^86^. For quality control, we excluded doublets (demuxlet for both; scDblFinder^87^ for AMP-RA; Scrublet for AMP-SLE) and removed low-quality cells with fewer than 500 detected genes or more than 20% mitochondrial unique molecular identifier (UMI). Then, to define major cell types using RNA expression alone, we assigned cell type annotation based on the expression of marker genes. Briefly, UMI counts were normalized to 10,000 per cell and log-transformed using the Seurat R package^88^. Then, we selected the top 3,000 highly variable genes (HVGs) and performed z-scaling and PCA. We applied the Harmony algorithm^58^ to correct for batch effects associated with individuals. Major cell types were identified through a two-stage clustering strategy. First, to classify cells into major lineages—specifically T/NK cells, B cells, myeloid cells, and proliferating cells—we applied the FindNeighbors and FindClusters functions to harmony-corrected PCs in Seurat R package using default settings at a resolution of 1.2. Then, harmony-corrected PCs are calculated from the top 2000 HVGs separately for each lineage of cells and again clustered with FindNeighbors and FindClusters with resolution of 2 (myeloid cells) or 3 (others) for defining major cell types (CD4+ T cells, CD8+ T cells, NK cells, B cells, Plasmablasts, classical monocytes, non-classical monocytes, conventional dendritic cells, and plasmacytoid dendritic cells). After the annotation of major cell types, we visualized cell annotation on the UMAP space calculated from the top 30 harmony-corrected PCs with the RunUMAP function in Seurat R package. Cell annotations were further validated by the expression of surface protein markers.

For the OneK1K dataset, we used a dataset which was already quality-controlled by previous studies^37,76,81^. Briefly, the analysis was limited to individuals of European ancestry with available genotype data. After the quality control by original study, the dataset maintained high-quality single-cell profiles, where all retained cells were confirmed to have ≥660 UMIs, >230 unique genes, and <7.8% mitochondrial reads. Doublets were excluded by a cluster-based approach. The counts matrix was normalized to 10,000 counts per cell and scaled using Scanpy. Batch effects across independent sequencing pools were corrected using Harmony, and UMAP was generated by Scanpy with default parameters. Cell type annotation was assigned to each cell by Azimuth reference mapping^89^ in the original publications. To align the fine-grained annotations from the previous publication with our major cell-type level annotations, we consolidated the original annotations into following categories: CD4+ T cells (T4; including CTL, Naive, TCM, TEM, and Treg), CD8+ T cells (T8; including Naive, TCM, TEM, and MAIT), NK cells (NK; CD56bright and NK), B cells (B; Naive, Memory, and Intermediate), Plasmablasts (PB), classical monocytes (cM; CD14 Mono), non-classical monocytes (ncM; CD16 Mono), conventional dendritic cells (cDC; cDC1 and cDC2), and plasmacytoid dendritic cells (pDC). For the visualization of the cell annotation, we used the UMAP from the previous study.

To confirm the cell annotations, we visualized mRNA and surface protein of marker genes by heatmap (**fig. S1D and E**).

### Generation of pseudobulk surface protein expression profiles for spQTL mapping

To identify surface proteins with signals exceeding background levels, we calculated the Kullback-Leibler (KL) divergence for each protein^90^. Specifically, we compared the distribution of cells with normalized expression above the 90th (KLD_90_) or 99th (KLD_99_) percentiles across major cell-type clusters against the global distribution of all cells across broad clusters. Proteins failing to meet both thresholds of KLD_90_ < 0.3 and KLD_99_ < 0.07 were excluded. Note that all negative control proteins, including EpCAM in the Perez et al. dataset and isotype control IgGs in the AMP-RA and AMP-SLE datasets, were removed with these thresholds. Following single-cell-level feature selection, protein expression data were normalized using a centered-log-ratio (CLR) transformation. Pseudobulk profiles were then generated by calculating the mean expression within each individual for each specific cell type (CLR→Mean method). After pseudobulking, we excluded samples containing fewer than five cells. We restricted our analysis to proteins encoded by autosomal non-HLA genes that were expressed in more than 50% of samples with antibody-derived tag (ADT) UMI counts ≥ 5. Protein to gene mapping was based on reference files released from the manufactures (Perez et al [Abseq: https://scomix.bd.com/hc/en-us/articles/360048878592-AbSeq-Data-Analysis-and-Reference-files, Total-seq: https://www.biolegend.com/Files/Images/BioLegend/totalseq/TotalSeq_A_Human_Universal_Cocktail_v1_163_Antibodies_399907_Barcodes.xlsx). Finally, a rank-based inverse normal transformation was applied to the normalized individual-level protein expression profiles prior to subsequent spQTL mapping.

### spQTL mapping and meta-analysis across datasets

We used TensorQTL^91^ with default parameters for spQTL mapping (window size = ±1Mbp, mode = cis_nominal, and maf_threshold = 0.1 for Perez et al dataset and 0.05 for AMP-RA and AMP-SLE datasets). We included following covariates for each dataset: Perez et al.— age, log(number of cells), batch, disease status (control or SLE), first four genotype PCs, first two protein expression PCs (sex was excluded as all samples were female); AMP-RA—age, sex, log(number of cells), batch, disease status (control, at-risk for RA, or RA), first five genotype PCs, first five protein expression PCs; AMP-SLE—age, sex, log(number of cells), batch, disease status (control or SLE), first five genotype PCs, first five protein expression PCs. After obtaining spQTL summary statistics for each dataset (Perez et al., Asian/European, AMP-RA, AMP-SLE), we performed meta-analysis using METAL^92^ (’SCHEME STDER’ option). To calculate protein-level P-values, we conservatively multiplied the number of tested variants by the lowest P-value for the gene, then applied the Benjamini-Hochberg (BH) method to determine FDR. Proteins with FDR < 0.05 were defined as significant spQTL signals. To define spQTL signals potentially driven by non-synonymous variants, we extracted all variants in Linkage disequilibrium (LD; *R*^2^ > 0.8) with the lead spQTL variant in any of the five major population groups from the 1000 Genomes Phase 3 data. Annovar^93^ was used for defining non-synonymous variants for the targeted protein in spQTL analysis. To evaluate the consistency of spQTL effects across datasets, we extracted lead spQTL variants from the meta-analysis results and compared the effect sizes in each dataset in a pairwise manner.

### Comparison of different methods for spQTL mapping

To evaluate the impact of pseudobulking and normalization methods for spQTL mapping, we tried two alternative normalization strategies for the AMP-RA dataset: aggregating raw ADT counts by individual and cell type, followed by either (i) log transformation (Aggregate→Log) or (ii) CLR transformation (Aggregate→CLR) in addition to the CLR→Mean method used in the main analysis. For evaluating the performance, we checked the numbers of significant spQTL signals detected by each pseudobulking and normalization strategy. We also evaluated the P values for lead variants, restricted to proteins consistently significant across all pseudobulking and normalization strategies. These analyses demonstrated that spQTL mapping is largely robust to the choice of pseudobulking and normalization method, with highly concordant results across all tested methods. However, among the robustly identified spQTL hits, the CLR-transformation followed by mean-based pseudobulking consistently yielded marginally lower P-values compared to other approaches. Consequently, we adopted the CLR-Mean method for our main analysis.

We also evaluated whether including surface protein expression PCs as covariates, analogous to mRNA expression PCs in eQTL studies^94^, improves the power of spQTL mapping. Using the AMP-RA dataset, we calculated expression PCs from the CLR-normalized mean protein profiles and performed mapping by incorporating 0, 1, 2, 3, 4, 5, 7, or 10 PCs as covariates. Other covariates were the same to our main analysis, namely age, sex, log-transformed cell count, batch, disease status, and the first five genotype PCs. we checked the numbers of significant spQTL signals and P values for lead variants, restricted to proteins consistently significant across all different numbers of expression PCs. We found that the inclusion of expression PCs increased the number of significant hits and generally improved the P-values for robust associations. This improvement was most pronounced within the range of 0 to 5 PCs; therefore, we selected first five expression PCs for datasets with a scale of approximately 150 samples (AMP-RA and AMP-SLE). For the Perez dataset (–50 samples), we used the first two PCs, following previous study scaling down the number of unobserved covariates in a data-driven manner according to sample size^95^.

### Cell-type specificity evaluation for the spQTL signals

First, we evaluated the number of cell types where each protein has significant spQTL effects (FDR < 0.05). Also, for defining spQTL signals shared within lineages, we clumped CD4+ T cells and CD8+ T cells into ‘T lineage’ and classical monocytes, non-classical monocytes, and conventional dendritic cells into ‘Myeloid lineage’. For proteins with more than one cell type with spQTL effects, we performed colocalization analysis of spQTL results across cell types using the coloc R package^96^. For cell type pairs that showed the highest PP.H3 (posterior probability of two distinct causal variants), we performed further evaluation of the cell type-specificity using a conditional analysis approach. We performed spQTL mapping for cell type 1 after conditioning on the lead spQTL variant in cell type 2, while keeping other covariates unchanged from the original analysis. As an orthogonal approach, we also evaluated the heterogeneity of the spQTL effects for all the lead variants with *I*^2^ statistics calculated with the metafor R package^97^.

### eQTL mapping and meta-analysis across datasets

To evaluate the concordance between eQTL and spQTL effects, we performed eQTL analysis using Perez et al (Asian/European), AMP-RA, AMP-SLE, and OneK1K datasets. We aggregated raw UMI counts by individual and cell type, followed by log-transformation using Seurat R package. After pseudobulking, we excluded samples containing fewer than five cells. We restricted our analysis to genes encoded by autosomal genes that were expressed in more than 25% (OneK1K) or 50% (Perez et, AMP-RA, and AMP-SLE) of samples with UMI counts ≥ 5. Finally, a rank-based inverse normal transformation was applied to the normalized individual-level protein expression profiles prior to subsequent eQTL mapping. We used TensorQTL with default parameters for eQTL mapping (window size = ±1Mbp, mode = cis_nominal, and maf_threshold = 0.05). For covariates in each dataset: Perez et al.—age, sex, log(number of cells), batch, disease status, first four genotype PCs, first five gene expression PCs; AMP-RA—age, sex, log(number of cells), batch, disease status, first five genotype PCs, first five gene expression PCs; AMP-SLE—age, sex, log(number of cells), batch, disease status, first five genotype PCs, first five gene expression PCs; OneK1K—age, sex, log(number of cells), batch, first six genotype PCs, first ten gene expression PCs. After obtaining spQTL summary statistics for each dataset (Perez et al., Asian/European, AMP-RA, AMP-SLE, OneK1K), we performed meta-analysis using METAL software (’SCHEME STDER’ option). We extracted genes evaluated in the spQTL analysis and calculate gene-level P-values by conservatively multiplying the number of tested variants by the lowest P-value for the gene, then applied the BH method to determine FDR. Genes with FDR < 0.05 were defined as significant eQTL signals.

### Comparison of spQTLs with eQTLs and plasma pQTLs

To compare genetic effects across different modalities (spQTLs, eQTLs, and plasma pQTLs), we extracted significant QTLs (FDR < 0.05) from a reference modality (either spQTL or eQTL) and evaluated the effect sizes of their lead variants in a query modality (reference→query: spQTL→eQTL, eQTL→spQTL, or spQTL→plasma pQTLs). For each significant lead variant identified in the reference modality, we further assessed its effect direction and P-value within the query modality. When using spQTLs as the reference, we performed stratified analyses based on whether the lead variant tagged a non-synonymous variant. For colocalization analysis between modalities, we utilized the coloc R package, incorporating all intersecting variants within the *cis*-window. For the integration of plasma pQTL summary statistics, we utilized data from the UKB-PPP study^13^, a large-scale proteomic analysis based on Olink technology. This comparative analysis focused on 118 proteins that overlapped between our study and the UKB-PPP dataset. For visualization with a locus plot, we calculated LD information from all independent samples in the 1000 Genomes Phase 3 data (*N* = 2,504).

### PheWAS and disease GWAS colocalization analysis for spQTL signals

To evaluate whether the identified spQTL signals share common causal variants with those of complex traits in GWASs, we performed a phenome-wide association study (PheWAS) and colocalization analysis using the R package coloc. We integrated our spQTL summary statistics with public GWAS data. We prioritized the following 10 immune-related diseases and 2 blood cell count traits: Atopic dermatitis^39^, Ankylosing spondylitis^44^, Asthma^42^, Graves’ disease^6^, Inflammatory bowel disease^5^, multiple sclerosis^4^, Psoriasis^40^, Rheumatoid arthritis^2^, systemic lupus erythematosus^43^, type 1 diabetes^41^, white blood cell counts^38^, and lymphocyte counts^38^. We integrated our spQTL summary statistics with GWAS data by harmonizing genomic positions and alleles. For significant spQTLs, we evaluated the GWAS P-value of the variant that had the smallest P-value in spQTL. We set a significance threshold for PheWAS at 2 × 10^−5^ (0.05 / 2388 tests) and applied colocalization analysis with coloc R package for the significant PheWAS associations. We designated loci with significant colocalization as those with posterior probability of H4 (“both traits are associated and share a single causal variant”) > 0.5. The admixed ancestry of the spQTL cohort and LD discrepancies with the GWAS cohorts may limit the power of this analysis, as previously mentioned^65^. For visualization with a locus plot, we calculated LD information from all independent samples in the 1000 Genomes Phase 3 data (*N* = 2,504).

### Evaluation of spQTL signals at the *PDCD1* loci across T cell subsets

First, we defined T cell subsets using T-CellAnnoTator^48^ (TCAT; https://github.com/immunogenomics/starCAT). We used discrete annotation in the ‘Multinomial_Label’ column of the *.scores.txt files. Then, we performed pseudobulking and normalization in a similar way to the main analysis, except pseudobulking was performed separately for each T cell subset. For spQTL mapping and meta-analysis, we followed the same pipeline as the main analysis using the same set of covariates. For conditional analysis, we added either the genotypes of rs34290285 or rs5839828 as a covariate and performed spQTL mapping and meta-analysis using the same pipeline as the main analysis, using the same set of covariates. For visualization with a locus plot, we calculated LD information from all independent samples in the 1000 Genomes Phase 3 data (*N* = 2,504).

### Evaluation of spQTL signals at the *CD40* loci across B cell subsets

B cell subsets were defined through manual annotation using the Seurat R package. We initially extracted all B cells and plasmablasts from each single-cell RNA-seq dataset. The data were log-normalized, and the top 2,000 highly variable genes were selected for downstream analysis. Following scaling and PCA, batch effects were corrected using the Harmony algorithm, with “individual” specified as the batch term across all datasets. Clustering was performed on the Harmony-corrected PC space using the FindNeighbors (default parameters) and FindClusters (resolution = 3.0) functions. B cell subsets (naive B, intermediate B, memory B, age-associated B (ABC), and plasmablasts) were manually assigned to each cluster based on the expression of marker genes (**fig. S6D**).

Then, we performed pseudobulking and normalization in a similar way to the main analysis, except pseudobulking was performed separately for each B cell subset. For spQTL mapping and meta-analysis, we followed the same pipeline as the main analysis using the same set of covariates. For conditional analysis, we added either the genotypes of rs1883832 or rs745307 as a covariate and performed spQTL mapping and meta-analysis using the same pipeline as the main analysis, using the same set of covariates. For visualization with a locus plot, we calculated LD information from all independent samples in the 1000 Genomes Phase 3 data (*N* = 2,504).

### Calculation of variance explained by rs1883832 for spQTL and eQTL

To quantify the proportion of variance in surface protein/gene expression explained by rs1883832, we fit two linear regression models using lm() function in R per cohort: a full model including allele dosage of the variant as a predictor alongside all covariates used in the spQTL/eQTL analysis, and a null model including covariates only. The difference of R² between full model and null model was used as the estimate of variance explained by the variant.

### *Trans*-QTL mapping for rs1883832 in B cells

To evaluate the *trans*-effects of rs1883832 on genome-wide gene expression, linear regression analyses were performed using the lm() function in R. These analyses utilized the quality-controlled, normalized expression profiles and covariate data for the spQTL and eQTL mapping as mentioned above. To synthesize the results across cohorts, we conducted fixed-effects meta-analyses using the metafor package in R. Specifically, for eQTL mapping, a meta-analysis was performed across the Perez et al. (Asian and European), AMP-RA, AMP-SLE, and OneK1K datasets. For spQTL mapping, the meta-analysis integrated the Perez et al. (Asian and European), AMP-RA, and AMP-SLE datasets.

### Defining candidate causal spQTL variant at the *CD40* loci

We used GWAS and fine-mapping results from our previous study^2^. Although rs6074022 was not included in the 95% credible set, we included this variant in our candidate variant set because this variant had small P-value (P = 4.2 × 10^−20^). For visualization with a locus plot, we calculated LD information from all independent samples in the 1000 Genomes Phase 3 data (N = 2,504). To identify variants overlapping with B cell ATAC-seq peaks, we used B cell ATAC-seq datasets from a previous study^55^. We used fastp^98^ for read QC (paired-end mode; –3 –5 –W 4 –M 20 –w 2 –q 30 –n 5 –l 25) and mapped to human reference genome (hg19) with bowtie2^99^ with default parameters and a maximum paired-end insert distance of 2 kbp following the original study. Following alignment, high-quality reads were retained by filtering for properly paired, non-duplicate reads with a mapping quality (MAPQ) score ≥ 30, while excluding unmapped, secondary, and low-quality alignments (–f 2 –F 1804) following the original study. Reads were then restricted to autosomal and sex chromosomes (chr1–22, X, and Y). PCR duplicates were identified and removed using Picard MarkDuplicates. ATAC-seq peaks were identified using MACS2^100^ with the following parameters following the original study: –-nomodel –-nolambda –-keep-dup all –-call-summits. A genome size of 2.7 × 10^9^ bp was used for the effective human genome. Identified peaks were merged using bedtools merge^101^, and peaks exceeding 3,000 bp were excluded. Overlap between RA-GWAS variants and B cell ATAC-seq peaks were identified using findOverlaps function in rtracklayer^102^ R package.

### Daudi cell line cultures and genomic editing for fine-mapping CRAFTseq experiment

For cell line culture and editing experiments, we followed the protocols from our previous publication with slight modifications^23^. Daudi cells (ATCC, CCL-213) were cultured in complete RPMI (cRPMI), RPMI 1640 supplemented with 10% heat-inactivated FBS and 1% non-essential amino acids, sodium pyruvate, HEPES, l-glutamine, penicillin-streptomycin, and 0.1% β-mercaptoethanol. In the editing experiment for rs1883832, rs4810485, and rs4239702, we used 1.5 μl of mRNA (1 μg/μl) encoding the prime editor PE7^103^, 1 μl of 50 μM modified nicking sgRNA (Synthego), and 1μl of 100 μM pegRNA (IDT). Cells were nucleofected with an mRNA/sgRNA/pegRNA mixture in an Amaxa 4D nucleofector (SF protocol: CA-137). pegRNAs were designed with PRIDICT2.0^104^ (https://pridict.it). Cells were immediately transferred to 24-well plates with pre-warmed media and cultured. Since editing efficiency was low in the prime-editing experiment, we performed nucleofection twice (day 0 and day 5), and cells were harvested for CRAFTseq on day 10. In the editing experiment for rs6074022 and splicing donor KO, we used 0.5 μl of mRNA (1 μg/μl) encoding the base editor SpRY-ABE8e^105^, 1 μl of 40 μM modified sgRNA (Synthego). Cells were nucleofected with an mRNA/sgRNA mixture in an Amaxa 4D nucleofector (SF protocol: CA-137). Cells were immediately transferred to 24-well plates with pre-warmed media and cultured. We performed nucleofection on day 0 and harvested cells for CRAFTseq on day 5. Editing efficiency was confirmed before CRAFTseq by Sanger-sequencing at Azenta. sgRNA and pegRNA used in this study are described in **Table S14** and **S15.**

### Daudi cell-line cultures and genomic editing for validation CRAFTseq experiment

Daudi cells were cultured in cRPMI as mentioned above. In the editing experiment for rs1883832, we used the same mRNA/sgRNA/pegRNA mixture and nucleofection settings as mentioned above. For this experiment, we performed only one editing on day 0 and harvested cells for CRAFTseq on day 5. In the editing experiment for ATG-KO and non-targeted control (NTC), we used 0.5 μl of mRNA (1 μg/μl) encoding the base editor SpRY-ABE8e, 1 μl of 40 μM modified sgRNA (Synthego). Cells were nucleofected and cultured as mentioned above. We performed nucleofection on day 0 and harvested cells for CRAFTseq on day 5. Editing efficiency was confirmed before CRAFTseq by Sanger-sequencing at Azenta. sgRNA and pegRNA used in this study are described in **Table S14** and **S15.**

### Daudi cell-line cultures and genomic editing for CRAFTseq experiment with CD40L stimulation

Daudi cells were cultured in cRPMI as mentioned above. In the editing experiment for rs1883832, we used the same mRNA/sgRNA/pegRNA mixture and nucleofection settings as mentioned above. For ATG-KO and NTC, the same mRNA/sgRNA mixture and nucleofection settings as mentioned above. Since editing efficiency was low in the prime-editing experiment (rs1883832), we performed nucleofection twice (day 0 and day 5), and cells were stimulated with CD40L (100 ng/ml; MEGACD40L, Enzo life science) on day 15. After 48h of CD40L stimulation, cells were harvested for CRAFTseq on day 17. For base-editing experiment (ATG-KO), we performed nucleofection on day 0, and cells were stimulated with CD40L on day 10. After 48h of CD40L stimulation, cells were harvested for CRAFTseq on day 12. Editing efficiency was confirmed before CRAFTseq by Sanger-sequencing at Azenta. sgRNA and pegRNA used in this study are described in **Table S14** and **S15.**

### Primary B cell cultures and genomic editing for CRAFTseq experiment

We used PBMC collected in the same way as in the previous study^23^. We recruited healthy individuals and processed 40–50 ml of peripheral blood under a Mass General Brigham (MGB) IRB-approved protocol (IRB# 2008P000427). PBMCs were isolated by layering Ficoll Paque (Sigma-Aldrich) underneath 1:1 PBS-diluted blood, followed by centrifugation. Buffy coat layers were extracted, washed in PBS, and then resuspended in cRPMI. Cells were stored by adding an equal volume of freezing medium (10% DMSO and 50% FBS in cRPMI) and frozen in liquid N_2_ until use.

To isolate B cells, frozen PBMCs were quickly thawed and put immediately in warm cRPMI media. Cells were washed twice, and B cells were isolated using a magnetic negative selection kit (Miltenyi Biotech, Pan B Cell Isolation Kit, human) according to the manufacturer’s protocols. After B cell isolation, cells were re-suspended with 500μl of StemMACS™ HSC Expansion Medium XF (human; Miltenyi biotech) supplemented with 5% of human serum (Sigma-Aldrich) and penicillin-streptomycin, stimulated with 4 μl of cross-linked CD40L from Human CD40-Ligand Multimer Kits (Miltenyi biotech), 1 μl of IL-4 (final concentration 50 IU/ml, Miltenyi biotech), and 1 μl of IL-21 (final concentration 40 ng/ml, Miltenyi biotech), and transferred to a 48-well plate and cultured. On day 3, we performed nucleofection for prime-editing or base editing. For prime-editing (C→T editing), 1.5 μl of mRNA (1 μg/μl) encoding the prime editor PE7, 1 μl of 50 μM modified nicking sgRNA (Synthego), and 1μl of 100 μM pegRNA (IDT) were used. For base editing (T→C editing and NTC), we used 0.5 μl of mRNA (1 μg/μl) encoding the base editor SpRY-ABE8e, and 1 μl of 40 μM modified sgRNA (Synthego). Cells were nucleofected with an mRNA/sgRNA(/pegRNA) mixture in an Amaxa 4D nucleofector (P3 protocol: EH-115). Cells were immediately transferred to 48-well plates containing pre-warmed B cell stimulation medium, as mentioned above, and cultured. On day 8 (5 days after nucleofection), cells were harvested for CRAFTseq experiments. Successful editing was confirmed before CRAFTseq by Sanger-sequencing at Azenta. sgRNA and pegRNA used in this study are described in **Table S14** and **S15.**

### Cell staining with indexing and oligo-conjugated antibodies for CRAFTseq

Cells were washed with PBS and stained with Zombie NIR (1:100 dilution in PBS) for 15 minutes on ice. In the fine-mapping experiment, we did not use Zombie NIR or other dead cell staining method following the original CRAFTseq protocol. Cells were washed and resuspended in a staining buffer composed of 0.5% BSA and 2 mM EDTA in PBS. For sample indexing, up to 500,000 cells were treated with Human TruStain FcX (BioLegend) for 10 minutes to block Fc receptors, followed by incubation with fluorophore-conjugated anti-CD45 antibodies for 25 minutes on ice. After indexing, the cells were washed and pooled. The pooled cell suspension was again Fc-blocked for 10 min and subsequently stained with the TotalSeq-A Human Universal Cocktail 1.0 (BioLegend) for 30 min on ice. For specific datasets, additional fluorophore-conjugated antibodies were included: anti-CD40-PE for the fine-mapping, validation, and primary B cell CRAFTseq, and anti-CD27-FITC for the primary B cell CRAFTseq. Finally, cells were washed and used for single-cell sorting with Invitrogen Bigfoot Spectral Cell Sorter. A comprehensive list of antibodies used for index sorting is provided in **Table S16**.

### CRAFTseq

We performed CRAFTseq based on the original protocol^23^ with slight modification. Lysis buffer containing 0.2% TritonX (Millipore Sigma), 1.2 U recombinant RNase inhibitor (Takara), 1.2 mM DTT, 1 M Betaine, 6 mM dNTP and 9 mM dCTP (ThermoFisher) was added into a 384-well PCR plate using a multi-channel pipette. An Agilent Bravo was used to add 2.5 μM of a well-specific OligoDT barcode into each well, then to distribute 1 μl of barcoded lysis buffer to multiple 384-well plates. These lysis plates were sealed and stored at −20 °C until use the next day for sorting. Single cells stained with fluorophore– and oligo-conjugated antibodies were sorted into lysis plates, which were kept sealed at −80 °C until use for a maximum of 4 weeks. For library generation, plates were thawed and incubated at 72 °C for 3 min, then kept on ice during further processing. RT-PCR mix (4 μl) containing 1.8 μM template switch oligo (Azenta), 0.8 M Betaine, 4.8 mM DTT, 9.7 mM MgCl2, 0.8 U RNase inhibitor, Maxima H Minus reverse transcriptase, 2× KAPA HiFi HotStart ReadyMix (Roche) was then added to each well using an Agilent Bravo. After first-strand synthesis (50 °C for 1 h and 85 °C for 5 min), a mixture of ADT primers (0.05 μM) and genomic DNA-specific primers (0.2 μM) were added using an I.DOT liquid handler (Dispendix) and amplified using the following program: 98 °C for 5 min then 21 cycles of 98 °C for 20 s, 65 °C for 20 s and 72 °C for 6 min, followed by 72 °C for 5 min and then hold at 4 °C.

After amplification, the samples were split for processing of cDNA, genomic DNA and ADT. For genomic DNA, using the Bravo, 1 μl of product per well was taken for further amplification of genomic DNA with nested primers containing a capture oligo with well-specific barcodes. The PCR mix contained 2× KAPA HiFi HotStart ReadyMix and 0.4 μM region-specific capture primer, 0.4 μM region-specific P7 primer and 0.4 μM capture oligo with well-specific barcode. The DNA was amplified using the following program: 98 °C for 5 min, followed by 25 cycles of 98 °C for 20 s, 65 °C for 20 s and 72 °C for 30 s, followed by 72 °C for 5 min, then hold at 10 °C. We then pooled 2 μl of each well containing nested genomic DNA using the Bravo and purified it with 1.2× solid-phase reverse immobilization (SPRI) beads. In brief, room-temperature SPRI beads were incubated at the given ratio by volume with PCR product for 10 min, then placed on a magnet for 5 min. Supernatant was removed and the beads were washed twice with 80% freshly made ethanol, dried for up to 15 min in air and resuspended in nuclease-free water. The resuspended beads were incubated at room temperature for 5 min, then placed back on the magnet for 5 min, and the supernatant was used for following steps.

For cDNA, 2 μl per well of the original amplification product was pooled using the Bravo and cleaned with 0.6× SPRI beads; the supernatant was saved for ADT sequencing. The cleaned cDNA product was treated with ExoI to remove primers (37 °C for 5 min, 85 °C for 5 min for ExoI denaturation) and cleaned again with 0.8× SPRI. cDNA concentrations were measured on a QuBit and 2 ng was used for tagmentation with a NexteraXT kit (Illumina) following the manufacturer’s directions in half-volume reactions (25 μl total). For ADT, saved supernatant from the 0.6× SPRI clean-up was re-cleaned with 1.4× SPRI (2× total) and ExoI was treated as above. The ADT product was quantified on a QuBit. For the final amplification of pooled samples, 2 ng of tagmented cDNA and 0.5 ng of genomic DNA and ADT samples underwent a final amplification step from the 3’ end with custom Illumina-compatible primers. The PCR mix contained 0.2 μM P7 and P5 custom oligos, 1× Q5 buffer, 0.5 μl Q5 High-Fidelity DNA Polymerase, 0.2 μM dNTP and water to a total volume of 25 μl. Amplification used the following programs: cDNA 72 °C for 3 min gap repair, 95 °C for 30 s, followed by 16 cycles of 95 °C for 10 s, 55 °C for 30 s and 72 °C for 30 s, followed by 72 °C for 5 min, then hold at 10 °C; genomic DNA and ADT 98 °C for 3 min, followed by 15 cycles of 98 °C for 15 s, 65 °C for 20 s and 72 °C for 45 s, followed by 72 °C for 10 min. All samples were re-cleaned using SPRI beads (0.8× for cDNA, 1.2× for gDNA and 1.6× for ADT) for sequencing. All primer sequences can be found in the original publication for CRAFTseq^23^ and **Table S17, S18, S19, S20, and S21**. The full protocol can be found in a more readable format deposited on protocols.io (https://www.protocols.io/view/craftseq-d7bk9ikw).

### Illumina sequencing

Sequencing was performed at the Genomics Platform at the Broad Institute. gDNA, ADT, and cDNA libraries were pooled at a 2:1:20 ratio based on nM concentration. Pooled libraries were sequenced on a NovaSeq X (26 cycles for R1, 151 cycles for R2) with a 10% PhiX spike-in.

### *in vitro* transcription of base editor and prime editor mRNAs

Base editor and prime editor mRNAs were generated by *in vitro* transcription using the HiScribe T7 High-Yield RNA synthesis kit (NEB, E2040S) by the method described previously^23^. Q5 High-Fidelity 2X Master Mix was used to PCR amplify template plasmids and install a functional T7 promoter and a 120-nucleotide polyadenine tail. Transcription reactions were set up with complete substitution of uracil by N1-methylpseudouridine (Trilink BioTechnologies, N-1080) and co-transcriptional 5’ capping using the CleanCap AG analogue (Trilink BioTechnologies, N-7113) to generate a 5’ Cap1 structure. mRNAs were purified using ethanol precipitation according to the kit’s instructions, dissolved in nuclease-free water and normalized to a concentration of 2 μg/μl using Nanodrop RNA quantification.

### Processing of the single-cell transcriptome and surface protein data from CRAFTseq

We follow the pipeline used in the original publication in the CRAFTseq with modifications. For cDNA sequencing data, reads were aligned to the human genome (GRCh38 2020-A) using STARsolo^106^ (v.2.7.6a). Then, cells were filtered on at least 500 (Daudi cells) or 1,000 (primary B cells) gene detection and fewer than 20% mitochondrial reads. Counts were log(CP10K + 1) normalized using Seurat R package.

For ADT sequencing data processing, Kallisto kite^107^ (v.0.27.3) was used to build an ADT reference transcriptome corresponding to the ADT panel and to align ADT reads. Then, cells were filtered on at least 30 (validation and CD40L-stimulation experiment with Daudi cells) or 50 (fine-mapping experiment with Daudi cells and primary B cell experiment) surface protein detection. Counts were CLR normalized (margin = 2) using Seurat R package.

### Processing of the single-cell genotyping data from CRAFTseq

gDNA reads were demultiplexed based on cellular barcodes. Then, amplicons were aligned to the amplicon reference using CRISPResso2^108^ (v.2.2.9). Allele usage statistics were calculated from CRISPResso2 outputs. We further extracted sequences encompassing the gRNA/pegRNA-binding sites and identified the precise location of the target allele within each read using the matchPattern function from the Biostrings R package (parameters: max.mismatch = 4 and with.indels = TRUE). From the resulting matches, only sequences showing an exact match to the expected alleles (either wild-type or the targeted variant) were categorized as ‘expected alleles (WT or variant)’, while all other variants were treated as unexpected alleles. As an exception, for the genotyping of SpD-KO conditions, sequences harboring both the intended KO mutation and additional bystander editing were collectively classified as KO alleles because alleles with the intended KO mutation will cause KO of CD40 regardless of the presence of adjacent bystander edits. We removed cells with fewer than 1,000 (fine-mapping experiment, Daudi cell experiment with CD40L-stimulation, primary B cell CRAFTseq [PE1]), 5,000 (primary B cell CRAFTseq [PE2 and ABE]), or 10,000 (validation CRAFTseq) total allele counts (expected allele + unexpected alleles). For genotyping of each editing, we collected cells from single editing condition and non-targeted control and removed cells with high frequency of the unexpected alleles (> Q3 + 3 × IQR). For SpD-KO in Daudi cell and T→C editing in primary B cell experiment, we observed cells with high ratio of the unexpected alleles, which inflated the IQR and inadvertently relaxed the filtering threshold. To address this, we implemented a pre-hard-filter to exclude any cells with an unexpected allele ratio greater than 0.2 before applying the aforementioned IQR-based filtering. Then, we calculated the ratio of WT allele and variant allele for each cell and applied min-max normalization separately for each plate to merge cells from different plates. Then, we applied k-means clustering using the kmeans function from the R stats package with k=3, assuming three distinct clusters corresponding to homozygous wild-type, heterozygous, and homozygous variant genotypes. Allelic dosages were subsequently assigned to each cluster based on its mean variant allele ratio.

### Clustering of the primary B cell CRAFTseq data

First, we retained only cells that passed the aforementioned quality control criteria for both cDNA and ADT modalities and were successfully assigned a genotype. We removed cells from non-targeted editing condition. We selected the top 1,000 HVGs, while excluding immunoglobulin genes (IGH, IGK, and IGL) and performed z-scaling and PCA. We applied the Harmony algorithm to correct for batch effects associated with both individuals (3 individuals) and plates (36 plates). The top 10 Harmony-corrected PCs were then utilized for Shared Nearest Neighbor graph construction and clustering with a resolution of 0.2 with seurat pipeline. To characterize each cluster, we performed differential expression analysis using the FindAllMarkers function to identify cluster-specific signatures. Based on the expression patterns of marker genes (**Fig. 4F**), we annotated four distinct B-cell states: activated naive, intermediate, plasmablast, and NR4A+ activated B cells. For visualization, RunUMAP function in Seurat R package was applied to the Harmony-corrected PCs.

### Modeling of gene and surface protein expression for genotype association analysis

To evaluate the associations between genetic variants and gene or surface protein expression, we employed negative binomial generalized linear modeling (GLM) using the R MASS package following the original CRAFTseq publication^23^. We modeled the raw counts of genes or ADT (*X_i_*) in each cell *i* as a function of the genotype dosage (0, 1, or 2) at the variant of interest. For analyses for evaluating the association between CD40 mRNA/ADT and genotypes, we modeled gene and ADT counts:

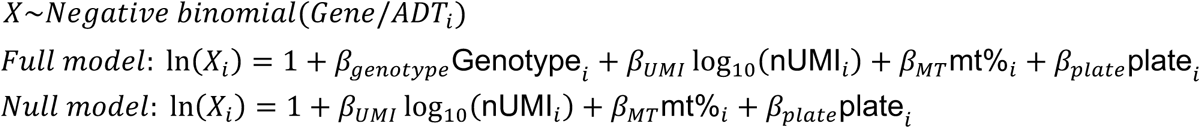

where gene/ADT counts are raw counts in each cell i, nUMI is total gene or ADT count in cell *i*, plate is plate identity, genotype is dosage at the edited nucleotide (0,1,2). For analyses for evaluating the association between CD40 mRNA/ADT and genotypes in primary B cells without clustering, we included top 2 PCs for gene expression to account for the heterogeneity of the cell states:

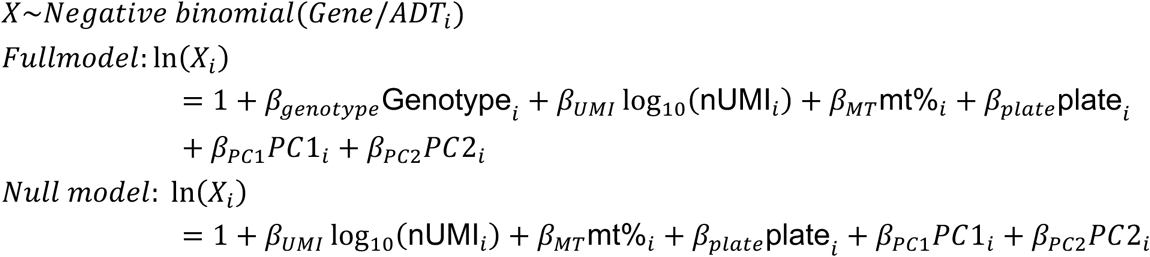

In primary B cell data analysis at cluster level, genes with mean UMI counts per cell > 1 and the percentage of cells with non-zero expression > 25% were analyzed for genome-wide analysis. For surface protein-wide analysis, features with mean UMI counts per cell > 1 and the percentage of cells with non-zero expression > 50% were analyzed. The association was tested using following model.

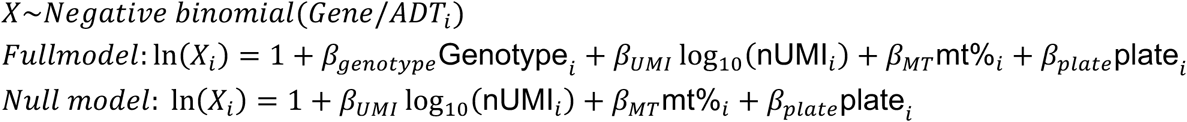

We did not include experiment batch as a covariate because it was redundant with plate identity (i.e., plate was nested within experiment batch). We used BH method to determine FDR. To ensure consistency across experimental batches, we also performed the regression analysis separately for each batch. For genes with *P* < 0.05 in each batch, the consistency of the effect size direction was validated using a one-sided sign test. For protein modalities, given the limited number of traits, we applied the sign test to all proteins without prior P-value filtering. In these tests, CD40 (exhibiting *cis*-effects) and CD154 (CD40L, which may capture the CD40L-multimer bound to CD40) were excluded. To evaluate differences in *trans*-effects between clusters, Pearson’s correlation coefficients of effect sizes were calculated for genes and surface proteins that reached FDR < 0.05 in activated naive B cells. Additionally, the correlation between effect sizes at the mRNA and surface protein levels was assessed. For evaluating the consistency of CRAFTseq effect sizes to the eQTL/spQTL meta-analysis of single-cell RNA-seq data, we performed a one-sided sign test on traits with FDR < 0.05 in activated B cells. We also calculated one-sided P-values for the *trans*-QTL sumstats based on the consistency of the effect direction to the CRAFTseq. CD40 and CD154 (CD40L) were consistently excluded from these statistical analyses.

In CD40L-stimulated Daudi cell analysis, genes with mean UMI counts per cell > 1 and the percentage of cells with non-zero expression > 25% were analyzed for genome-wide analysis. For surface protein-wide analysis, features with mean UMI counts per cell > 1 and the percentage of cells with non-zero expression > 50% were analyzed. The association was tested using a same model described above for each condition.

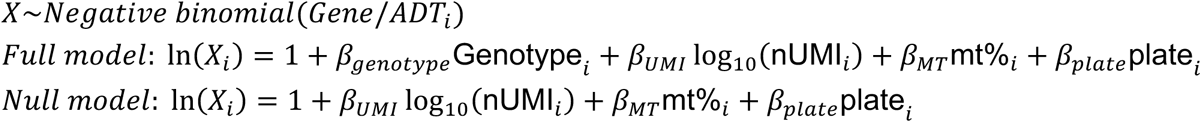

To account for the difference between the stimulated and non-stimulated conditions, we also performed sensitivity analysis with down-sampling on the stimulated condition so that the cell count for each genotype was comparable to the non-stimulated condition. We used BH method to determine FDR. For genes showing significant effects of the KO genotype under the CD40L-stimulated condition (FDR < 0.05), we evaluated the correlation of their effect sizes with those from primary B cell CRAFTseq (activated naive B cells) and 5’UTR genotype effects from Daudi cell CRAFTseq. The correlations were quantified using Spearman’s correlation coefficient (one-sided). In this analysis, CD40 and CD154 were excluded from calculation.

### Bulk RNA-seq for Daudi cells stimulated with CD40L

Daudi cells were cultured in cRPMI as mentioned above. Cells were stimulated with CD40L (100 ng/ml; MEGA-CD40L, Enzo life science) before being harvested for bulk RNA-seq. Total RNA was extracted from Daudi cells with Monarch Total RNA Miniprep Kit (New England Biolabs) according to the manufacture’s instruction.

Bulk mRNA-seq was performed at the Molecular Biology Core Facilities (MBCF) at Dana-Farber Cancer Institute. Libraries were prepared using Roche Kapa mRNA HyperPrep strand specific sample preparation kits from 200 ng of purified total RNA according to the manufacturer’s protocol on a Beckman Coulter Biomek i7. The finished dsDNA libraries were quantified by Qubit fluorometer and Agilent TapeStation 4200. Uniquely dual indexed libraries were pooled in an equimolar ratio and shallowly sequenced on an Illumina MiSeq to further evaluate library quality and pool balance. The final pool was sequenced on an Illumina NovaSeq X Plus targeting 40 million 150 bp read pairs per library at the Dana-Farber Cancer Institute Molecular Biology Core Facilities.

Sequencing reads were aligned to the human genome (GRCh38 2020-A) using STAR (v.2.7.6a). Then, genes with a count per million (CPM) ≥ 1 in ≥50% of samples were retained for differentially expressed gene (DEG) analysis. We used the DESeq2 package^109^ for DEG analysis, and genes with FDR < 0.05 were considered significant DEGs. For significant DEGs, we compared their effect sizes with those from CRAFTseq with primary B cells and Daudi cells by a one-sided test based on Spearman’s correlation. For these tests, we removed CD40 and CD154.

### Gene set enrichment analysis

We performed gene set enrichment analysis using the R package fgsea to identify biological pathways associated with genotype-dependent expression changes. All genes tested in the negative binomial models were included in the analysis. Gene ranks were defined by the Z-score obtained from the negative binomial models. For pathway definitions, we utilized the MSigDB Curated Gene Sets (C2) collection via the msigdbr package. Gene symbols were mapped to Entrez IDs using org.Hs.eg.db. Enrichment was tested for pathways containing between 15 and 500 genes.

### Reference mapping of B cell scRNA-seq data to the Tonsil atlas

We downloaded the scRNA-seq Seurat objects for germinal center B cells (GCBC), naive and memory B cells (NMBC), and plasma cells (PC) from the Tonsil Atlas Zenodo repository (https://doi.org/10.5281/zenodo.6340174)^61^. From these objects, only cells profiled via 3’ scRNA-seq in the discovery dataset were retained. We then identified the top 3000 highly variable genes (HVGs), excluded immunoglobulin-related genes from HVGs, and constructed a reference using the buildReference function in Symphony^62^ (d = 30) with gem_id designated as the batch term.

For query datasets, B cells were extracted from the Primary B cell CRAFTseq, AMP-RA PBMC, and AMP-RA synovium datasets. For AMP-RA synovium dataset, we used only cells from RA patients. Reference mapping was performed using the mapQuery function in Symphony, specifying plate, donor, and donor as the batch terms for the respective datasets. Finally, cell-type annotations from the Tonsil Atlas were transferred to each query cell using the knnPredict function in Symphony (k = 5).

### Bulk RNA-seq for primary B cells cultured with CD40L, IL-4, and IL-21

To isolate naive B cells, frozen PBMCs mentioned above were quickly thawed and put immediately in warm cRPMI media. Cells were washed twice, and B cells were isolated using a magnetic negative selection kit (Miltenyi Biotech, Naive B Cell Isolation Kit II, human) according to the manufacturer’s protocols. After naive B cell isolation, cells were re-suspended with 250μl of StemMACS™ HSC Expansion Medium XF (human; Miltenyi biotech) supplemented with 5% of human serum (Sigma-Aldrich) and penicillin-streptomycin, stimulated with 2μl of cross-linked CD40L from Human CD40-Ligand Multimer Kits (Miltenyi biotech), 0.5μl of IL-4 (final concentration 50 IU/ml, Miltenyi biotech), and 0.5μl of IL-21 (final concentration 40 ng/ml, Miltenyi biotech), and transferred to a 96-well round bottom plate and cultured for 4 h, 24 h, or 72 h. B cells before and after culturing are re-suspended with buffer RLT and kept in –80 ℃ until RNA extraction. Total RNA was extracted with RNeasy Micro Kit (QIAGEN) according to the manufacture’s instruction.

Bulk low input total RNA-seq was performed at the MBCF at Dana-Farber Cancer Institute. Libraries were prepared using SMARTer Stranded Total RNAseq v3 Pico Input Mammalian sample preparation kits from up to 2ng of purified total RNA according to the manufacturer’s protocol. Final dsDNA libraries were quantified using a Qubit fluorometer and fragment size distribution was assessed using the Agilent TapeStation 4200. Libraries were separately evaluated by shallow sequencing on an Illumina MiSeq for quality assessment. Libraries were subsequently sequenced on an Illumina NovaSeq X Plus targeting 50 million paired-end 150 bp reads at the Dana-Farber Cancer Institute Molecular Biology Core Facilities.

Sequencing reads were aligned to the human genome (GRCh38 2020-A) using STAR (v.2.7.6a). Then, genes with a CPM ≥ 1 in ≥50% of samples were retained for DEG analysis. We performed linear regression analysis using lm() function in R software with following formula separately for each time point: Gene expression [log(CPM+1)] ∼ genotype (T/T) + sex + log_10_(total read count) + RNA quality (RIN). Effect sizes in DEG analysis were compared to those from primary B cell CRAFTseq, as mentioned above. Also, one-sided P-values were calculated based on the consistency of the effect direction with the primary B cell CRAFTseq data.

For calculating the score for T-allele– and C-allele-associated genes in CRAFTseq, we performed linear regression analysis using lm() function in R software with following formula using all samples: Gene expression [log(CPM+1)] ∼ genotype (T/T) + time points + sex + log_10_(total read count) + RNA quality (RIN). Then, we adjusted the normalized gene expression for sex, log_10_(total read count), and RNA quality (RIN) using the coefficients from the linear regression analysis. We aggregated the adjusted gene expression level weighted by effect sizes from CRAFTseq.

### Mixed-effect association of single-cells for rs1883832

To test whether the rs1883832 variant is associated with B cell subtype proportions, we applied Mixed-effects model Association with Single Cells (MASC)^64^ across five independent cohorts: Perez et al (Asian/European), AMP-RA, AMP-SLE, and OneK1K. Samples contributing fewer than 5 total cells were excluded from all analyses. For each cell type, a logistic generalized linear mixed model (GLMM) was fit using the glmer function (lme4 package in R; nAGQ of 9 is specified) with a binomial outcome indicating whether each cell belonged to the target subtype. Allele dosage of rs1883832, age, sex, disease status (not for OneK1K), batch, and the top genetic principal components (gPCs; 4 for Perez cohorts; 5 for AMP-RA, AMP-SLE; 6 for OneK1K) were included as fixed effects. Results from all the cohorts were meta-analyzed with fixed-effect meta-analysis using metafor R package.

### Quantification and statistical analysis

Please refer to figure legends and method details for details of statistical analysis. Unless specified, statistical tests were conducted as two-sided. Number of the samples used in the analyses are described in **Table S1 and Table S6**. Throughout this study, the boxplot indicates the median values (center lines) and IQRs (box edges), with the whiskers extending to the most extreme points within the range between (lower quantile − [1.5 × IQR]) and (upper quantile + [1.5 × IQR]).

## Acknowledgements

We thank the International Multiple Sclerosis Genetics Consortium (IMSGC) for providing GWAS summary data. We thank members of the Raychaudhuri lab for helpful discussions.

## Author contribution

Conceptualization: Y.T., S.R.; Experiment: Y.T., Z.M., C.L., H.M., V.J.; Data analysis: Y.T., Z.M., Y.Z., N.S., J.A.S.; Resources: Y.T., Z.M., C.L., H.M., V.J., J.A.S.; Supervision: S.R.; Writing – original draft: Y.T., Z.M., H.M., Y.Z., S.R.; Writing – review & editing: All other authors; Funding acquisition: S.R.

AMP RA/SLE Network members contributed to this work by managing patient recruitment, curating clinical data, obtaining and processing synovial tissue samples, managing biorepositories, conducting histologic or computational analysis, providing software code, providing website support and/or providing input on data analysis and interpretation.

## Data, code, and materials availability

Raw sequencing data and processed data for CRAFTseq will be deposited upon publication of this manuscript. The AMP RA/SLE data (https://www.synapse.org/Synapse:syn26710600/datasets/) is available via Synapse under Data Usage Agreement. The custom scripts used in this manuscript are available at Zenodo^110^. The scRNA-seq and CITE-seq data from Perez et al ^33^ are available in the Human Cell Atlas Data Coordination Platform and at GEO accession number GSE174188. Genotypes from Perez et al are available at dbGap accession number phs002812.v1.p1. The tonsil scRNA-seq data is available on Zenodo ^111^.

**List of Supplementary**

**Materials: Figs. S1 to S20**

**Tables S1 to S21**

