## Supplementary Figure for "Identifying the causal allele in the CD40 autoimmune locus enables discovery of context-specific trans-effects in B cells"

Tomofuji Y et al.

Corresponding to Soumya Raychaudhuri

**Table of the contents**

**■Supplementary Figures........................................................................................ 1**
1. Overview for the datasets used for spQTL and eQTL mapping ........................... 2
2. Comparison of spQTL mapping methods ............................................................. 4
3. Comparison of spQTL mapping result across datasets........................................ 5
4. Cell-type specificity and cross-modality comparison of spQTLs........................... 6
5. Disease-associated spQTL signals to ITGA4 and PDCD1 surface proteins ......... 8
6. Disease-associated spQTL signals to the CD40 surface protein .......................... 10
7. Validation of successful CRISPR base- or prime-editing at the CD40 locus ......... 12
8. Fine-mapping of a causal CD40 spQTL variant using CRAFTseq in Daudi cells.. 13
9. Validation of causal CD40 spQTL variant using CRAFTseq in the Daudi cells..... 15
10. Quality controlling for the CRAFTseq with primary B cells ................................. 16
11. *cis*- and *trans*-effects of rs1883832 defined with primary B cell CRAFTseq ....... 18
12. Genes showing *trans*-effects from rs1883832 ................................................... 20
13. Bulk RNA-seq for Daudi cells stimulated with CD40L ........................................ 22
14. Quality controlling for the CRAFTseq with Daudi cells stimulated with CD40L ... 23
15. CRAFTseq for Daudi cells stimulated with CD40L ............................................ 25
16. CD40L-dependent *trans*-effects of rs1883832 in Daudi cells .............................. 26
17. Expression of CD40 downstream genes in Tonsil atlas ..................................... 28
18. Reference mapping of B cell single-cell RNA-seq datasets to the Tonsil atlas ... 30
19. Discordance between trans-effects identified from trans-QTL meta-analysis and CRAFTseq ........................................................................................................... 31
20. Consistency between bulk eQTL analysis and CRAFTseq of cultured primary B cells....................................................................................................................... 32

**■Contents of the supplementary tables ................................................................ 33**

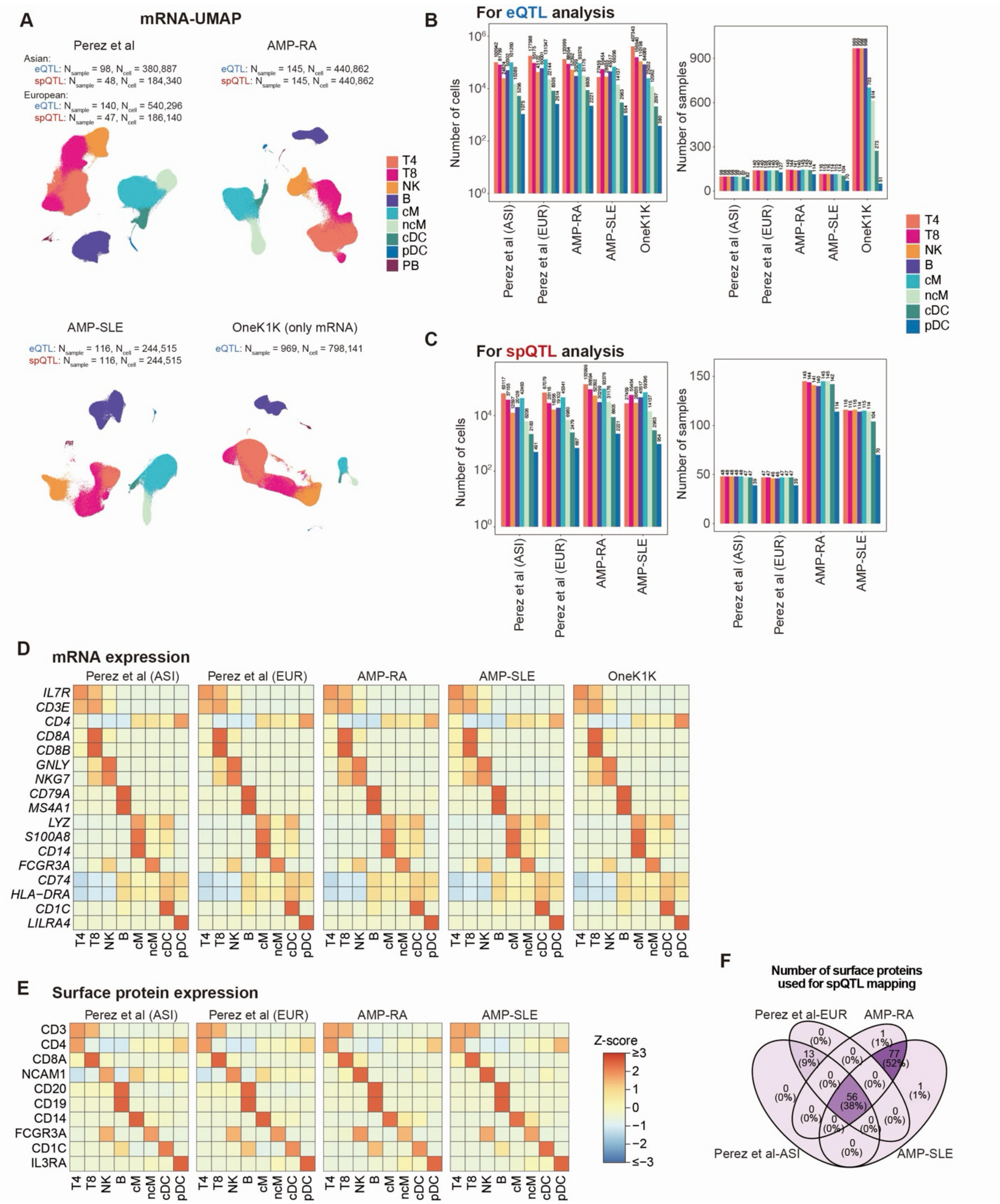


**Fig. S1. Overview for the datasets used for spQTL and eQTL mapping.** (**A**) UMAP generated from mRNA expression for each dataset. Dots are colored by cell annotation. (**B, C**) Bar plots showing the numbers of cells (left) and samples (right) used for spQTL (B) and eQTL (C) mapping, colored by cell annotation. (**D, E**) Heatmaps of marker gene expression across cell types in the mRNA (D) and surface protein (E) modalities. Color scales represent Z-normalized expression levels calculated from the mean of normalized

pseudobulk profiles across samples. (**F**) Venn diagram showing the numbers of surface proteins used for spQTL mapping in each dataset.
eQTL: expression quantitative trait loci; mRNA: messenger RNA; spQTL: surface protein quantitative trait loci; UMAP: uniform manifold approximation and projection.

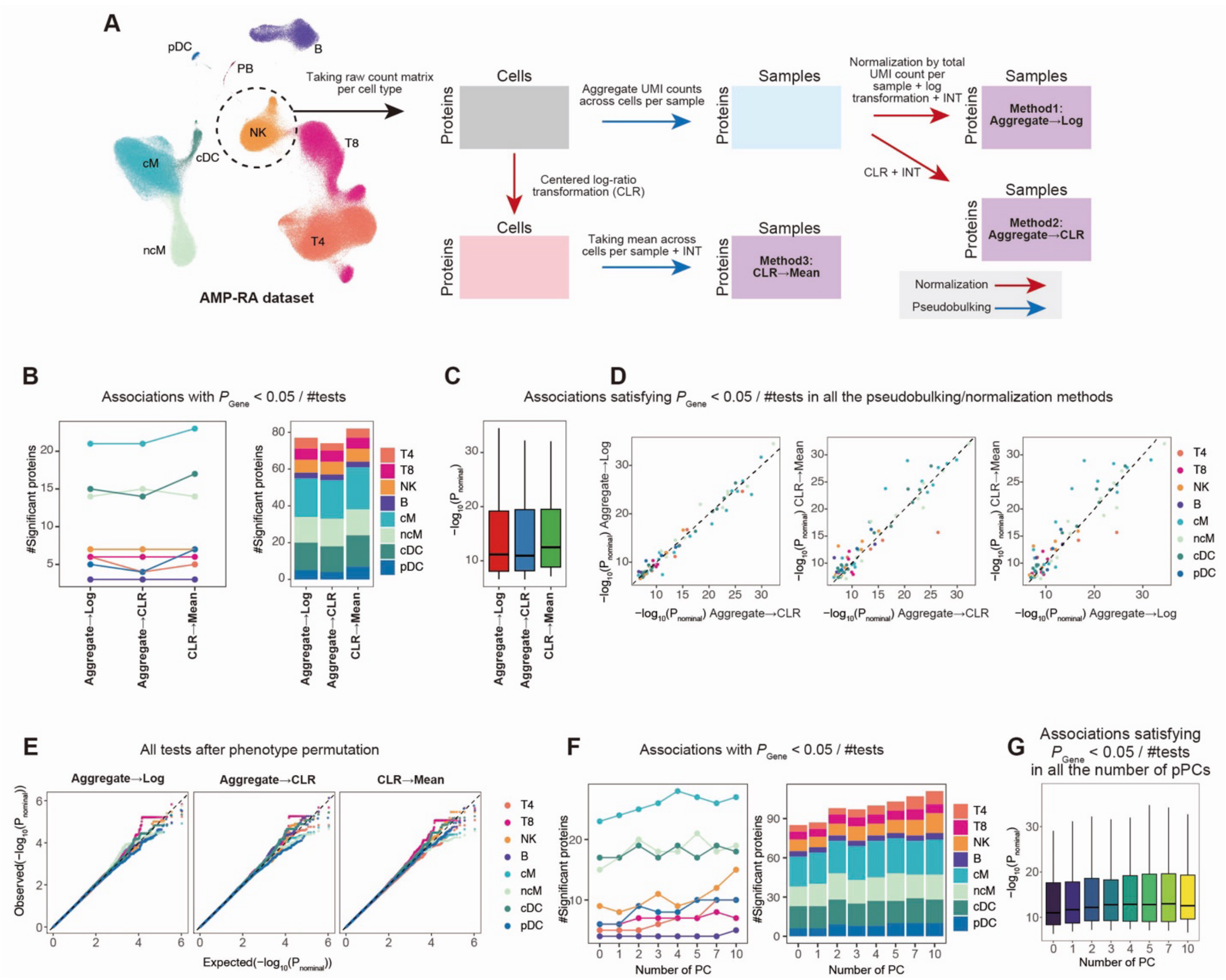


**Fig. S2. Comparison of spQTL mapping methods.** (**A**) Schematic overview of the three pseudobulking and normalization strategies compared in this analysis: Aggregate→Log, Aggregate→CLR, and CLR→Mean. (**B**) Numbers of significant spQTL signals detected by each pseudobulking and normalization strategy across cell types (left) and in total (right). Dots and bars are colored by cell annotation. (**C**) Box plots of P-values for lead variants, restricted to proteins consistently significant across all pseudobulking and normalization strategies. (**D**) Comparison of P-values for lead variants across pseudobulking and normalization strategies. Proteins were restricted to those consistently significant across all strategies. Dashed line indicates $x = y$. (**E**) Quantile–quantile plot of spQTL mapping results after permutation of protein expression across samples. Dashed line indicates $x = y$. (**F**) Numbers of significant spQTL signals detected using different numbers of protein PCs across cell types (left) and in total (right). (**G**) Box plots of P-values for lead variants across different numbers of protein PCs, restricted to proteins consistently significant across all PC settings.

CLR: centered log-ratio transformation; PC: principal component; spQTL: surface protein quantitative trait loci.

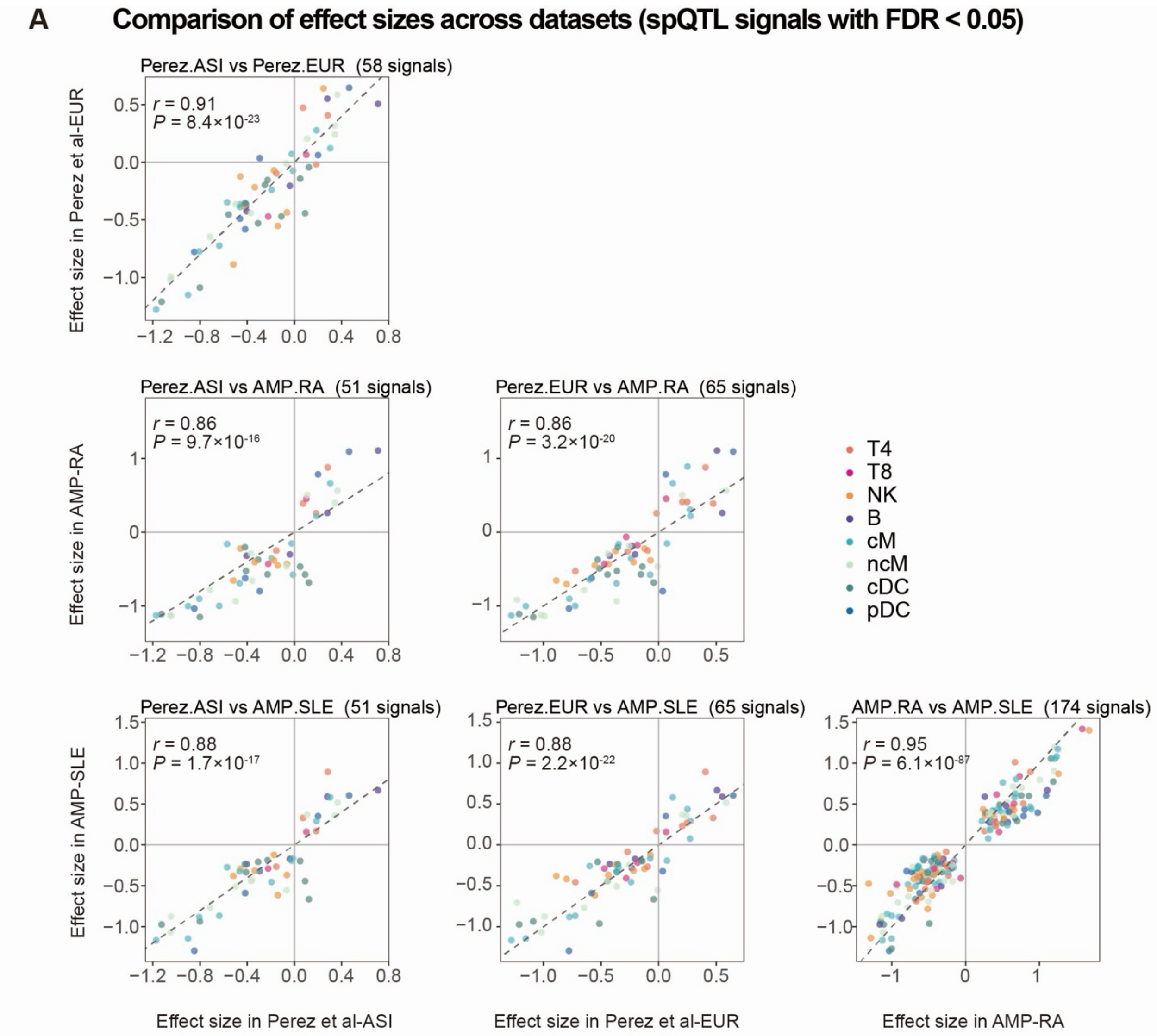


**Fig. S3. Comparison of spQTL mapping result across datasets.** (**A**) Comparison of spQTL effect sizes across datasets for significant signals identified by meta-analysis. r indicates Pearson's correlation coefficient. Dots are colored by cell annotation. Dashed line indicates *x* = *y.*
spQTL: surface protein quantitative trait loci.

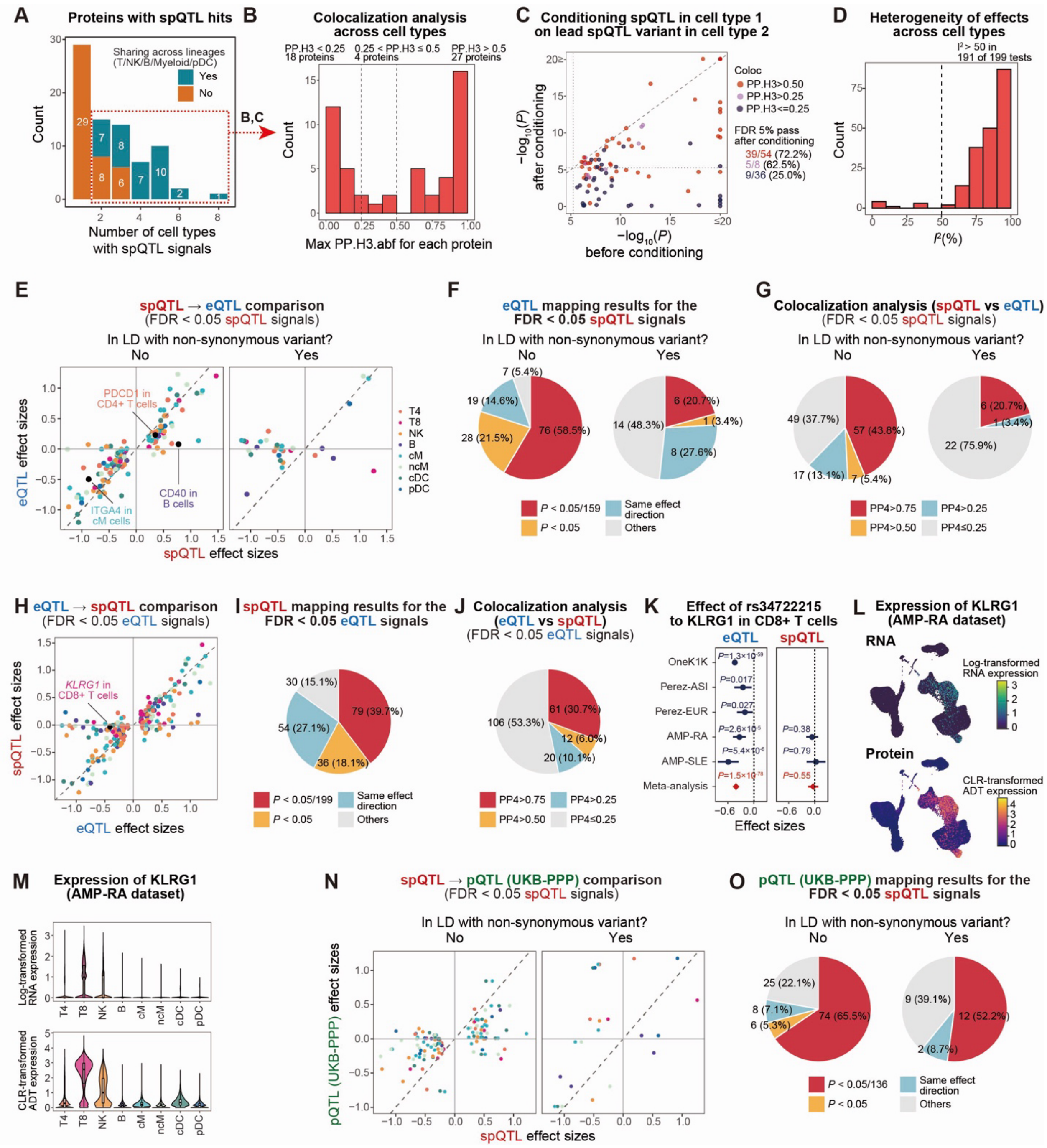


**Fig. S4. Cell-type specificity and cross-modality comparison of spQTLs.** (**A**) A histogram displays the number of cell types with spQTL signals for each protein. Bar colors show whether signals are shared across lineages (T: CD4+ and CD8+ T cells, Myeloid: classical monocytes, non-classical monocytes, conventional dendritic cells). (**B**) A histogram displays the posterior probabilities for the scenario in which two cell types have distinct causal spQTL signals (PP.H3) for each protein identified with spQTLs in at least two cell types. For each protein, cell type pairs with the highest PP.H3 are shown. Vertical dashed lines indicate $x$ = 0.25 and 0.5 (**C**) Conditional analysis of the spQTL signals between cell types. For the cell type pairs shown in (B), spQTL signals in cell type

1 are conditioned on the lead spQTL variant in cell type 2, and vice versa. -$\log_{10}$(P-value) before (x-axis) and after (y-axis) conditioning is shown. Colors of the dots indicate coloc PP.H3 values. Dashed lines indicate $x = y$ and the FDR 5% threshold before and after conditioning. (**D**) Histogram for the heterogeneity of the spQTL effects ($I^2$) across cell types for proteins with at least 1 spQTL signal. Vertical line indicates $x = 0.5$. (**E**) Comparison of the spQTL (x-axis) and eQTL (y-axis) effect size for the lead variant of significant spQTLs. Dots are colored by cell type. Genes highlighted in the fig. 2. are colored with black and labeled. The diagonal dashed line indicates $y = x$. spQTL signals are stratified based on whether lead variants are in LD ($R^2 > 0.8$ in any major 1000 Genomes Project population) with non-synonymous variants (right) or not (left). (**F,G**) Pie charts showing the distribution of eQTL P-values for the lead spQTL variants (F) and the posterior probability of a shared causal variant between eQTL and spQTL for significant spQTL loci (coloc PP.H4; G). spQTL signals are stratified based on whether lead variants are in LD with non-synonymous variants (right) or not (left). (**H**) Comparison of the eQTL (x-axis) and spQTL (y-axis) effect size for the lead variant of significant eQTLs. Dots are colored by cell type. *KLRG1* gene in CD8+ T cells highlighted in the Fig. 1 are colored with black and labeled. The diagonal dashed line indicates $y = x$. (**I,J**) Pie charts showing the distribution of spQTL P-values for the lead eQTL variants (I) and the posterior probability of a shared causal variant between eQTL and spQTL for significant eQTL loci (coloc PP.H4; J). (**K**) Forest plots showing the eQTL (left) and spQTL (right) signals of rs34722215 to KLRG1 in CD8+ T cells for each dataset. Error bars indicate 95% CI, and the dashed bar indicates $x = 0$. (**L,M**) UMAPs (L) and violin plots (M) showing the expression of KLRG1 mRNA (top) and surface protein (bottom) in the AMP-RA dataset. (**N**) Comparison of effect sizes between spQTLs (x-axis) and plasma pQTLs from the UKB-PPP study (y-axis) for the lead variants of significant spQTLs. Dots are colored by cell type. The diagonal dashed line indicates $y = x$. spQTL signals are stratified based on whether lead variants are in LD with non-synonymous variants (right) or not (left). (**O**) Pie charts showing the distribution of plasma pQTL P-values from the UKB-PPP study for the lead variants of significant spQTLs. spQTL signals are stratified based on whether lead variants are in LD with non-synonymous variants (right) or not (left).

CI: confidence interval; eQTL: expression quantitative trait loci; LD: linkage disequilibrium; mRNA: messenger RNA; PP.H3: posterior probability of a shared causal variant between cell types; PP.H4: posterior probability of a shared causal variant between eQTL and spQTL; spQTL: surface protein quantitative trait loci; UMAP: uniform manifold approximation and projection.

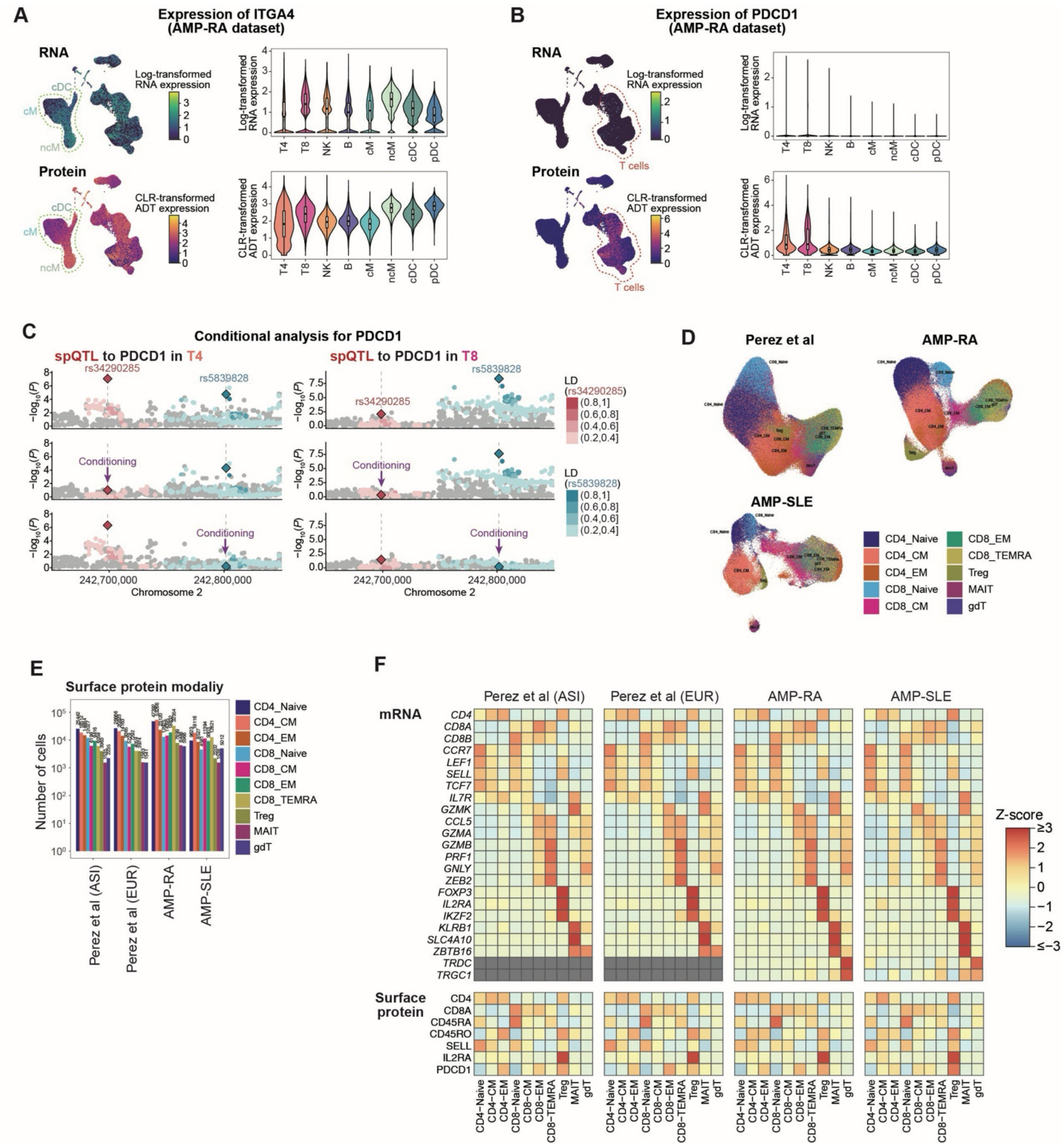


**Fig. S5. Disease-associated spQTL signals to ITGA4 and PDCD1 surface proteins.** (**A,B**) UMAPs (left) and violin plots (right) showing the expression of ITGA4 (A) or PDCD1 (B) mRNA (top) and surface protein (bottom) in the AMP-RA dataset. (**C**) Locus plot showing PDCD1 spQTL signals before conditioning (top), after conditioning on rs34290285 (middle), or after conditioning on rs5839828 (bottom) in CD4$^+$ T cells (left) and CD8+ T cells (right). Dots are colored according to the linkage disequilibrium (LD; $R^2$) from the lead variants in CD4 T cells (red; rs34290285) and CD8 T cells (blue; rs5839828) calculated with 1000 Genome project reference panel ($N$ = 2,504). (**D**) UMAPs generated from the mRNA expression of T cells (CD4+ and CD8+ T cells) for each dataset. Dots are colored according to cell annotations assigned by the TCAT software. (**E**) Bar plots showing the numbers of cells with surface protein information, colored by T cell annotation. (**F**) Heatmaps of marker gene expression across T cell annotations in the mRNA (top) and surface protein (bottom) modalities. Color scales represent Z-normalized expression levels calculated from the mean of normalized single-cell-level expression for each T cell annotation.

LD: linkage disequilibrium; mRNA: messenger RNA; spQTL: surface protein quantitative trait loci; UMAPs: uniform manifold approximation and projections.

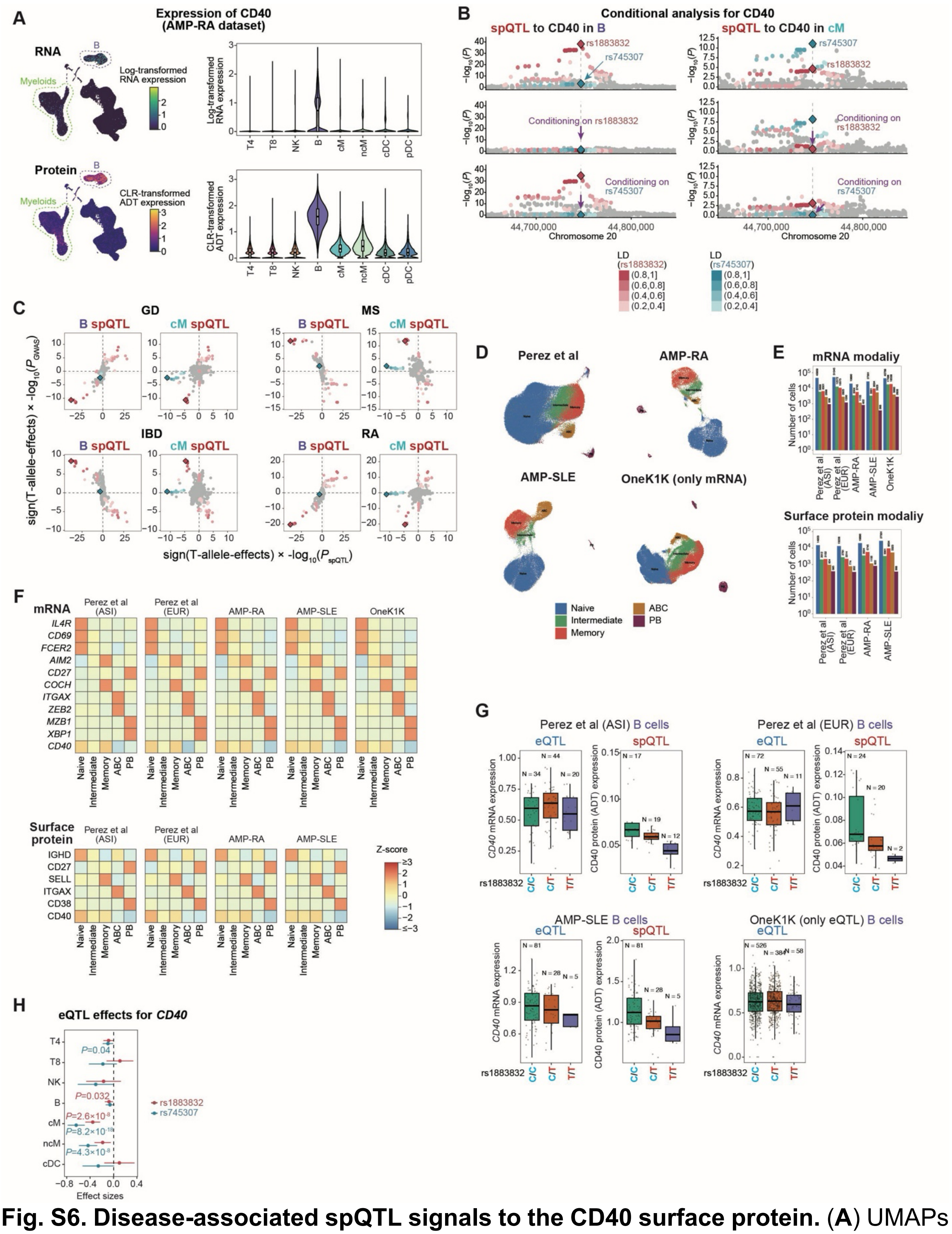

**Fig. S6. Disease-associated spQTL signals to the CD40 surface protein.** (**A**) UMAPs (left) and violin plots (right) showing the expression of CD40 mRNA (top) and surface protein (bottom) in the AMP-RA dataset. (**B**) Locus plot showing CD40 spQTL signals before conditioning (top), after conditioning on rs1883832 (middle), or after conditioning

on rs745307 (bottom) in B cells (left) and classical monocytes (right). Dots are colored according to the LD ($R^2$) from the lead variants in B cells (red; rs1883832) and classical monocytes (blue; rs745307) calculated with 1000 Genome project reference panel ($N$ = 2,504). (**C**) Comparison of genetic effects between disease GWAS and spQTLs in B cells (left) or classical monocytes (right). The scatter plots show signed $-\log_{10}$(P-values) for spQTLs (x-axis) and disease GWAS (y-axis). Dots are colored according to the LD ($R^2$) from the lead variants in B cells (red; rs1883832) and classical monocytes (blue; rs745307) calculated with 1000 Genome project reference panel ($N$ = 2,504). (**D**) UMAPs generated from the mRNA expression of B lineage cells (B cells and plasma cells) for each dataset. Dots are colored according to cell annotations. (**E**) Bar plots showing the numbers of cells with mRNA (top) and surface protein (bottom) information, colored by B cell annotation. (**F**) Heatmaps of marker gene expression across B cell annotations in the mRNA (top) and surface protein (bottom) modalities. Color scales represent Z-normalized expression levels calculated from the mean of normalized single-cell-level expression for each B cell annotation. (**G**) Boxplot showing the expression of CD40 mRNA (left) and surface protein (right) across rs1883832 genotypes in Perez et al, AMP-SLE, and OneK1K datasets. (**H**) Forest plots showing the eQTL signals of rs1883832 (red) and rs745307 (blue) to CD40 for each major cell type. Error bars indicate 95% CI, and the dashed line indicates $x$ = 0. Although the T allele is the reference allele in the hg19 human reference genome, we show the effect size of the T allele because T is a minor allele in the population and the edited allele in most of the following editing experiments. P-values below 0.05 are described.
CI: confidence interval; eQTL: expression quantitative trait loci; LD: linkage disequilibrium; mRNA: messenger RNA; spQTL: surface protein quantitative trait loci; UMAPs: uniform manifold approximation and projections.

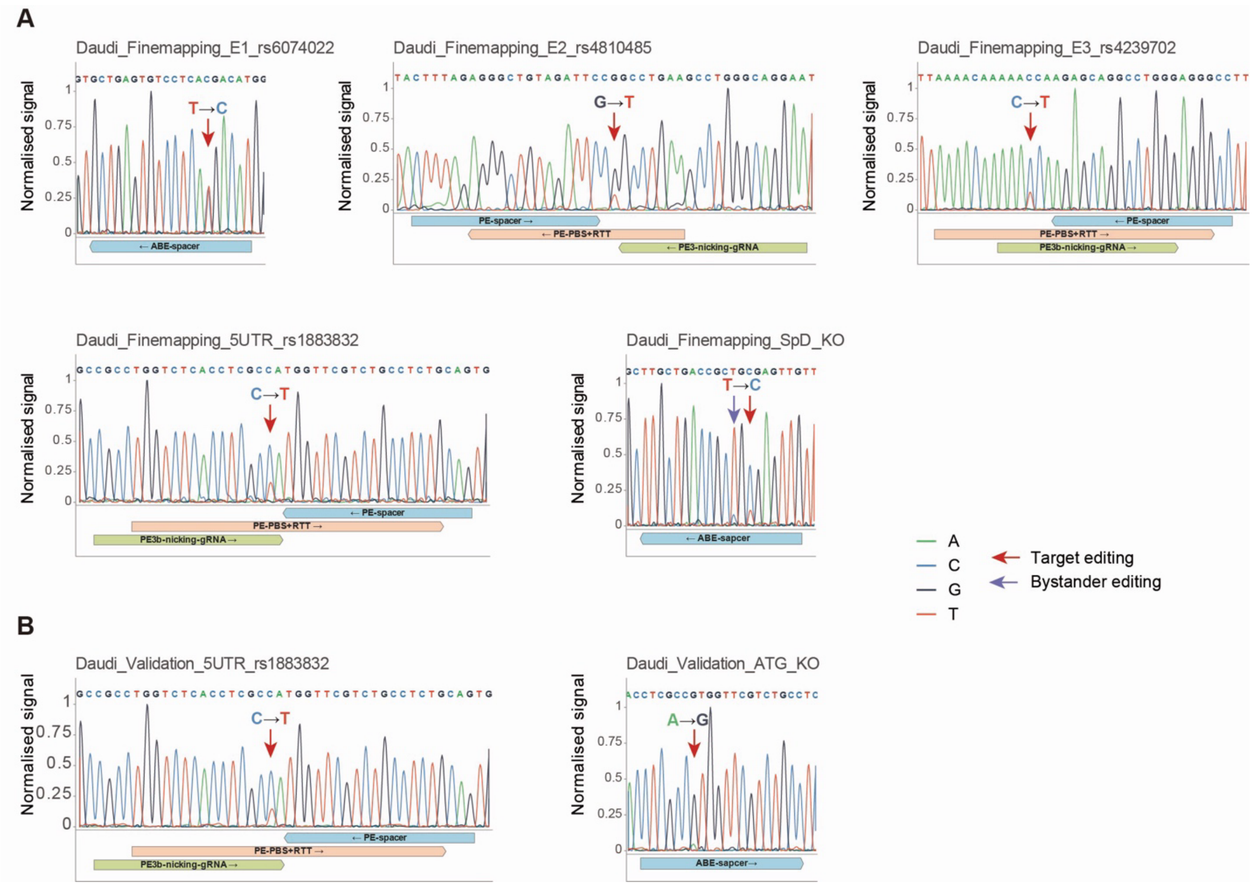


**Fig. S7. Validation of successful CRISPR base- or prime-editing at the CD40 locus.** (**A, B**) Sanger sequencing chromatograms showing the genomic DNA of Daudi cells after CRISPR editing in fine-mapping (A) and validation (B) CRAFTseq experiment. The target site is indicated by the red arrow, and the bystander editing site is indicated by the purple arrow. The ribbons indicate the sgRNA or pegRNA sequences, and the corresponding called bases are shown above the chromatograms.

pegRNA: prime editing guide RNA; sgRNA: single guide RNA.

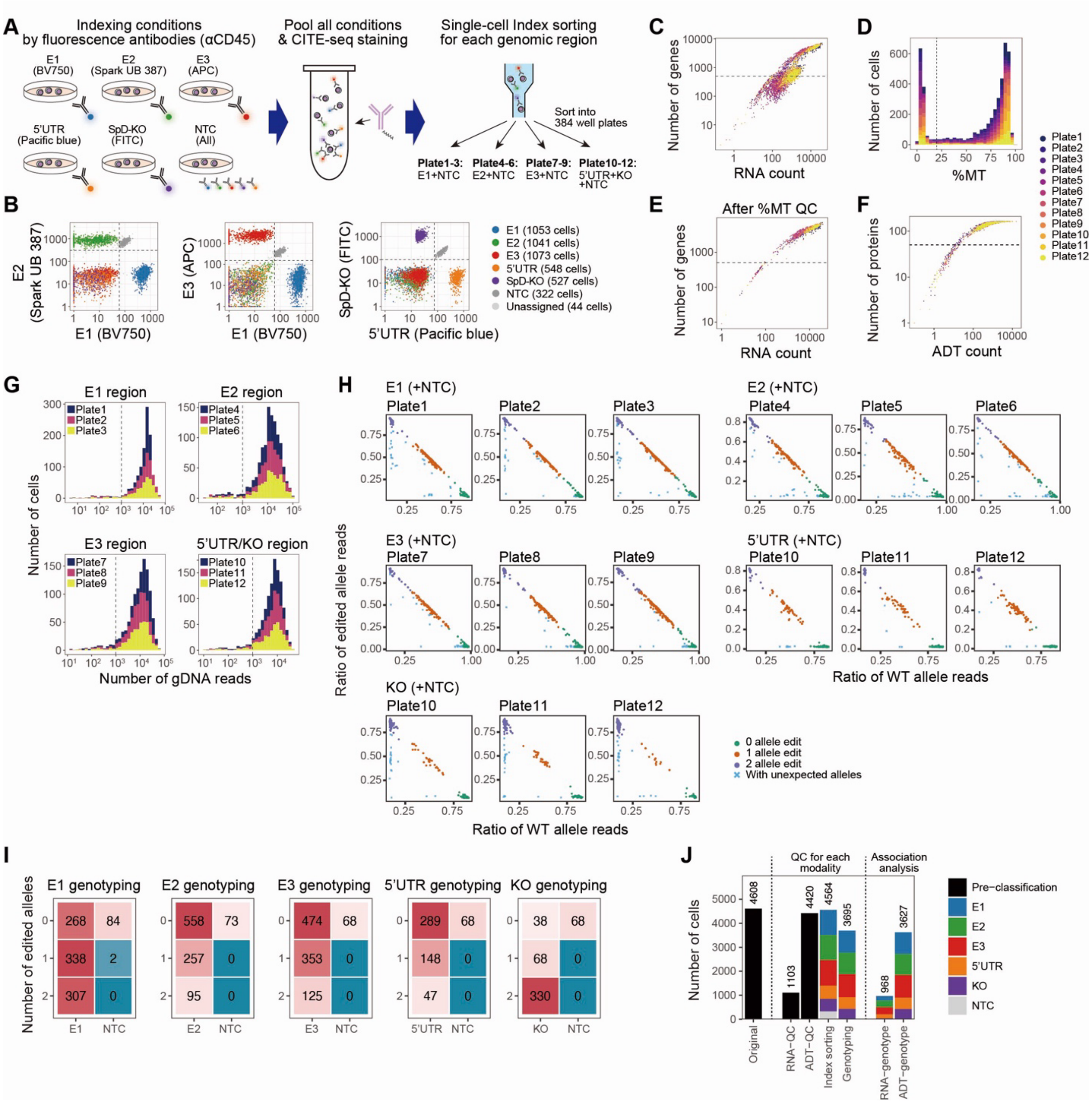


**Fig. S8. Fine-mapping of a causal CD40 spQTL variant using CRAFTseq in Daudi cells.**

(**A**) Schematic overview of the experimental design. Cells from each condition are labeled with unique combinations of fluorescent markers. After pooling all cells, they are sorted into separate plates for targeted genomic regions. The 5' UTR and SpD-KO conditions can be genotyped using the same primer pairs and sorted into the same plates. (**B**) Indexing flow cytometry data with gating for defining each condition. (**C**) A scatter plot showing the number of UMI for mRNA (x-axis) and the number of detected genes (y-axis). Dots are colored by plates. A dashed line indicates the number of detected genes = 500. (**D**) A histogram showing the percentage of mitochondrial gene expression for each cell. Bars are colored by plates. A dashed line indicates a percentage of mitochondrial gene expression = 20. (**E**) A scatter plot showing the number of UMIs for mRNA (x-axis) and the number of detected genes (y-axis) after removing cells with high mitochondrial gene expression. Dots are colored by plates. A dashed line indicates the number of detected genes = 500. (**F**) A scatter plot showing the number of UMIs for protein (x-axis) and the number of detected proteins (y-axis). Dots are colored by plates. A dashed line indicates the number of detected proteins = 50. (**G**) Histograms showing the number of gDNA reads from single cells for each genomic region. Bars are colored by plates. A dashed line indicates a number of gDNA reads = 1,000. (**H**) Scatter plots showing the ratio of WT alleles (x-axis) and edited alleles (y-axis) for each plate. Dots are colored by genotyping results. (**I**) Heatmaps showing the genotyping results for each variant. (**J**) A bar plot showing the number of cells at each quality control step. Bars are colored according to the condition.

gDNA: genomic DNA; mRNA: messenger RNA; SpD-KO: splicing donor knockout; UMI: unique molecular identifier; UTR: untranslated region; WT: wild-type.

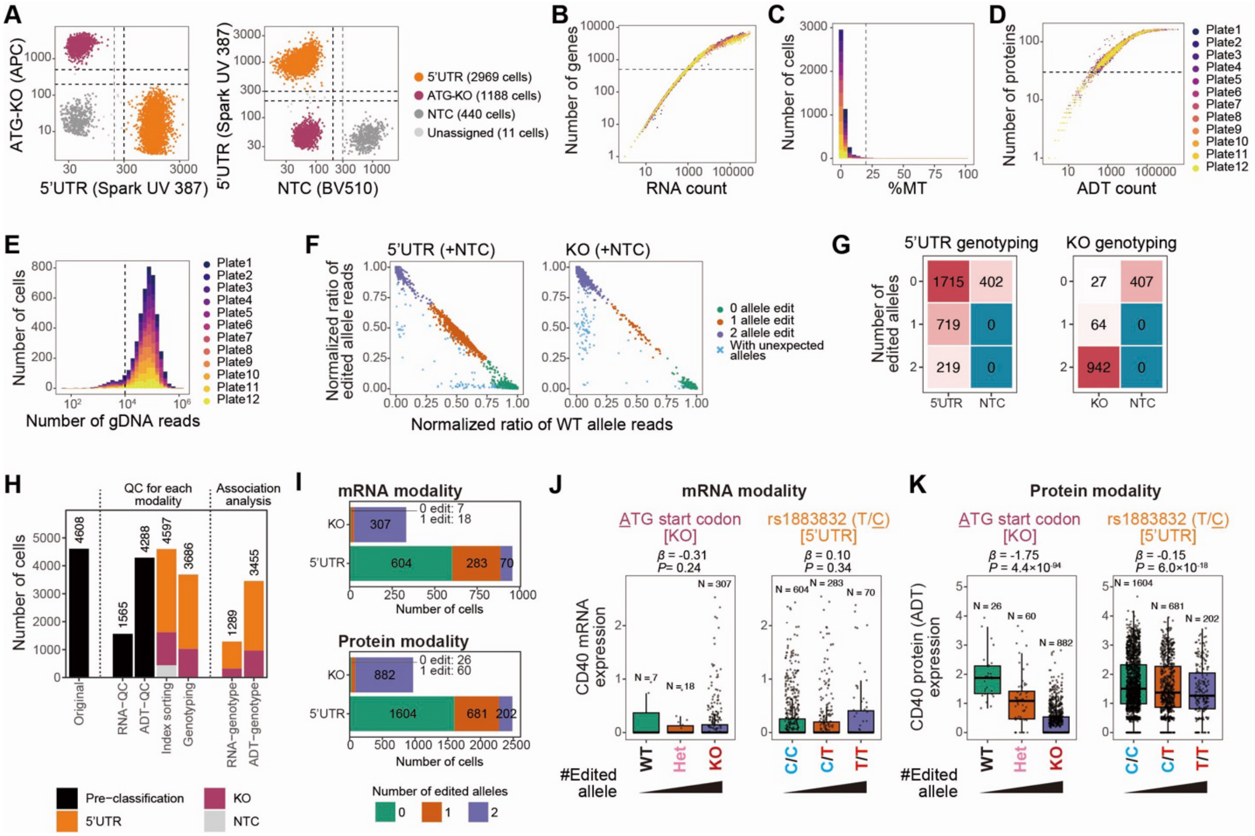


**Fig. S9. Validation of causal CD40 spQTL variant using CRAFTseq in the Daudi cells.** (**A**) Indexing flow cytometry data with gating for defining each condition. (**B**) A scatter plot showing the number of UMI for mRNA (x-axis) and the number of detected genes (y-axis). Dots are colored by plates. A dashed line indicates the number of detected genes = 500. (**C**) A histogram showing the percentage of mitochondrial gene expression for each cell. Bars are colored by plates. A dashed line indicates a percentage of mitochondrial gene expression = 20. (**D**) A scatter plot showing the number of UMIs for protein (x-axis) and the number of detected proteins (y-axis). Dots are colored by plates. A dashed line indicates the number of detected proteins = 30. (**E**) A histogram showing the number of gDNA reads from single cells. Bars are colored by plates. A dashed line indicates a number of gDNA reads = 10,000. (**F**) Scatter plots showing the normalized ratio of wild-type (WT) alleles (x-axis) and edited alleles (y-axis) for cells from all plates. Dots are colored by genotyping results. (**G**) Heatmaps showing the genotyping results for each variant. (**H**) A bar plot showing the number of cells at each quality control step. Bars are colored according to the condition. (**I**) Bar plots showing the cell counts for each edited genomic position used in mRNA-genotype association analysis (top) and surface protein-genotype association analysis (bottom). Bars are colored according to the number of edited alleles. (**J,K**) Boxplot showing the expression of CD40 mRNA (J) and surface protein (K) across different genotypes of start codon (left) and rs1883832 (right).

gDNA: genomic DNA; mRNA: messenger RNA; UMI: unique molecular identifier; WT: wild-type.

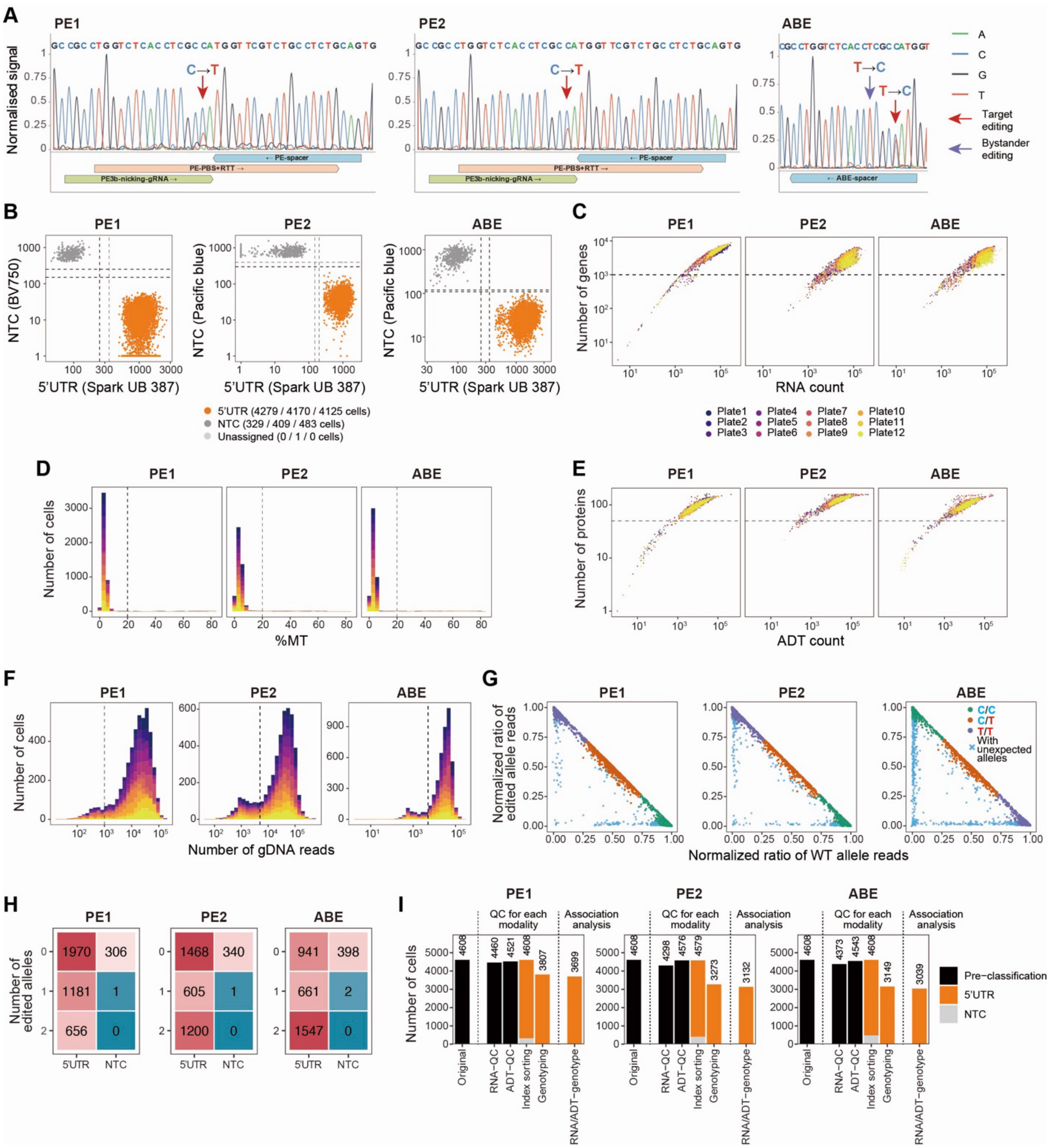


**Fig. S10. Quality controlling for the CRAFTseq with primary B cells. (A)** Sanger sequencing chromatograms showing the genomic DNA of primary B cells after CRISPR editing. The target site is indicated by the red arrow, and the bystander editing site is indicated by the purple arrow. The ribbons indicate the sgRNA or pegRNA sequences, and the corresponding called bases are shown above the chromatograms. (**B**) Indexing flow cytometry data with gating for defining each condition. Results from three independent experiments are shown separately. (**C**) A scatter plot showing the number of UMI for mRNA (x-axis) and the number of detected genes (y-axis) for each experiment. Dots are colored by plates. A dashed line indicates the number of detected genes = 1,000.

(**D**) A histogram showing the percentage of mitochondrial gene expression for each cell in each experiment. Bars are colored by plates. A dashed line indicates a percentage of mitochondrial gene expression = 20. (**E**) A scatter plot showing the number of UMIs for protein (x-axis) and the number of detected proteins (y-axis) for each experiment. Dots are colored by plates. A dashed line indicates the number of detected proteins = 50. (**F**) A histogram showing the number of gDNA reads from single cells for each experiment. Bars are colored by plates. A dashed lines indicate number of gDNA reads = 1,000 (PE1) or 5,000 (PE2 and ABE). (**G**) Scatter plots showing the normalized ratio of WT alleles (x-axis) and edited alleles (y-axis) for cells from all plates. Results from three independent experiments are shown separately. Dots are colored by genotyping results. (**H**) Heatmaps showing the genotyping results for each variant. (**I**) A bar plot showing the number of cells at each quality control step for each experiment. Bars are colored according to the condition.

ABE: adenine base editing; gDNA: genomic DNA; mRNA: messenger RNA; PE: prime editing; pegRNA: prime editing guide RNA; sgRNA: single guide RNA; UMI: unique molecular identifier; WT: wild-type.

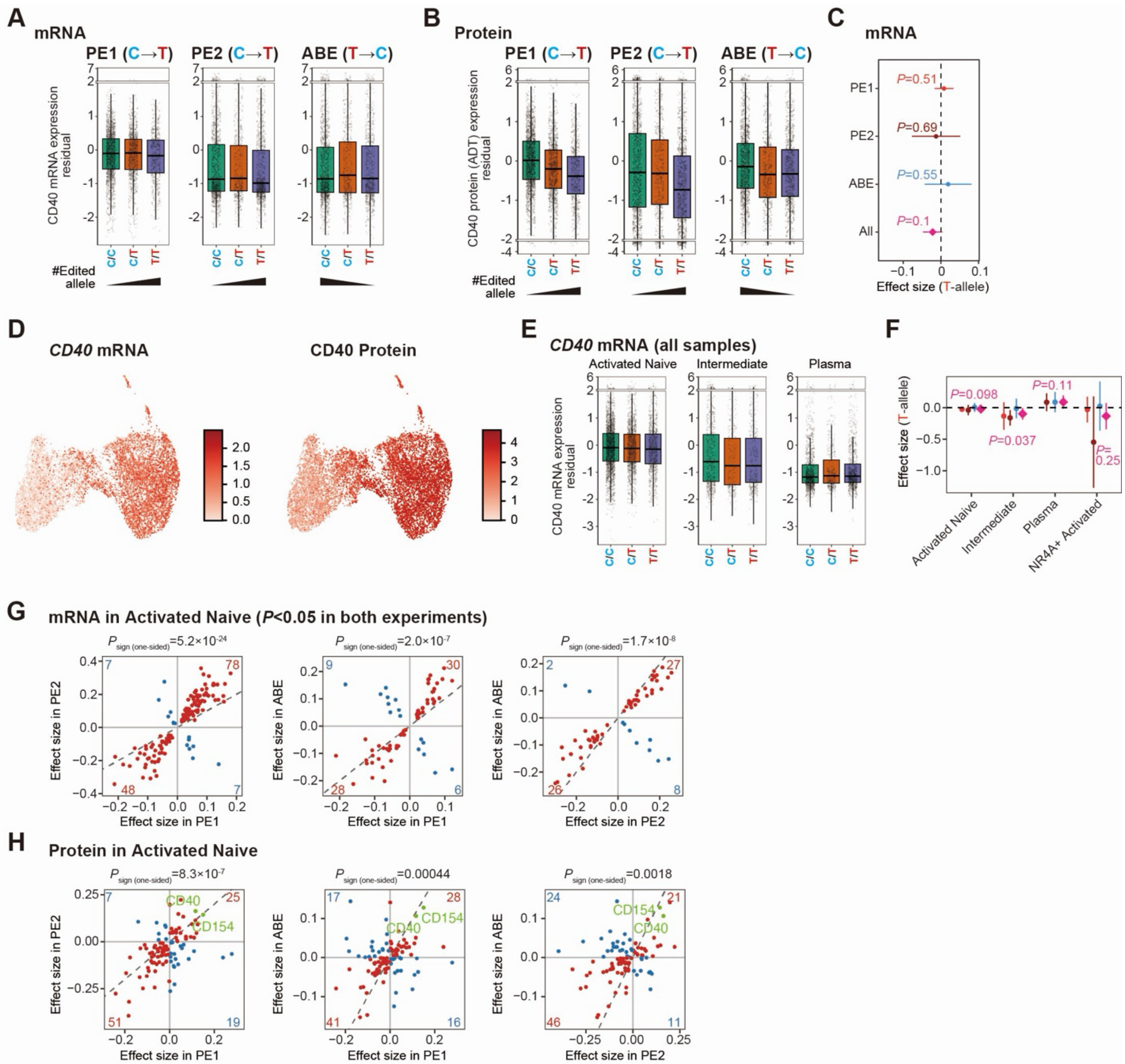


**Fig. S11. *cis*- and *trans*-effects of rs1883832 defined with primary B cell CRAFTseq.** (**A,B**) Boxplots showing CD40 mRNA (A) and (B) surface protein expression residualized for covariates including top 2 expression principal components across rs1883832 genotypes in each experiment. The y-axis includes scale breaks. (**C**) Forest plots showing the association between the expression of CD40 mRNA and rs1883832 genotypes in all the cultured B cells, for each individual experiment and the combined analysis. Error bars indicate 95% CI, and the dashed bar indicates $x = 0$. (**D**) UMAPs showing the expression of *CD40* mRNA (left) and surface protein (right) in the CRAFTseq. (**E**) Boxplots showing *CD40* mRNA expression residualized for covariates across rs1883832 genotypes in different cell states. The y-axis includes scale breaks. (**F**) Forest plots showing the association between *CD40* mRNA expression and rs1883832 genotypes across cultured B-cell subsets, for each individual experiment and the combined analysis. Error bars indicate 95% CI, and the dashed bar indicates $y = 0$. (**G**) Comparison of the CRAFTseq effect sizes across experiment in activated naive B cells at mRNA modality. Genes with $P < 0.05$ in both experiments are shown. Dots are colored based on the consistency of

the effect directions (red, same direction; blue, opposite direction). Consistency of the effect direction is tested with a one-sided sign test. Dashed lines indicate *x* = *y*. (**H**) Comparison of the CRAFTseq effect sizes across experiment in activated naive B cells at surface protein modality. Proteins which were evaluated in both experiments are shown. Dots are colored based on the consistency of the effect directions (red, same direction; blue, opposite direction). Consistency of the effect direction is tested with a one-sided sign test. Dashed lines indicate *x* = *y*.
CI: confidence interval; FDR: false discovery rate; mRNA: messenger RNA; pQTL: protein quantitative trait loci; UMAPs: uniform manifold approximation and projections.

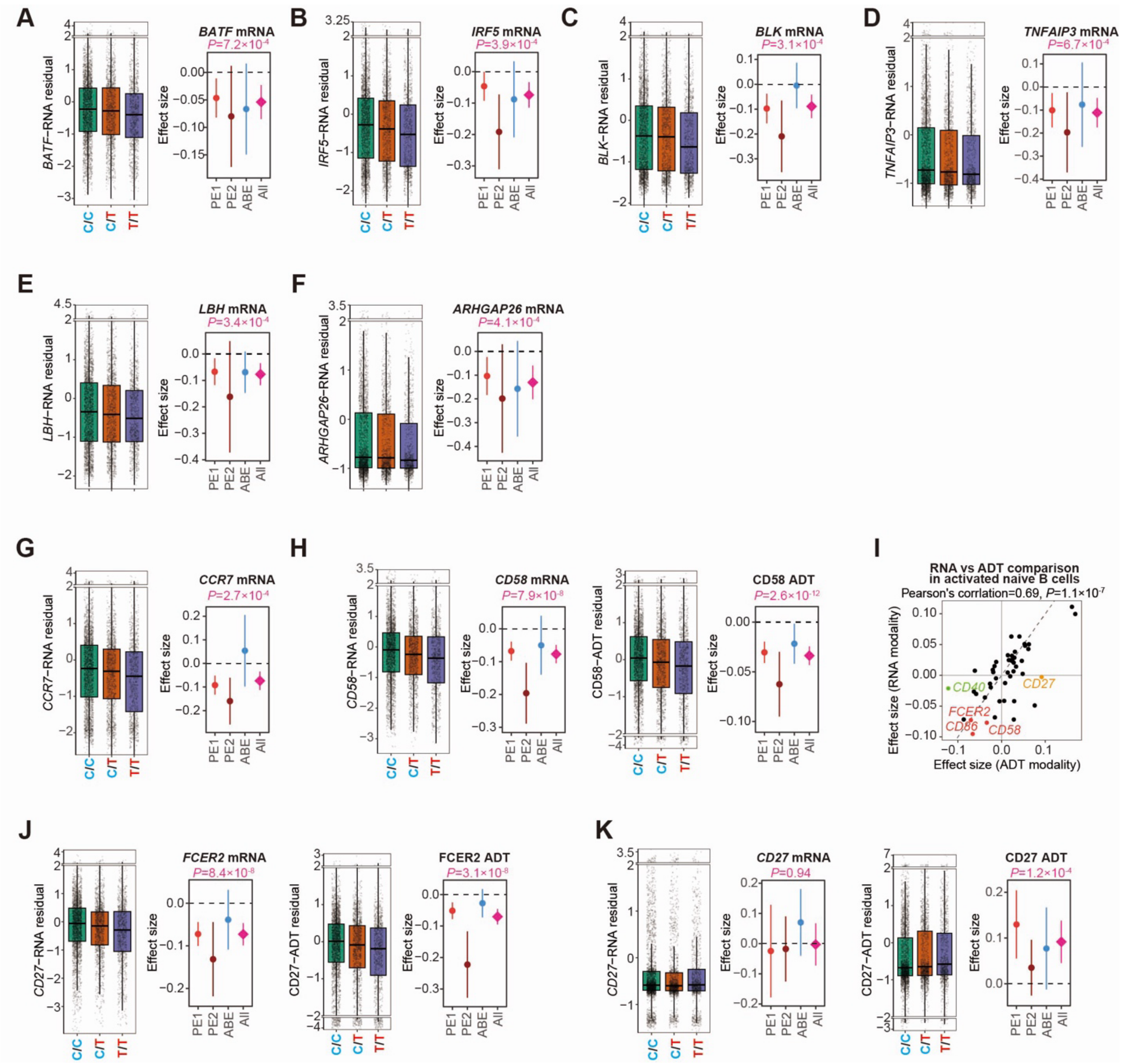


**Fig. S12. Genes showing *trans*-effects from rs1883832.**
(**A-H**) Boxplots showing mRNA or surface protein expression of RA-GWAS genes (A-F) and genes involved in B cell migration (G,H) residualized for covariates across rs1883832 genotypes in activated naive B cells. The y-axis includes scale breaks. The horizontal dashed line in the forest plots indicate $y = 0$. (**I**) Comparison of the CRAFTseq effect sizes in activated naive B cells between surface protein modality (x-axis) and mRNA modality (y-axis). Dashed lines indicate $x = y$. For the calculation of the Pearson's correlation, we did not include CD40 because the CD40 spQTL signal is driven by *cis*-spQTL of the rs1883832 (CD154 is not evaluated at mRNA modality). (**J,K**) Boxplots showing mRNA (left) and surface protein (right) expression levels of FCER2 (J) and CD27 (K) in activated naive B cells, stratified by rs1883832 genotypes. Expression values are residualized for covariates. The y-axis includes scale breaks. The horizontal dashed line in the forest plots indicate $y = 0$.

GWAS: genome-wide association study; mRNA: messenger RNA; RA: rheumatoid arthritis; spQTL: surface protein quantitative trait loci.

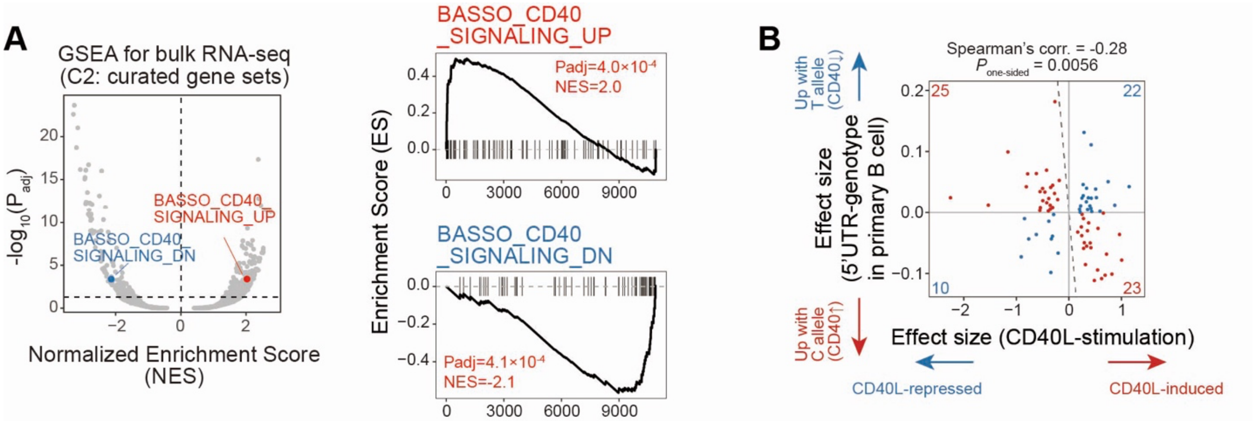


**Fig. S13. Bulk RNA-seq for Daudi cells stimulated with CD40L.** (**A**) Volcano plots showing the normalized enrichment scores (x-axis) and -log10(P-values) (y-axis) from the GSEA of the CD40L stimulation effects on mRNA expression in Daudi cells (left). GSEA enrichment plots are also shown for pathways related to CD40 signaling pathways (right). The horizontal dashed line indicates the FDR 5% threshold, and the vertical dashed line indicates $x$ = 0. (**B**) Comparison of effect sizes of CD40L stimulation from bulk Daudi cell RNA-seq (x-axis) and rs1883832 from primary B cell CRAFTseq (y-axis). Dots are colored based on the consistency of the effect directions (red, opposite direction; blue, same direction). Consistency of the effect direction is tested with a one-sided sign test. Dashed lines indicate $y = -x$. Features satisfying FDR < 0.05 in Daudi cell bulk RNA-seq experiment are included.

GSEA: gene set enrichment analysis; mRNA: messenger RNA; RNA-seq: RNA sequencing.

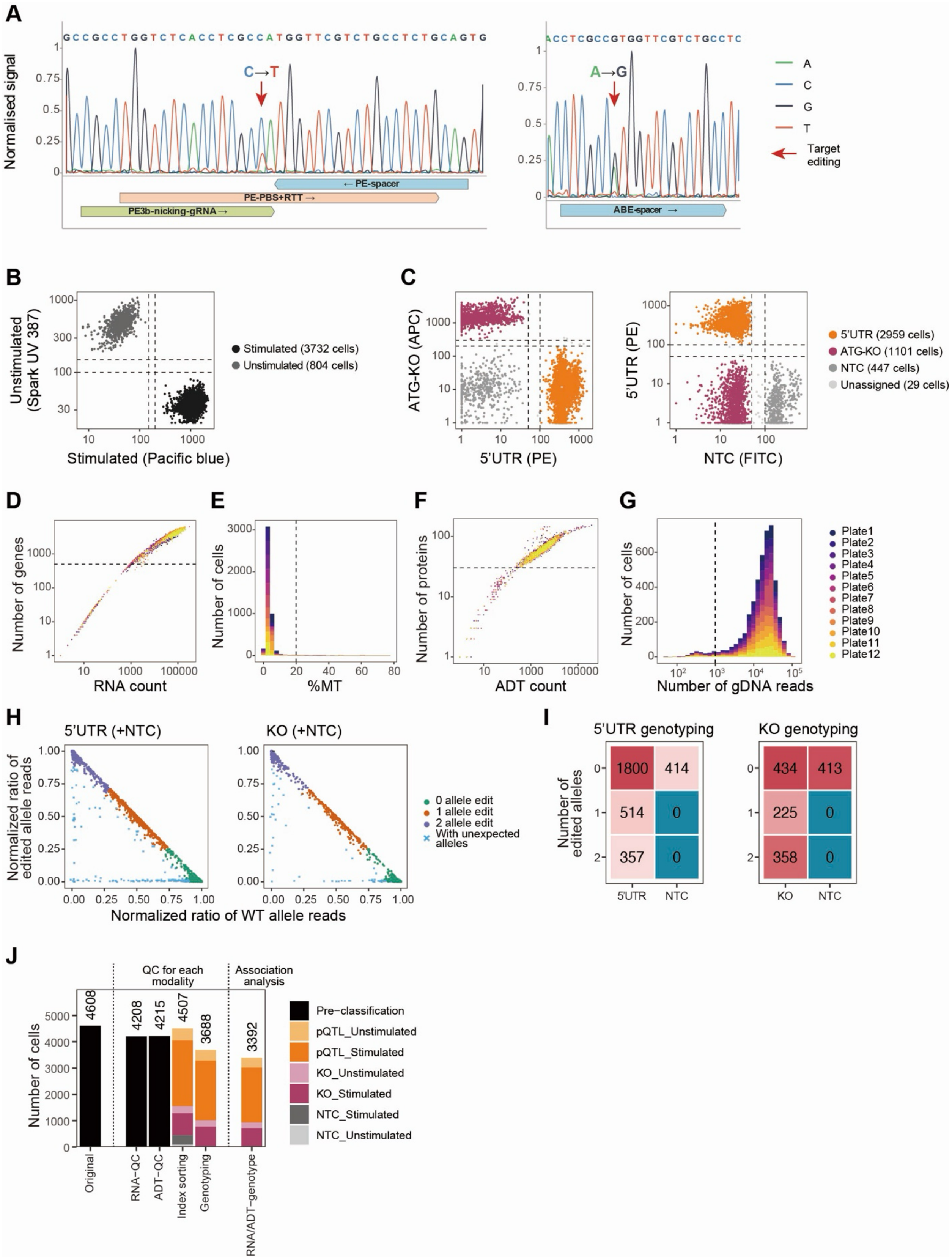


**Fig. S14. Quality controlling for the CRAFTseq with Daudi cells stimulated with CD40L. (A)** Sanger sequencing chromatograms showing the genomic DNA of Daudi cells

after CRISPR editing. The target site is indicated by the red arrow, and the bystander editing site is indicated by the purple arrow. The ribbons indicate the sgRNA or pegRNA sequences, and the corresponding called bases are shown above the chromatograms. (**B**) Indexing flow cytometry data with gating for defining cells stimulated (black) or unstimulated (grey) with CD40L. (**C**) Indexing flow cytometry data with gating for defining cells edited at rs1883832 (orange), ATG start codon (purple), and NTC (grey). (**D**) A scatter plot showing the number of UMI for mRNA (x-axis) and the number of detected genes (y-axis). Dots are colored by plates. A dashed line indicates the number of detected genes = 500. (**E**) A histogram showing the percentage of mitochondrial gene expression for each cell. Bars are colored by plates. A dashed line indicates a percentage of mitochondrial gene expression = 20. (**F**) A scatter plot showing the number of UMIs for protein (x-axis) and the number of detected proteins (y-axis). Dots are colored by plates. A dashed line indicates the number of detected proteins = 30. (**G**) A histogram showing the number of gDNA reads from single cells. Bars are colored by plates. A dashed line indicates a number of gDNA reads = 1000. (**H**) Scatter plots showing the normalized ratio of WT alleles (x-axis) and edited alleles (y-axis) for each variant. Dots are colored by genotyping results. (**I**) Heatmaps showing the genotyping results for each variant. (**I**) A bar plot showing the number of cells at each quality control step. Bars are colored according to the condition.
gDNA: genomic DNA; mRNA: messenger RNA; NTC: non-targeted control; pegRNA: prime editing guide RNA; sgRNA: single guide RNA; UMI: unique molecular identifier; WT: wild-type.

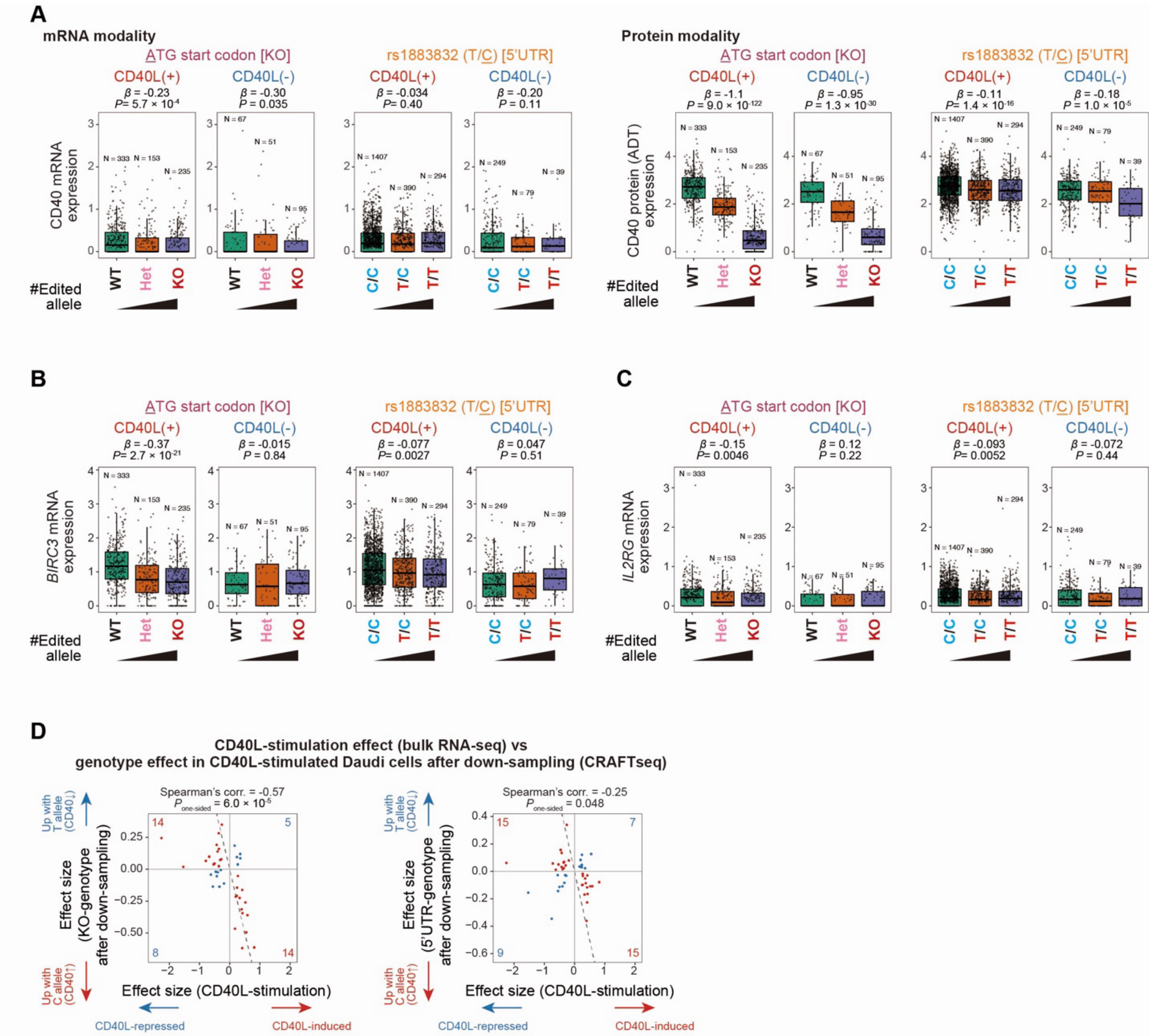


**Fig. S15. CRAFTseq for Daudi cells stimulated with CD40L.** (**A**) Boxplot showing the expression of CD40 mRNA (left) and surface protein (right) across different genotypes of start codon (KO) and rs1883832 (5'UTR), and with or without CD40L stimulation. (**B,C**) Boxplot showing the expression of *BIRC3* (D) and *IL2RG* (E) mRNA across different genotypes of start codon (KO) and rs1883832 (5'UTR), and with or without CD40L stimulation. (**D**) Comparison of the CD40L-stimulation effect in bulk RNA-seq (x-axis) and CRAFTseq effect sizes (y-axis) for ATG-KO (left) and 5'UTR (right). CRAFTseq with CD40L-stimulated Daudi cells is shown after down-sampling to the number of cells from the unstimulated condition. Dots are colored based on the consistency of the effect directions (red, opposite direction; blue, same direction). Consistency of the effect direction is tested with a one-sided sign test. Dashed lines indicate $y = -x$. Features satisfying FDR < 0.05 in Daudi cell bulk RNA-seq experiment (CD40L stimulated vs unstimulated) are included.

KO: knockout; mRNA: messenger RNA; RNA-seq: RNA sequencing; UTR: untranslated region.

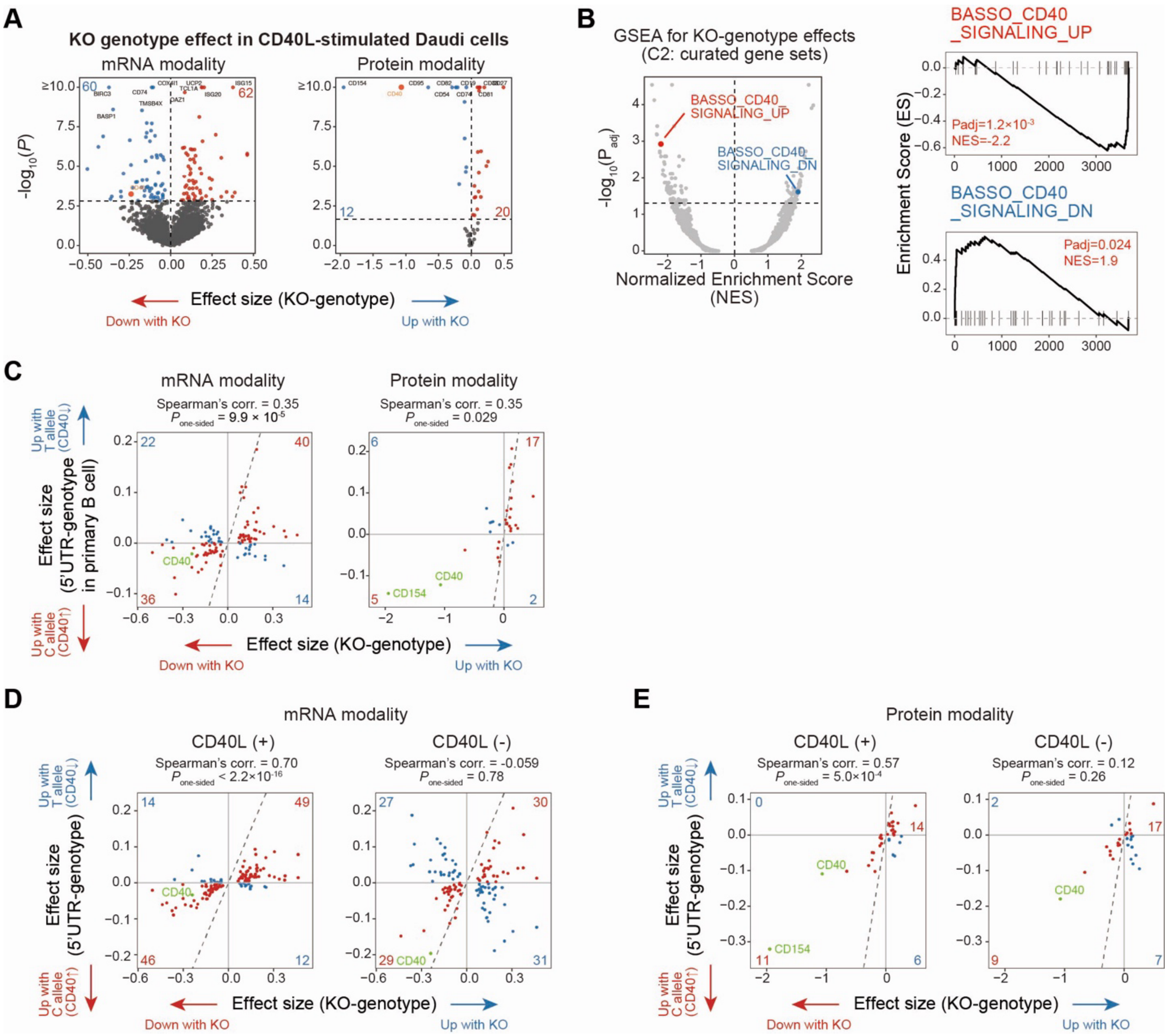


**Fig. S16. CD40L-dependent *trans*-effects of rs1883832 in Daudi cells.** (**A**) Volcano plots showing the effect sizes of the CD40 KO genotype (x-axis) and -log10(P-values) (y-axis) for mRNA (left) and surface protein (right) modalities from CRAFTseq with CD40L-stimulated Daudi cells. The horizontal dashed line indicates the significance threshold (FDR < 0.05), and the vertical dashed line indicates $x = 0$. Significant positive and negative associations are highlighted in red and blue, respectively, with the total number of significant *trans*-associations indicated at the top of each plot. The top 10 associations with the lowest P-values are labeled. (**B**) Volcano plots showing the normalized enrichment scores (x-axis) and -log10(P-values) (y-axis) from the GSEA of the CD40 KO genotype effects on mRNA expression in CD40L-stimulated Daudi cells (left). GSEA enrichment plots are also shown for pathways related to CD40 signaling pathways (right). The horizontal dashed line indicates the FDR 5% threshold, and the vertical dashed line indicates $x = 0$. (**C**) Comparison of effect sizes of CD40 KO genotype from CRAFTseq with CD40L-stimulated Daudi cells (x-axis) and rs1883832 from primary B cell CRAFTseq (y-axis; activated naive B cells) for mRNA (left) and surface protein (right) modalities. Dots are colored based on the consistency of the effect directions (red, same direction; blue,

opposite direction). Consistency of the effect direction is tested with a one-sided sign test. Dashed lines indicate $y = x$. Features satisfying FDR < 0.05 in CRAFTseq with CD40L-stimulated CD40-KO Daudi cells are included. (**D,E**) Comparison of the CD40 KO genotype effect in CRAFTseq with CD40L-stimulated Daudi cells (x-axis) and CRAFTseq effect sizes (y-axis) for 5'UTR in Daudi cells at mRNA (D) and surface protein (E) modalities. CRAFTseq with Daudi cells stimulated (left) or not stimulated (right) with CD40L is shown. Dots are colored based on the consistency of the effect directions (red, same direction; blue, opposite direction). Consistency of the effect direction is tested with a one-sided sign test. Dashed lines indicate $y = x$. Features satisfying FDR < 0.05 in CRAFTseq with CD40L-stimulated CD40 KO Daudi cells are included. For Spearman's correlation calculation (C to E), CD40 and CD154 (only for CRAFTseq) are not included because these two signals are driven by *cis*-pQTL effect of the rs1883832.
FDR: false discovery rate; GSEA: gene set enrichment analysis; KO: knockout; mRNA: messenger RNA; UTR: untranslated region.

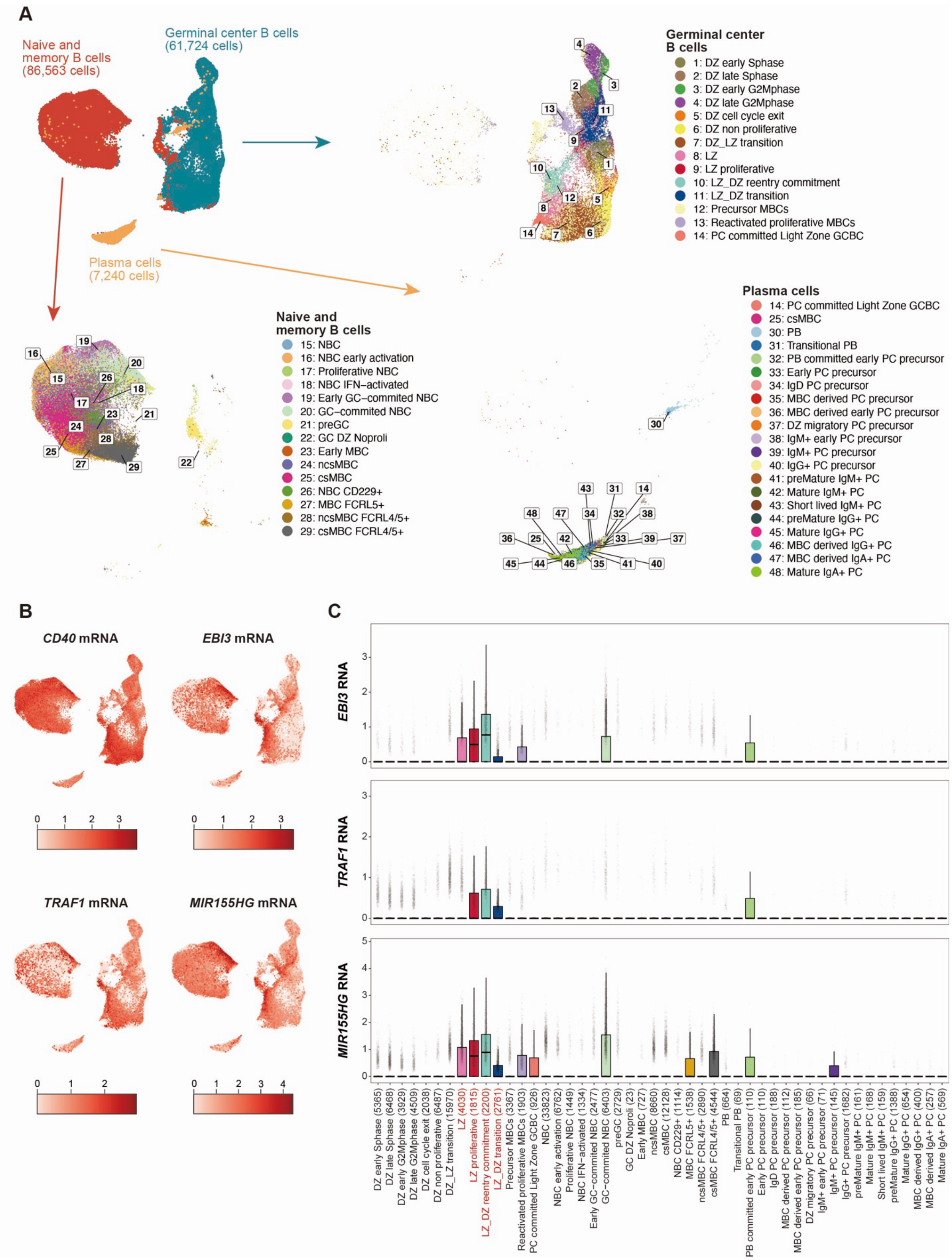
A
Naive and memory B cells (86,563 cells)
Germinal center B cells (61,724 cells)
Plasma cells (7,240 cells)
Germinal center B cells
1: DZ early Sphase
2: DZ late Sphase
3: DZ early G2Mphase
4: DZ late G2Mphase
5: DZ cell cycle exit
6: DZ non proliferative
7: DZ_LZ transition
8: LZ
9: LZ proliferative
10: LZ_DZ reentry commitment
11: LZ_DZ transition
12: Precursor MBCs
13: Reactivated proliferative MBCs
14: PC committed Light Zone GCBC
Naive and memory B cells
15: NBC
16: NBC early activation
17: Proliferative NBC
18: NBC IFN−activated
19: Early GC−commited NBC
20: GC−commited NBC
21: preGC
22: GC DZ Noproli
23: Early MBC
24: ncsMBC
25: csMBC
26: NBC CD229+
27: MBC FCRL5+
28: ncsMBC FCRL4/5+
29: csMBC FCRL4/5+
Plasma cells
14: PC committed Light Zone GCBC
25: csMBC
30: PB
31: Transitional PB
32: PB committed early PC precursor
33: Early PC precursor
34: IgD PC precursor
35: MBC derived PC precursor
36: MBC derived early PC precursor
37: DZ migratory PC precursor
38: IgM+ early PC precursor
39: IgM+ PC precursor
40: IgG+ PC precursor
41: preMature IgM+ PC
42: Mature IgM+ PC
43: Short lived IgM+ PC
44: preMature IgG+ PC
45: Mature IgG+ PC
46: MBC derived IgG+ PC
47: MBC derived IgA+ PC
48: Mature IgA+ PC
B
*CD40* mRNA
*EBI3* mRNA
*TRAF1* mRNA
*MIR155HG* mRNA
C
*EBI3* RNA
*TRAF1* RNA
*MIR155HG* RNA
DZ early Sphase (5365)
DZ late Sphase (6468)
DZ early G2Mphase (3929)
DZ late G2Mphase (4509)
DZ cell cycle exit (2038)
DZ non proliferative (6487)
DZ_LZ transition (15970)
LZ (4030)
LZ proliferative (1815)
LZ_DZ reentry commitment (2200)
LZ_DZ transition (2761)
Precursor MBCs (3367)
Reactivated proliferative MBCs (1903)
PC committed Light Zone GCBC (926)
NBC (33823)
NBC early activation (6762)
Proliferative NBC (1449)
NBC IFN−activated (1334)
Early GC−commited NBC (2477)
GC−commited NBC (6403)
preGC (2729)
GC DZ Noproli (23)
Early MBC (727)
ncsMBC (8660)
csMBC (12128)
NBC CD229+ (1114)
MBC FCRL5+ (1538)
ncsMBC FCRL4/5+ (2890)
csMBC FCRL4/5+ (4544)
PB (664)
Transitional PB (69)
PB committed early PC precursor (110)
Early PC precursor (110)
IgD PC precursor (188)
MBC derived PC precursor (112)
MBC derived early PC precursor (185)
DZ migratory PC precursor (66)
IgM+ early PC precursor (71)
IgM+ PC precursor (145)
IgG+ PC precursor (1682)
preMature IgM+ PC (161)
Mature IgM+ PC (168)
Short lived IgM+ PC (159)
preMature IgG+ PC (1388)
Mature IgG+ PC (654)
MBC derived IgG+ PC (400)
MBC derived IgA+ PC (257)
Mature IgA+ PC (569)

**Fig. S17. Expression of CD40 downstream genes in Tonsil atlas. (A)** B cell annotations of the Tonsil atlas. UMAP colored by B cell annotations in the original study (Massoni-Badosa et al., 2024). **(B)** UMAPs showing the expression of CD40 and activated naive B cell markers (EBI3, TRAF1, and MIR155HG) in the Tonsil atlas. **(C)** Boxplot showing the expression of EBI3, TRAF1, and MIR155HG in the Tonsil atlas across different B cell annotations. Labels for light zone B cells are colored with red.
UMAPs: uniform manifold approximation and projections.

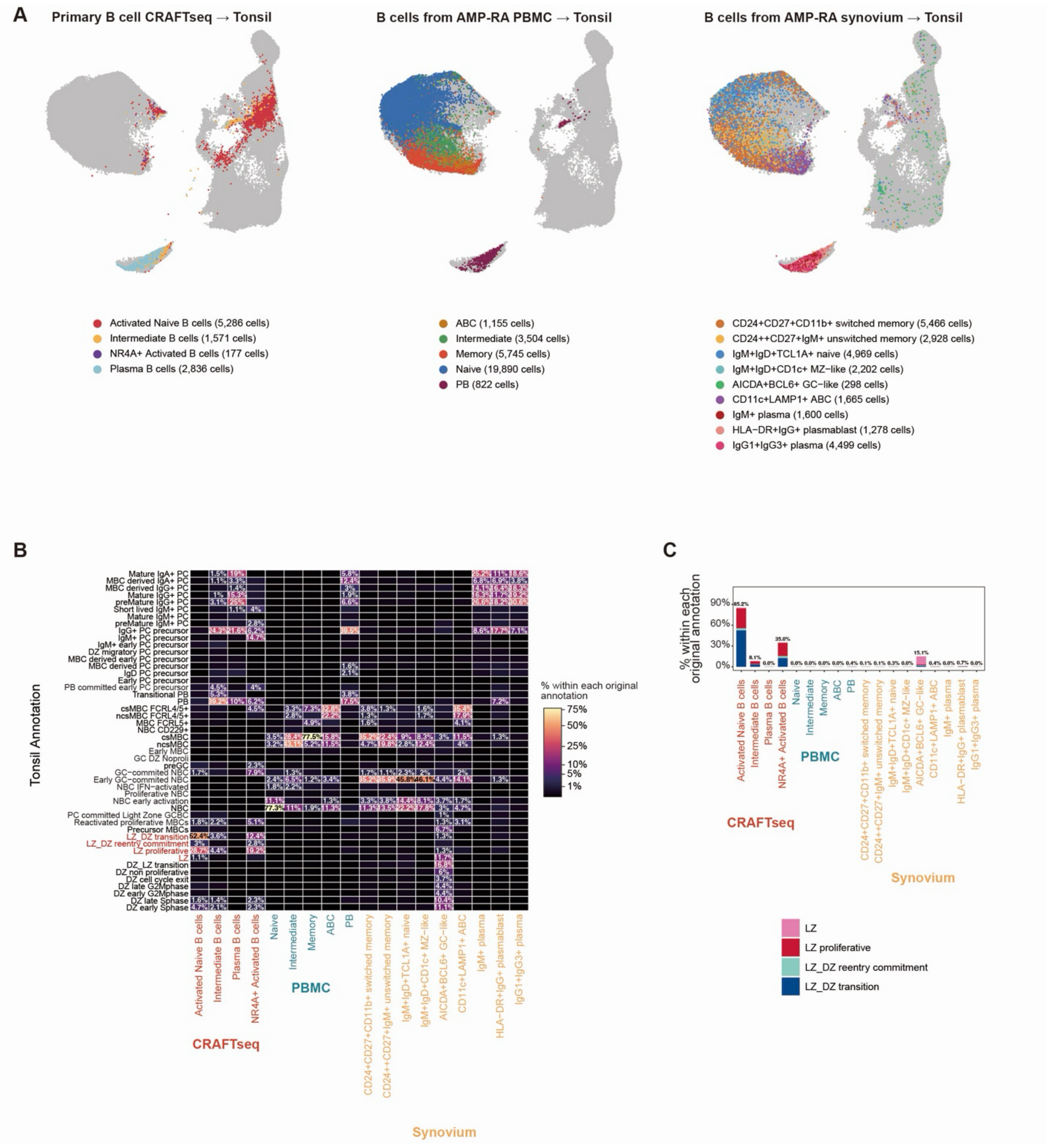


**Fig. S18. Reference mapping of B cell single-cell RNA-seq datasets to the Tonsil atlas. (A)** UMAPs showing the reference mapping of CRAFTseq (left), AMP-RA PBMC (middle), and AMP-RA synovium (right) B. cell datasets to the Tonsil atlas. Dots are colored according to the annotation in each original study for B cell scRNA-seq datasets. **(B)** Heatmap showing the ratio of the tonsil annotation within each original study annotation. Percentages for the tonsil annotation are shown if they exceed 1%. **(C)** Bar plots showing the ratio of the tonsil annotation within each original study annotation.

PBMC: peripheral blood mononuclear cell; scRNA-seq: single-cell RNA sequencing; UMAPs: uniform manifold approximation and projections.

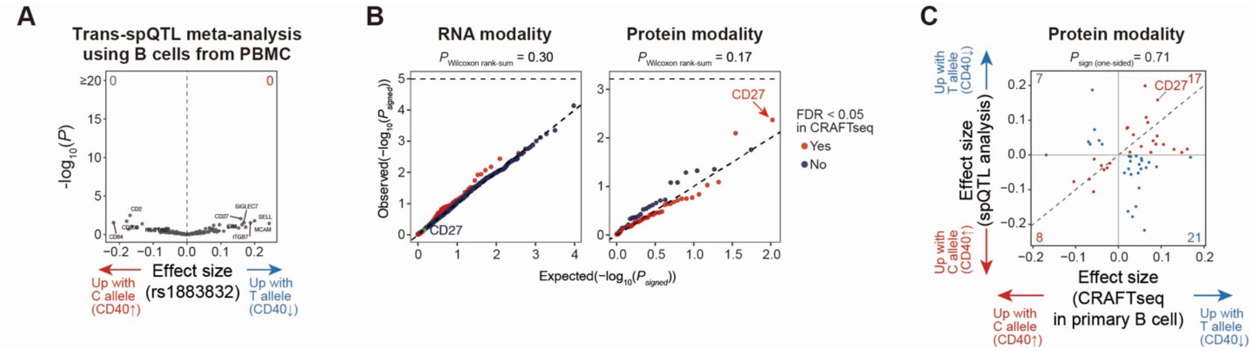


**Fig. S19. Discordance between *trans*-effects identified from *trans*-QTL meta-analysis and CRAFTseq.** (**A**) A volcano plot showing the effect size of rs1883832 (x-axis) and -log10(P-values) (y-axis) for all tested proteins in the trans-spQTL meta-analysis using B cells from CITE-seq data. The vertical dashed line indicates x = 0. The top 10 associations with the lowest P-values are labeled. (**B**) QQ plot of one-tailed P-values for B cell *trans*-QTL meta-analysis for mRNA (left) and surface protein (right). The one-sided P-values are calculated according to the effect directions in the primary B cell CRAFTseq dataset (activated naive B cells). Dots are colored based on whether the genes satisfied FDR < 0.05 in CRAFTseq. The diagonal dashed line indicates $y = x$, and the horizontal dashed line indicates Bonferroni-corrected significance threshold ($\alpha = 0.05$). (**C**) Comparison of effect sizes of rs1883832 from primary B cell CRAFTseq (x-axis; activated naive B cells) and *trans*-QTL meta-analysis (y-axis) for surface protein. Dots are colored based on the consistency of the effect directions (red, same direction; blue, opposite direction). Consistency of the effect direction is tested with a one-sided sign test. Dashed lines indicate $x = y$. Features satisfying FDR < 0.05 in CRAFTseq are included. Note that CD40 and CD154 (only for CRAFTseq) are not included in this analysis because these two signals are driven by *cis*-pQTL effect of the rs1883832.
FDR: false discovery rate; mRNA: messenger RNA; QQ plot: quantile–quantile plot; QTL: quantitative trait loci.

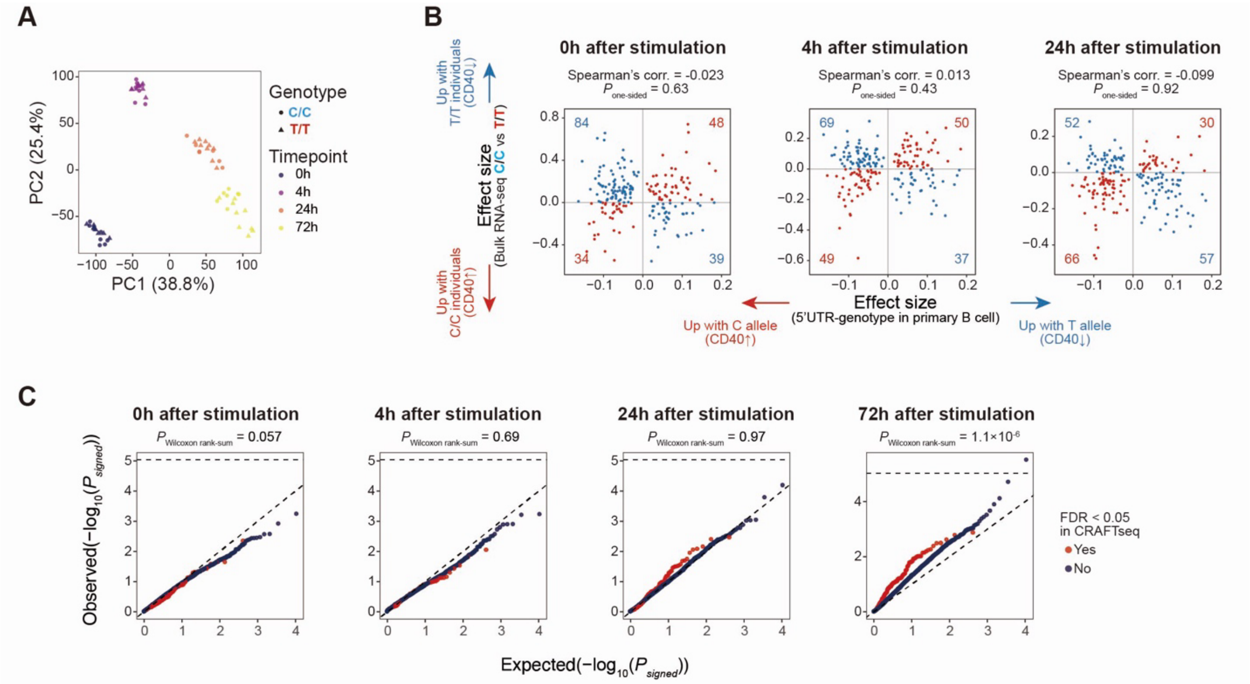


**Fig. S20. Consistency between bulk eQTL analysis and CRAFTseq of cultured primary B cells.** (**A**) Principal component analysis for genes used for eQTL mapping. PC1 and PC2 scores for the 64 samples (16 individuals × 4 time points) are shown, colored by time point. Shape of dots indicate genotypes. (**B**) Comparison of effect sizes of rs1883832 from primary B cell CRAFTseq (x-axis; activated naive B cells) and bulk RNA-seq (y-axis) at 0 h (left), 4 h (middle), and 24 h (right) post-stimulation. Dots are colored based on the consistency of the effect directions (red, same direction; blue, opposite direction). (**C**) QQ plot of one-tailed P-values for *trans*-eQTL analysis from bulk B cell RNA-seq at 0, 4, 24, and 72 h post-stimulation. One-tailed P-values are calculated based on the effect directions observed in the primary B cell CRAFTseq dataset (activated naive B cells). Dots are colored based on whether the genes satisfied FDR < 0.05 in CRAFTseq.

eQTL: expression quantitative trait loci; FDR: false discovery rate; PC: principal component; QQ plot: quantile–quantile plot; RNA-seq: RNA sequencing.

**Contents of the supplementary tables**

Table S1. Number of samples and cells used for QTL analysis (separate file)

Table S2. Significant spQTL signals (separate file)

Table S3. Significant eQTL signals (separate file)

Table S4. PheWAS and colocalization analysis between spQTL and GWAS (separate file)

Table S5. CD40 Surface protein and mRNA expression variance explained by rs1883832 (separate file)

Table S6. Number of cells and features inCRAFTseq experiments (separate file)

Table S7. CRAFTseq analysis of candidate CD40 spQTL causal variants in Daudi cells (separate file)

Table S8. CRAFTseq analysis of rs1883832 in primary B cells (separate file)

Table S9. Differentially expressed gene analysis in CD40L-stimulated Daudi cells (separate file)

Table S10. CRAFTseq analysis of Daudi cells stimulated with CD40L (separate file)

Table S11. Differentially expressed gene analysis in CD40L-stimulated B cells comparing rs1883832 TT and CC genotypes (separate file)

Table S12. Mixed-effects modeling of associations of single cell analysis for B cell and PBMC (separate file)

Table S13. Association between rs1883832 and B cell related traits in Orrù et al (separate file)

Table S14. sgRNA sequences for base-editing (separate file)

Table S15. pegRNA and sgRNA sequences for base-editing (separate file)

Table S16. Antibodies used for CRAFTseq (separate file)

Table S17. CD40 locus-specific primer sequences for CRAFTseq (separate file)

Table S18. CRAFTseq common primers (separate file)

Table S19. CRAFTseq capture primers (separate file)

Table S20. CRAFTseq oligo-dT primers (separate file)

Table S21. CRAFTseq P7 primers (separate file)
